# Effective Polarization in Potassium Channel Simulations: Ion Conductance, Occupancy, Voltage Response, and Selectivity

**DOI:** 10.1101/2024.10.07.616972

**Authors:** Chenggong Hui, Reinier de Vries, Wojciech Kopec, Bert L. de Groot

## Abstract

Potassium (K^+^) channels are widely distributed in many types of organisms. They combine high efficiency (∼100 pS) and K^+^/Na^+^ selectivity by a conserved selectivity filter (SF). Molecular Dynamics (MD) simulations can provide detailed, atomistic mechanisms of this sophisticated ion permeation. However, currently there are clear inconsistencies between computational predictions and experimental results. Firstly, the ion occupancy of the SF in simulations is lower than expected (∼2.5 in MD compared to ∼4 in X-ray crystallography). Secondly, in many reported MD simulations of K^+^ channels, K^+^ conductance is typically an order of magnitude lower than experimental values. This discrepancy is in part because the force fields used in MD simulations of potassium channels do not account for polarization. One of the proposed solutions is the Electronic Continuum Correction (ECC), a force field modification that scales down formal charges, to introduce the polarization in a mean-field way. When the ECC is used in conjunction with the Charmm36m force field, the simulated K^+^ conductance increases 13-fold. Following the analysis of ion occupancy states using Hamiltonian Replica Exchange (HRE) simulations, we propose a new parameter set for Amber14sb, that also leads to a similar increase in conductance. These two force fields are then used to compute, for the first time, the full current-voltage (I-V) curves from MD simulations, approaching quantitative agreement with experiments at all voltages. In general, the ECC-enabled simulations are in excellent agreement with experiment, in terms of ion occupancy, conductance, current-voltage response, and K^+^/Na^+^ selectivity.

**Significance Statement:** Potassium (K^+^) channels are essential membrane proteins that facilitate the selective conduction of K^+^ ions while excluding Na^+^ ions. Incorporation of electronic polarization effects through the Electronic Continuum Correction (ECC) method enables molecular dynamics simulations to reproduce rapid K^+^ permeation under physiological voltage conditions. In this study, we demonstrate that employing ECC approximated polarization in molecular dynamics simulations allows for the prediction of multiple channel properties: conductance, ion occupancy, voltage response, and selectivity, with unprecedented accuracy. Furthermore, our simulations provide atomistic insights into the asymmetrical current-voltage (I-V) relationship observed in the MthK channel.

## 1. Introduction

Potassium channels play an important role in many cell types. They set the resting potential in non-excitable cells or reset the potential in depolarization in excitable cells. Potassium channels are known for their efficient and selective permeation of potassium ions. For instance, the NaK2K channel conducts Na^+^ ions at ∼10 pS in NaCl solution while conducting K^+^ ions at 120 pS in KCl solution.^1^ All potassium channels share a conserved selectivity filter (SF), which forms 4 adjacent ion binding sites made of backbone carbonyl oxygens (sites S1-S3), plus typically 4 threonine hydroxyl oxygens at the S4 site. When the high-resolution crystal structure of KcsA was first published^2^, the electron density in the 4 binding sites was initially interpreted as two K^+^ ions occupying two non-adjacent binding sites and two water molecules placed in between the ions. This interpretation suggests that the four peaks in the electron density represent an overlap of two distinct states: K^+^-water-K^+^-water and water-K^+^-water-K^+^.^3^ Consequently, the hypothesis of a water-mediated “soft knock-on” permeation mechanism was coined. More recent anomalous X-ray diffraction studies in NaK2K and KcsA channels suggested that all of the 4 sites are fully occupied by K^+^ ions,^4–6^ which supports an alternative hypothesis of direct knock-on mechanism, observed in numerous Molecular Dynamics (MD) studies,^7–10^ that does not include any co-permeation of water. A more detailed discussion can be found in our recent review.^11^

Despite the debated permeation mechanism, low conductance and low K^+^ occupancy of the binding sites were consistently observed in MD simulations of potassium channels displaying both soft and direct knock-on mechanisms, although direct knock-on typically results in higher conductance^7,12^. In our previous work, direct knock-on was observed during continuous permeation under membrane voltage, with the SF typically containing 2 or 3 ions.^7^ Specifically, spontaneous permeation under voltage has been observed on MthK, KcsA, NaK2K, BK, Kv1.2, TREK2, and TRAAK,^8,13,14^ but the conductance was generally one order of magnitude lower than the experimental values. Ion occupancy in the SF, permeation mechanism, and conductance in MD simulations all eventually result from a subtle balance of various interactions between ions, protein, and water. Even tiny changes in the interaction parameters between them might change the kinetics, thermodynamics, and mechanism of ion permeation.

One often discussed cause for the observed shortcomings in current simulations of K^+^ channels is the lack of explicit electronic polarization in classical force fields.^15^ In a classical, fixed-charged force field (e.g. Amber, Charmm, OPLS-AA), atoms are modeled as spheres (particles) with assigned partial charges, as an approximation of positively charged nuclei with clouds of electrons around them. In MD simulations, electrostatic interactions are computed between these particles by applying the Coulomb law and using the dielectric constant of vacuum. In reality however, the electrostatic interactions between the nuclei will be (differently) screened by the instantaneous distribution of the electrons, that will be in turn affected by the position of the nuclei (i.e. electronic polarization), resulting in the relative permittivity >1. Therefore, for the more accurate modeling of electrostatic interactions, two general solutions have been proposed. In the first one, force fields with explicit electronic polarization are developed, whereas in the second one, the so-called Electronic Continuum Correction (ECC) is applied to existing, classical force fields. Ren and coworkers have tested both solutions in MD simulations of a K^+^ channel^16^, focusing on the different K^+^ occupancy states of the SF. They found that when the ECC is applied, Charmm36m behaves similarly to the AMOEBA polarizable force field, and both show the SF being fully occupied with four K^+^ ions to be energetically the most stable configuration. In practice, the ECC is achieved by rescaling the formal partial charges of ionized groups (such as side chains of arginine, lysine, glutamine, and aspartate amino acids) and ions, which is mathematically the same as changing the dielectric constant in computing electrostatic interactions via Coulomb law.^17,18^ This approach has been recently applied to improve force fields for ions, proteins, and lipids.^19–23^ In general, the argument is made that the rescaling factor should be ∼sqrt(1/2) ≈ 0.7, because the electronic screening factor (high-frequency dielectric constant) is about 2 for most organic media.^24^ However, there are at least two reasons why directly applying this number might not be optimal. First, the exact dielectric constant of the approximated medium between MD particles is not known and might be system-dependent. Our main interest here is the magnitude of channel conductance and corresponding free energy difference between relevant SF occupancies. Second, most of the force field parameters were developed without ECC in mind, and might compensate for the mean polarization to some extent. Therefore, instead of focusing on a single scaling factor, in this work, we combine the ECC approach with two common fixed-charge force fields: Charmm36m and Amber14sb, at various scaling factors (1.0-0.65), to study ion permeation in three different potassium channels. We carefully developed new Lennard-Jones (LJ) parameters, describing van der Waals (vdW) interactions, at each scaling factor, so that the ion radius and solvation free energy are consistent with the original force field. We observe that the ECC dramatically increases the simulated conductance in Charmm36m, to a level that quantitatively agrees with the experimental single-channel conductance. In contrast, the ECC has only a minor effect with the Amber14sb force field. These discrepancies are rationalized using Hamiltonian Replica Exchange (HRE) simulations, providing insights into the effect of the ECC on the SF ion occupancy, and resulting in adjusted SF partial charges for Amber14sb, which consequently predicts conductance close to the experiments as well. We further apply these force fields in simulations under physiological voltage <-150, +150 mV>, recording, for the first time, full I-V curves, compatible with experiments. The K^+^/Na^+^ selectivity is preserved, indicating the necessity of adjusting LJ parameters of ions in ECC. Finally, applying the recently introduced permeation cycle analysis, we provide cues into the physical basis of inward rectification in the MthK channel.

## 2. Results and Discussion

### 2.1. Low conductance and SF occupancy in MD with Charmm36m and Amber14sb force fields

To obtain a comprehensive view on ion permeation in K^+^ channels, we simulated three different channels, namely NaK2K, TRAAK, and MthK. Whereas the archaeal MthK channel shares the SF sequence (TVGYGD) with the NaK-derived NaK2K channel, the SF sequence of TRAAK is more varied: being a dimer, TRAAK has two SF strands (called SF1) with a TIGYGN sequence, and two with TVGFGD (SF2). The inclusion of TRAAK simulations allows us to generalize our findings beyond the most common SF sequence. The channels were simulated using both the Charmm36m and Amber14sb force fields. When compared with experiments, a clear discrepancy in quantities describing ion permeation, such as conductance and SF occupancy is seen, as previously reported by us and others.^7–9,13^ Generally speaking, single channel conductance (Fig. 1a) is underestimated by an order of magnitude in all channels and force fields tested. The instantaneous occupancy of the SF binding sites during permeation (Fig. 1b) predominantly shows (despite differences between force fields, Table S1) an alternation between 3-ion and 2-ion (counting ions in S1 to S4) SF occupancy states. Although this quantity is not readily available from experiments, and the SF occupancy during permeation is debated (see Introduction), we note here that the 3-ion and 2-ion states underestimate the possible number of ions in the SF, as compared to anomalous diffraction data (full SF occupancy in S1-S4 at 100 K and 3.44 total occupancy at 292 K).^4,25^ These results clearly show the need for improved force fields for MD simulations of K^+^ channels. Efforts have been made in this direction. For example a special exception interaction (NBFIX) was developed to weaken the interaction between carbonyl oxygens and K^+^ ions,^26^ but reproducing the conductance was found to be challenging.^27^ We reason that single channel potassium conductance is the central quantity describing ion permeation, which can be directly compared between simulations and experiments. Therefore, an improved force field should predict the single channel conductance close to the experimental value, across the same voltage range as used in electrophysiology (-150 ∼ 150 mV), and ideally also in the physiological range (resting potential: -10 ∼ -100 mV, action potential: ∼+100 mV).^28,29^ We further reason that such a force field would likely better describe any secondary observables characterizing ion permeation (i.e. SF occupancy states, rate limit steps, SF geometries etc.) as well.

**Figure 1.**
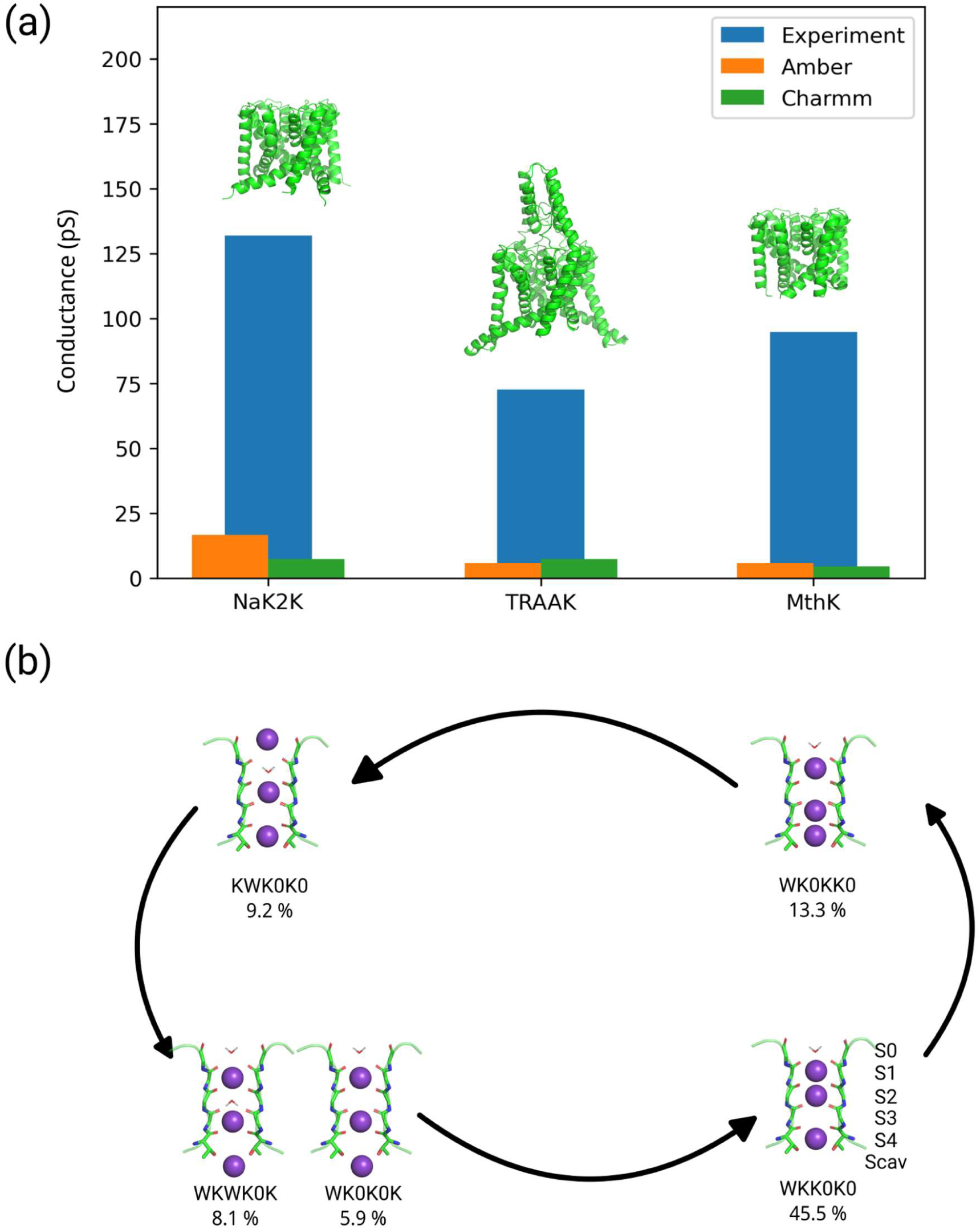
Main features of K^+^ permeation through K^+^ channels in MD simulations with traditional force fields. **(a)** Comparison of experimental conductances^1,10,30^ and MD simulated conductances in 3 channels (NaK2K, TRAAK, MthK), with 2 force fields (Charmm36m and Amber14sb), at +100 mV. **(b)** A schematic mechanism of ion permeation in NaK2K in MD simulations using Charmm36m at +100 mV. Trajectories were discretized based on the number of K^+^ ions in positions S0-SCav. The SF occupancy is represented using a 6-letter code showing the occupancy at binding sites S0 to SCav: ‘K’ for a K^+^ ion, ‘W’ for water, and ‘0’ for an unoccupied site. Populations of SF states are expressed as percentages, with WKK0K0, WK0KK0, KWK0K0, WKWK0K, and WK0K0K being the only states exceeding a 5% population.

In this contribution, we show how the ECC-introduced polarization in Charmm36m and Amber14sb force fields improves simulated single channel conductance in all channels tested. Interestingly, the application of charge scaling simultaneously results in SFs occupied by more K^+^ ions (3-ion and 4-ion states) than in non-scaled force fields, agreeing well with experimental data, and further highlighting the role of direct ion-ion contact in rapid K^+^ permeation through K^+^ channels.

### 2.2. ECC-Charmm36m improves the conductance, but ECC-Amber does not

We performed simulations with different scaling factors, starting at the nominal charge of +1.0, with a step of 0.05, down to +0.65, in Charmm36m and Amber14sb. Similarly to simulations with unscaled charges, we initially used an applied voltage of +150 mV to drive ion permeation. The conductance at different levels of ECC (scaling factors) in the NaK2K channel is shown in Fig. 2a-c (see Fig. S4 and S5 for TRAAK and MthK channels, respectively). We first focus on simulations using Charmm36m (Fig. 2a, d, g). Compared to the standard parameters (scaling factor of 1.00), the conductance decreases at +0.9 charge, but then significantly increases around ∼+0.75. At the scaling factor of 0.75, the conductance is 122.4 ± 10.1 pS, which is a 13-fold increase compared to standard Charmm36m (9.2 ± 1.7 pS), and in apparent excellent agreement with the experimental single channel conductance of ∼120 pS.^1^ The equilibrated populations of the SF states show major changes with the applied scaling factor. Compared to standard Charmm36m, 4- ion states become gradually more populated with stronger scaling (Fig. 2d, g): the WKKKK0 state is dominant (79%) with 0.7 scaling, in stark contrast to unscaled Charmm36m (WKK0K0 (37%), WK0KK0 (20%), WK0K0K (20%)). The increase of ion occupancy is consistent with the previously reported ECC-Charmm36m simulation in KcsA.^16^ A similar effect of increased ion occupancy with charge scaling is observed in Amber14sb (Fig. 2e,h), however there is no effect on conductance, as it remains low (∼20 pS) at all scaling factors tested, also in the other two channels (Fig. S4 and S5).

**Figure 2.**
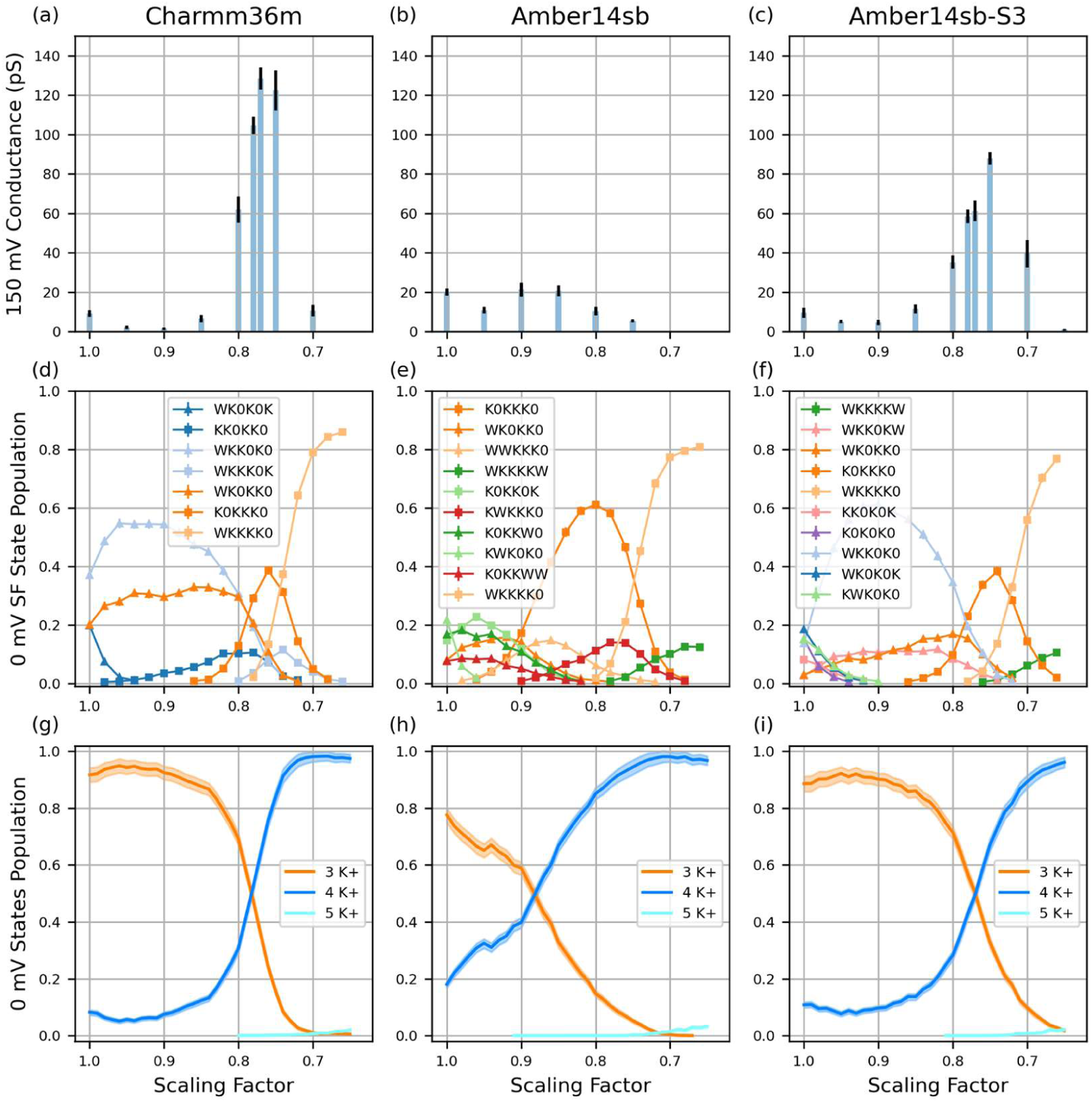
Charge scaling has a major impact on ion permeation characteristics in the NaK2K channel. (**a-c**) Average conductances at different scaling factors in MD simulations using Charmm36m, Amber14sb, and Amber14sb-S3 (see text for explanation) force fields under +150 mV in NaK2K. The error bar is the standard error of the mean (N=10). **(d-f)** Equilibrated populations of the SF states recorded in HRE in NaK2K, using a 6-letter code showing the occupancy at binding sites S0 to Scav: ‘K’ for a K^+^ ion, ‘W’ for water, and ‘0’ for an unoccupied site. For clarity, only the even replicas are plotted (replica 0, 2, 4 etc at scale 1.00, 0.98, 0.96 etc). The odd replicas are not shown (replica 1, 3, 5 etc at scale 0.99, 0.97, 0.95 etc). **(g-i)** Ion occupation state distribution from HRE simulation, the states with the same number of ions in the SF are summed together. In (d-i) The error bar is the 95% confidence interval when bootstrapping frames.

### 2.3. Redistributing the charge in the SF in Amber14sb increases the conductance

Prompted by the conductance insensitivity to charge scaling in Amber14sb, we decided to investigate this issue in detail. By comparing simulations carried out with Charmm36m and Amber14sb we noticed two important (and possibly related) differences: 1. In Charmm36m, the carbonyl oxygen has a partial charge of -0.51, whereas it is -0.5679 in Amber14sb, potentially resulting in stronger binding of a K^+^ ion to the SF, and 2. For the <0.85, 0.75> range of charge scaling in Amber14sb, the K0KKK0 state is the dominant one, and, for scalings < 0.75, WKKKK0 dominates (Fig. 2e). These observations suggest that K^+^ ions in Amber14sb are potentially too strongly bound to the SF, especially to its lower part, which might slow down permeation. To test this idea, we reduced the negative charge on the carbonyl oxygen atoms at S3 to different values (-0.55, -0.53, -0.51), and evenly distributed the remaining charge on the other atoms in the SF (see Fig S1 for the illustration of oxygen charge in the SF). -0.53 gave the SF state population similar to Charmm36m (see Fig. 2 and Fig. S2 for SF state population). We further tested the -0.53 charge modification and call it Amber14sb-S3 from now on.

With this modified force field at hand, we repeated all the simulations for all three channels (Fig. 2c, f, i, Fig. S4, Fig. S5). Interestingly, the state populations (Fig. 2f and i) are now more similar to ECC-Charmm36m. When compared to the original Amber14sb, the major difference is that the population of K0KKK0 which dominated the distribution at 0.8 (Fig. 2e) is now significantly reduced (Fig. 2f). Importantly, these effects resulted in a major increase in conductance, akin to ECC-Charmm36m (Fig. 2c, maximal conductance at the scaling factor of 0.75: 88.0 ± 3.2 pS). We observe essentially the same behavior in simulations of TRAAK and MthK channels (Fig. S4 and S5).

To obtain clearer mechanistic insights into the factors governing rapid ion permeation at around 0.80-0.75 scaling in both Charmm36m and Amber14sb-S3, we looked again at the populations of SF states, but this time combining states together with the same number of K^+^ ions bound to the SF (Fig. 2g-i, Fig. S4, Fig. S5). We find that in high conductance regions, there is a similar population for 3-ion and 4-ion population states (crossing of blue and orange lines), suggesting a smooth (barrierless) free energy landscape between microscopic states (SF occupancies). We hence hypothesize that a balanced (low barrier) free energy surface between multiple 3- and 4-ion states is a prerequisite for efficient, rapid ion permeation through the SF of K^+^ channels. Accordingly, if certain states dominate the SF state distribution the permeation cycle would be slowed down by the barrier to leaving this state. For example, in Charmm36m at 0.9, the SF is mostly populated by a single WKK0K0 state, whereas in Amber14sb at 0.8, the SF is mostly populated by K0KKK0. Those extra stable states might contribute to the underestimation of conductance in MD simulations.

Our proof-of-concept Amber14sb-S3 modification opens up a new degree of freedom for force field development in K^+^ channel simulations. The default backbone parameters for the SF in Amber14sb do not yield reasonable conductance, instead they result in an SF being stuck at a certain SF state (e.g. K0KKK0 at 0.8 scale). By a slight modification of a partial charge on the backbone carbonyl oxygen (from -0.5679 e to -0.53 e) at the S3 binding site, we adjusted the relative free energies of SF states resulting in an efficient ion permeation. We do not suggest our applied parameter changes as a general force field patch, but a special parameter set for K^+^ channel SF.

Comparing the conductance-scaling relationship for all three channels (Fig. 2a-c, Fig. S4a-c, Fig. S5a-c), the maximal conductance is observed at remarkably similar scaling factors. In ECC- Charmm36, the conductance for TRAAK and NaK2K reached their maximum at 0.78 and 0.77, whereas for MthK at 0.75. In ECC-Amber14sb-S3, 0.78 for TRAAK, 0.75 for NaK2K and MthK. Similarly, the crossing point of 3-ion and 4-ion state populations appears in high conductance regions for all three channels, following the same order: TRAAK 0.82, NaK2K 0.78, MthK 0.77 (Charmm36m, Fig. 2g, Fig. S4, Fig. S5), TRAAK 0.79, NaK2K 0.77, MthK 0.77 (Amber14sb-S3, Fig. 2i, Fig. S4, Fig. S5). These results agree well with our hypothesis that the presence of multiple 3- and 4-ion states at similar (balanced) free energy levels is a prerequisite for high conductance in K^+^ channels. And also agree with the room temperature anomalous scattering occupancy result.^25^

### 2.4. The ECC force fields predict I-V curves in agreement with the experiment

Encouraged by the unprecedented increase of the simulated single channel conductance at a single voltage of +150 mV when using ECC-Charmm36 and ECC-Amber14sb-S3, we decided to test these force fields in the whole experimental voltage range. As a result, we obtained, for the first time, the full current-voltage (I-V) curves of K^+^ channels derived purely from MD simulations, for experimentally and physiologically relevant voltage ranges (Fig. 3). The ECC correction improves conductance at all voltages tested, bringing simulated currents with the ∼0.80-0.75 scaling factors very close to the experimental currents (black traces) for all three channels, and in both force fields. This effect is particularly pronounced for the experimentally relevant voltage range (<-150 mV, +150 mV>), where both outward and inward currents with unscaled charges are very low (blue traces). Notably, in all conditions tested, the rapid ion permeation occurs through some variations of the direct knock-on mechanism, that is without any co-permeating water molecules. Generally, the probability of finding water in SF is very low. (K^+^/Water occupancy in SF is listed in Table S3). Similar to our findings in the previous section, the highest conductance at all voltages is enabled by the co-existence of several 3-ion and 4-ion states in the SF with similar populations.

**Figure 3.**
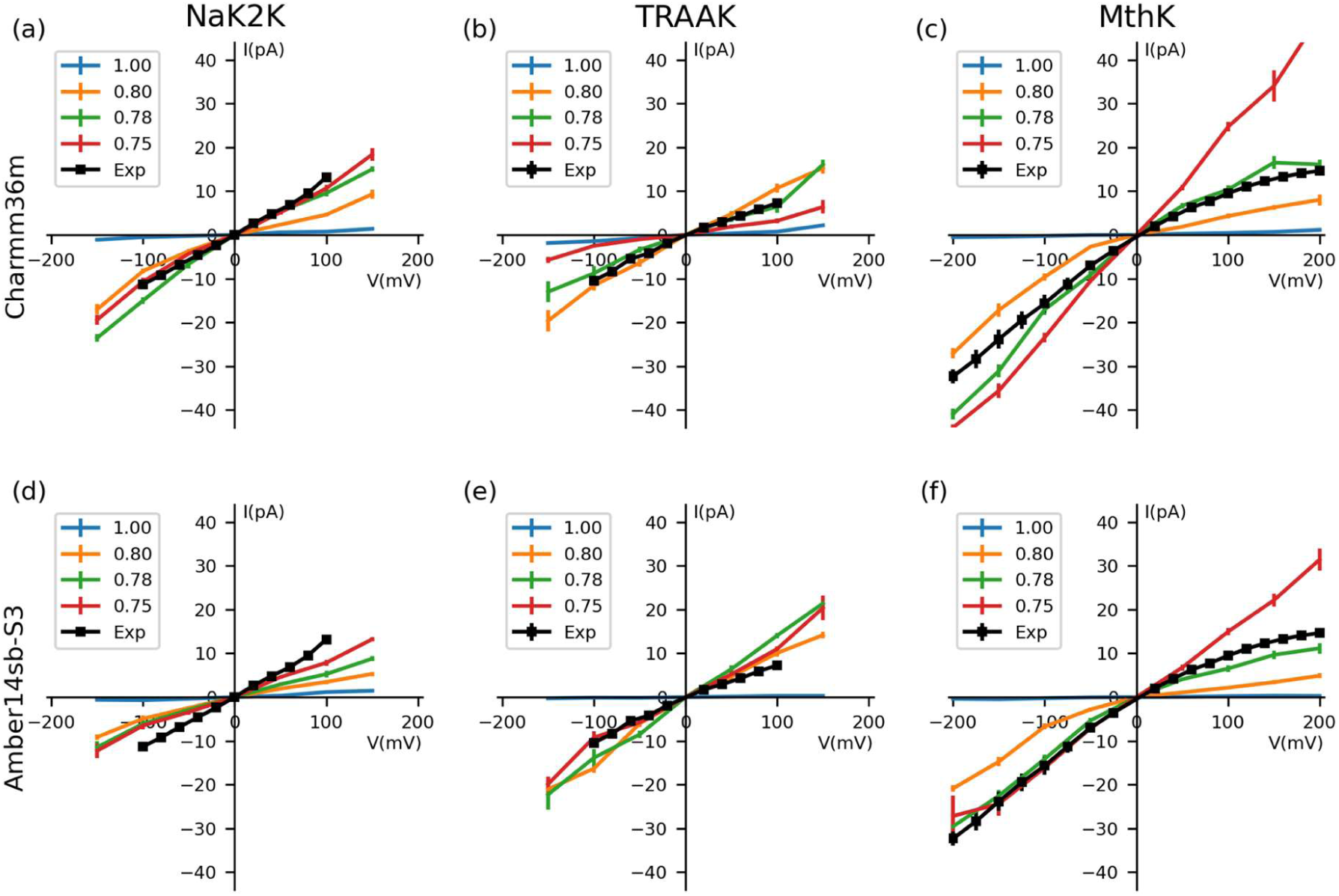
Simulated full current-voltage (I-V) curves for NaK2K, TRAAK, and MthK channels at different scaling factors. **(a-c)** Simulated I-V curves with Charmm36m. **(d-f)** Simulated I-V curve with Amber14sb-S3. MD simulations were performed at ± 50 mV, ± 100 mV, ± 150 mV, ± 200 mV (if the experiment data covered ± 200 mV). The experimental data are shown in black and were provided by the authors of respective publications.^1,10,30^ Simulated data are shown in colors for different scaling factors. The error bar is the standard error of the mean in all cases. The digital data can be found in the SI file.

In some cases, the application of the ECC correction leads to simulated currents that are higher than their experimental counterparts (e.g. MthK in ECC-Charmm36m 0.75 scaling at positive voltages, or TRAAK ECC-Amber14sb-S3 0.80 scaling at all voltages), which, to the best of our knowledge, has never been reported before for any MD simulation. This observation poses a challenge in deciding which scaling factor is the optimal one (for a given force field). In this context, it is worth recalling that the shapes of experimental I-V curves might substantially vary between K^+^ channels. For the channels tested here, both NaK2K and TRAAK display linear I-V curves^1,30^, whereas MthK shows inward rectification^10^. Taking this into account, we notice that ECC-Charmm36m with the 0.78 scaling factor (Fig. 3 (a-c), green traces) is in best agreement with the experimental data, reproducing well shapes of experimental I-V curves, including the inward rectification phenomenon in MthK. A more complex situation is seen in ECC-Amber14sb-S3. Here, the 0.78 scaling factor (Fig. 3 (d-f), green traces) is in almost perfect agreement with experimental data for MthK, and it also predicts the linear I-V curves for NaK2K and TRAAK, but underestimates the current for Nak2K and TRAAK. These examples show that none of the ECC scaling tested here is ideal, and further reparameterization work is needed to obtain a robust set of parameters applicable to MD of any K^+^ channel. The observed differences between studied channels (and also scaling factors) might be rationalized by sequence variations in the selectivity filter and the second layer of residues surrounding the selectivity filter. Indeed, while the SF sequence is highly conserved, the residues around it show a higher degree of variability, likely leading to ion occupancy and conductance changes, due to subtle differences in dynamics and intermolecular interactions. However, these observations remain secondary to the primary effect of the ECC correction, that is the substantial (order of magnitude) increase of the single channel current (conductance), observed in the 0.80-0.75 scaling range, to the level comparable with experimental estimates. Thus our work opens the possibility to use, for the first time, MD simulations of K^+^ channels in a quantitative manner to study a wide range of phenomena, using the same observable (the current-voltage relationship) as in electrophysiological experiments.

Our ECC simulations carried out at voltages outside of the experimental range of <-150 mV, +150 mV> provide additional insights. For example, the scaling factor showing the maximal conductance in ECC-Charmm36m for NaK2K at 200 mV appears to be 0.75, and, in the absence of experimental data at this voltage, it could have been deemed optimal. However, simulations at lower positive voltages show that this is not the case. Therefore, we conclude that in the future MD simulations of ion channels should be performed, whenever possible, at experimentally and/or physiologically relevant voltages. High voltages have been traditionally used to speed up ion permeation in MD simulations of K^+^ channels,^7,10,13,27^ occasionally reaching 500 mV or even 750 mV,^27,31,32^ although the effects of such high voltages on the protein structure and the permeation mechanism itself are not clear.^7^ With the ECC approach, there does not seem to be a need to use such high voltages anymore, since robust currents are achievable at experimental voltages as well.

Finally, we tested two more channels, Kv1.2-2.1 and KcsA E71A with the optimal parameters for ECC-Charmm36m at 0.78 scaling factor. Good agreement with experimental currents at both +150 mV and -150 mV are observed. (Fig. S7)

### 2.5. K^+^/Na^+^ selectivity is sensitive to both the ion partial charge and LJ parameters

All K^+^ channels show exquisite K^+^/Na^+^ selectivity (>10:1 permeability ratio, i.e. the ratio of conductances in pure K^+^ and Na^+^ solutions),^1,33^ therefore this important characteristic should be captured in simulations as well. In this section, we first present thermodynamic properties of our newly developed radius-corrected LJ parameters for ECC ions (All of the simulation results above were using these radius-corrected LJ parameters). Secondly, we compare ECC K^+^ channel simulations with these new parameters versus unmodified LJ parameters, highlighting the importance of compatible ion LJ parameters in ECC force fields.

The straightforward application of the charge scaling procedure in the ECC has a big effect on ion properties in water. Particularly relevant for ion permeation is the radius of an ion, which is typically carefully checked versus experimental data during the development of ion parameters. When scaling of the charge is applied, cations will have effectively larger radii, due to the reduced interactions of a scaled charge on an ion with the surrounding water molecules (Fig. 4a), thermodynamically leading to an underestimation of the solvation free energies (Fig. 4b, see Methods for the derivation of the solvation free energy scaling in the ECC).

**Figure 4.**
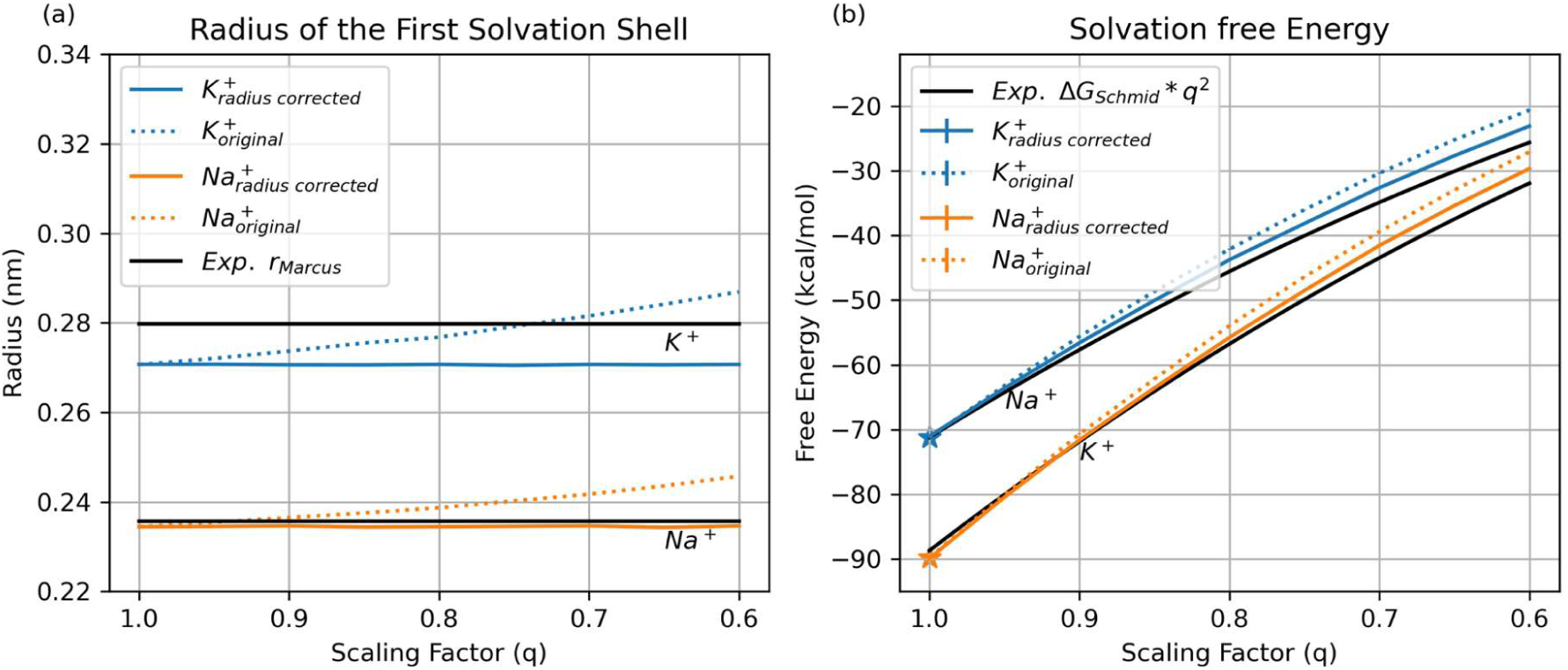
Comparison of the radius corrected LJ parameters for ECC ion in this work (solid line) and the original vdW (dotted line) in Charmm36m. (a) The radius of the first solvation shell for K^+^ and Na^+^ with two sets of LJ parameters, compared to the experimental value.^67^ (b) The solvation free energy for K^+^ and Na^+^ with two sets of vdW parameters. The experimental solvation free energy from Schimdt et al.^68^ was scaled at each scaling factor, following the Born model (see Methods for details). The experiment value was from the respective references. The vdW parameters at 1.0 scale were taken from the original force field. The same plot for Amber14sb is shown in Fig. S10.

To amend this situation, we developed new radius-corrected LJ parameters for ECC ions. This was done in a way that at any given scaling factor, the radius of the first solvation shell reproduces the radius in the original (not scaled) force field. We decided to choose the original force field as the fitting target rather than the experimental value to: 1. facilitate the comparison with the original force field, and 2. reduce the perturbation to the original force field. As shown in Fig. 4a, our new parameters for K^+^ and Na^+^ consistently reproduce the originally parametrized radius. Importantly, also solvation free energies of our radius-corrected ECC ions closely follow experimental values (Fig. 4b).

To test the radius-corrected LJ parameters, we performed MD simulations of the same K^+^ channels with Na^+^ ions at +150 mV at different scaling factors, and compared simulated outward conductance with K^+^ conductance and experimental values (Fig. 5, Fig. S14). The unmodified vdW is only used in Fig. 5 (d-e) and Fig. S14 (d-e) for comparison, it is not recommended to use the unmodified vdW in charge scaling simulations.

**Figure 5.**
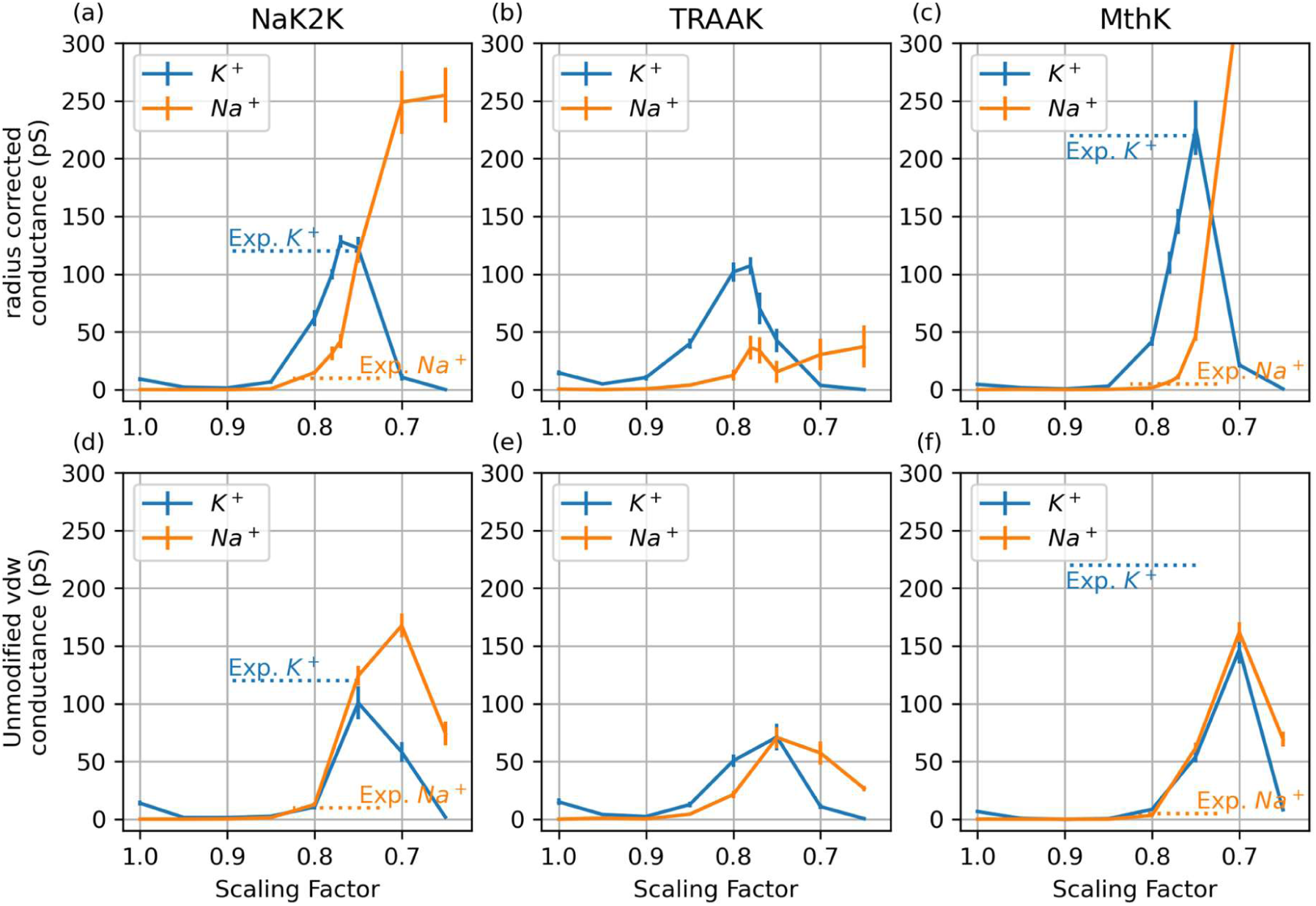
K^+^ and Na^+^ conductance with two treatments of LJ parameters in Charmm36m. **(a-c)** Simulated conductances with our newly developed radius-corrected LJ parameters for ECC ions. **(d-f)** Simulated conductance with unmodified LJ parameters in Charmm36m. All simulations were performed under +150 mV voltage. The analogous plot for Amber14-S3 is in Fig S14. The experimental values were taken from the respective references.^1,33^

With our radius-corrected LJ parameters (Fig. 5 a-c, Fig. S14, Fig. 2, Fig. 3), we observe consistently larger K^+^ conductances than Na^+^ conductances at all scaling factors larger than ∼0.7. Specifically, combined with our I-V curves, we deem the scaling factor of 0.78 optimal for current simulations, i.e. it produces I-V curves that qualitatively match experimental ones, and simultaneously provides robust K^+^/Na^+^ selectivity for all three tested channels (Table 1). In contrast, for a smaller scaling factor (<0.7), all three channels conduct more Na^+^ than K^+^ ions, which indicates incorrect selectivity. This observation highlights the balance of strong interactions governing ion permeation and selectivity in K^+^ channels, leading to the necessity of careful validation of both the scaling factor and LJ parameters of their accurate simulations. We acknowledge that we did not achieve a perfect match for the conductance ratio across all channels (Table 1), but accurately obtaining energy barrier differences to predict the exact ratio may exceed the precision limits of the classical force field. According to the Eyring equation,^34^ a 10:1 ratio in permeation rates at 310K corresponds to approximately the 1.4 kcal/mol difference in the barrier height, and the 2.5:1 ratio corresponds to the 0.6 kcal/mol difference.

**Table 1.** K^+^/ Na^+^ conductance ratio at 0.78 scale, at +150 mV.

|  | NaK2K | TRAAK | MthK |
| --- | --- | --- | --- |
| Charmm36m | 3.3 | 2.5 | 16.1 |
| Amber14sb-S3 | 2.5 | 7.7 | 8.4 |

In contrast, when the original (unmodified) LJ parameters are used, K^+^/Na^+^ selectivity is lost in almost all conditions tested (Fig. 5 d-e, Fig. S14). Our results therefore suggest that the adjustment of LJ parameters, and, consequently, maintaining the correct radii of ions, is necessary to capture their proper permeation and selectivity behavior in K^+^ channels with the ECC correction, and will be likely relevant in other systems showing significant ion selectivity.

### 2.6. Mechanistic Basis of Inward Rectification

As a final insight from our ECC simulations, we decided to take a closer look at simulations displaying inward rectification in the MthK channel (ECC-Charmm36m at 0.78 scaling, Fig. 3c, green curve), that is the larger current at negative voltages than at positive ones. This inward rectification is in excellent agreement with experiment (Fig. 3c, black curve). Importantly, this inward rectification is observed, both experimentally and computationally, for the pore domain only (i.e. without the additional intracellular calcium-binding domains), indicating that it is an inherent property of the K^+^ permeation mechanism in this channel. To study it in more detail, we applied a recently proposed analysis of ion permeation mechanisms using permeation cycles.^9^ We simplified the cycle by lumping together several states when their exchange rate was above a certain cutoff. The comparison of permeation cycles at +150 mV and -150 mV (Fig. 4) reveals that the rate-limiting step at both voltages is the transition between WKKK0K and WKK0KK states (i.e. a K^+^ ion transitioning from S4 to S3 sites during outward permeation, and in the opposite direction during inward permeation; while both S1 and S2 remain occupied by K^+^ ions). However, the rate (inverse of the mean first passage time) of this transition differs by 3-fold for these two voltages (0.318 ns^-1^ for +150 mV and 0.912 ns^-1^ for -150 mV). Our approach allows finding such rate limiting states in MD simulations of K^+^ channels, and thus deciphering the molecular determinants governing the existence of such states, and rectification properties in different K^+^ channels in general.

## 3. Conclusion

In this work, we introduced and tested a range of parameters for K^+^ and Na^+^ ions consistent with the ECC framework, i.e. including effective electronic polarization, in MD simulations of potassium channels. We applied the same methodology to two fundamentally different force fields (Charmm36m and Amber14sb), ran simulations of three different K^+^ channels over the whole range of physiologically and experimentally relevant voltages. Despite minor differences between specific cases, our findings can be broadly summarized in three major points: 1. The introduction of the electronic polarization of cations (here through the ECC approach) is crucial to accurately model K^+^ ion permeation through potassium channels, allowing to simulate, for the first time, full I-V curves from atomistic MD simulations that are in very good agreement with experimental data; 2. This efficient K^+^ permeation occurs in all cases through the direct knock-on mechanism, with an even higher number of ions occupying the SF than seen in previous simulations, aided by reduced ion-ion repulsion due to electronic polarization (Fig. 7). Specifically, a highly conductive K^+^ channel can accommodate 3 or 4 ions in the SF. Furthermore, achieving balanced free energy between the 3-ion and 4-ion states is crucial for facilitating low barrier permeation (Fig. 7). Consequently, throughout all simulations conducted in this study, virtually no water molecules cross any potassium channels; 3. A careful adjustment of ion LJ parameters is necessary to recover correct ion radii and solvation free energies, which, as we showed, is critical for correct K^+^/Na. selectivity in K. channels. We therefore postulate, based on our and other works^4,16^, that the conductive state of the SF is characterized by near-full occupancy of K^+^ in many conditions.

**Figure 6.**
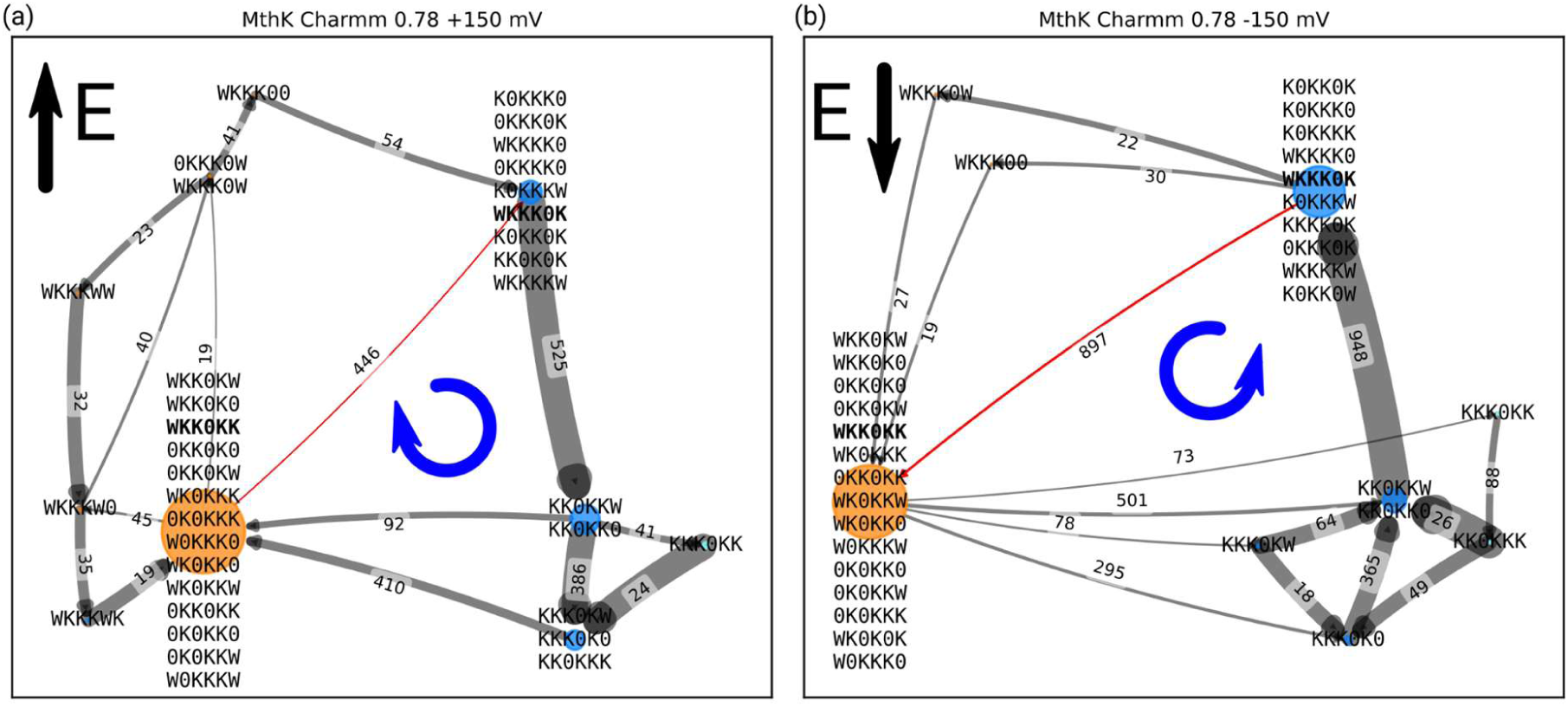
The mechanism graph from simulations of MthK in Charmm36m with 0.78 scaling under +150 mV or -150 mV. 515 (a) and 970 (b) permeation events were observed in 500 ns x 10 replica simulations at each voltage, respectively. The numbers shown by the edges are the net flux. The thickness of the edge (arrow) is proportional to the rate (defined as the inverse of the mean first passage time). The size of a node corresponds to the overall population of all states in this node. Nodes with 4 K^+^ ions in SF are shown in orange, and nodes with 3 K^+^ ions in SF are shown in blue. In panel (a), states A and B are lumped together when rate exceeds 125.0 ns^-1^, similarly, in panel (b), they are lumped when this product exceeds 180.0 ns^-1^. The rate-limiting step is marked in red. The exact flux and rate between all state pairs in the rate limiting step can be found in Table S4. The state pair WKKK0K-WKK0KK contributes to more than 85% of the net flux, and is identified as the rate-limiting step without lumping.

**Figure 7.**
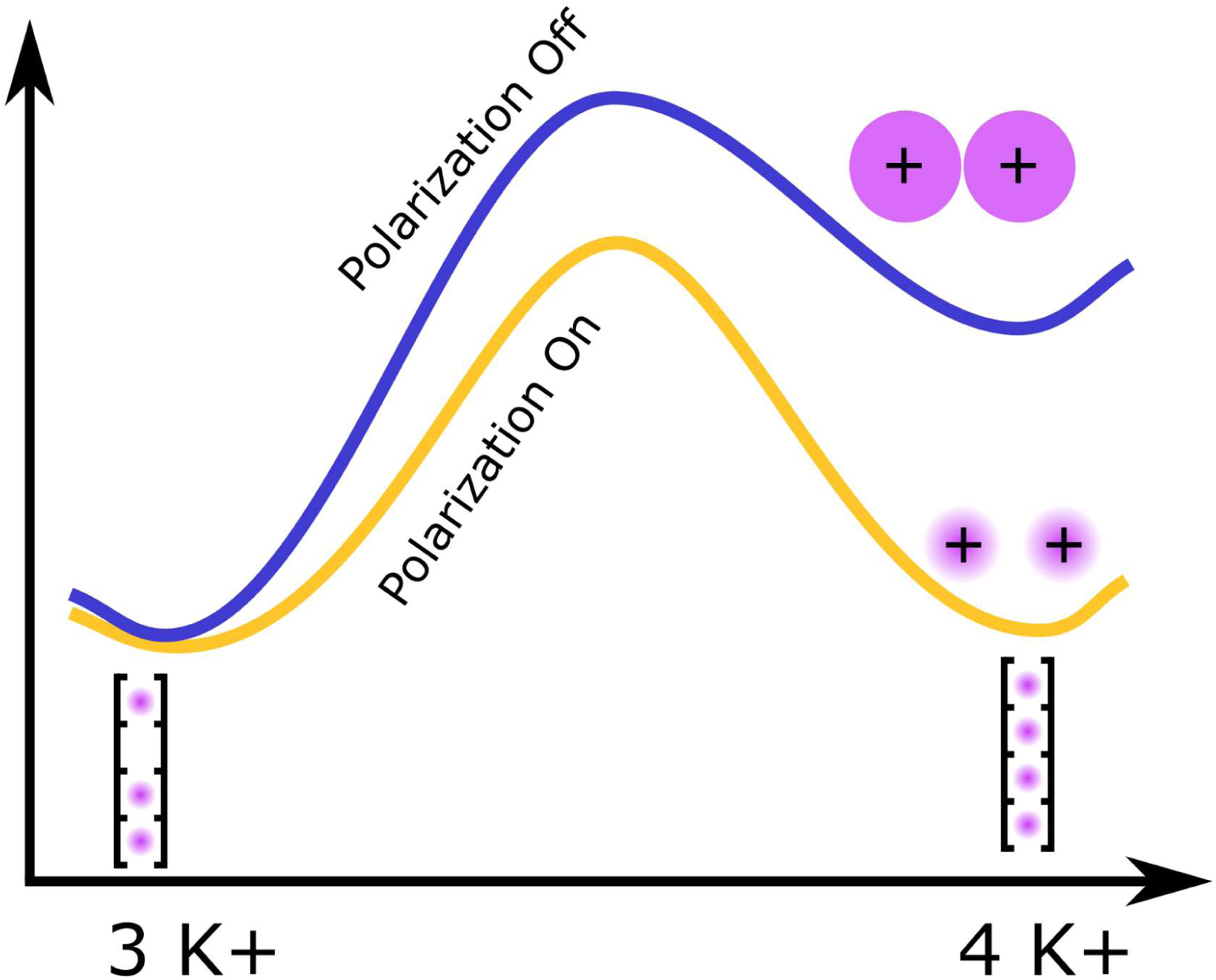
Schematic energy landscape between 3 and 4 K^+^ states with/without polarization. Only 1 SF state is drawn for 3 and 4 K^+^ states, and in reality, multiple 3 K^+^ and multiple 4 K^+^ states are visited. When the ECC polarization is introduced, the dominating effect is a decreased ion-ion repulsion, leading to more ion-ion contacts and fully ion filled filters, as well as a reduced barrier to form such contacts, and thus a more efficient knock-on permeation.

In our simulations, the best overall performing force field is CHARMM36m at a scaling factor of 0.78 with radius-corrected LJ parameters for ECC ions. This force field correctly reproduces I-V curves in all three channels, as well as their K^+^/Na^+^ selectivity and inward rectification in the MthK channel. We would therefore recommend it for simulations of other potassium channels, however slight adjustments of the scaling factor might result in a better performance for a specific channel.

Our newly developed parameters allow for MD simulations of K^+^ channels at an unprecedented level of accuracy. As almost any process involving K. channels is related to the SF behavior and its occupancy by K^+^ ions, we envision future simulation studies to benefit from these findings. Computational insights into the mechanisms of, for example, current regulation, selectivity filter gating or channel modulation by small molecules are now accessible at a much higher accuracy, especially as current readout is now readily accessible at unprecedented accuracy.

## 4. Methods

### 4.1. System preparation

#### 4.1.1. NaK2K

The crystal structure 3ouf^1^ was used and oriented in the membrane using OPM.^35^ 8 water molecules present in the crystal structure, behind the SF of NaK2K, were preserved. The current- increasing F92A mutation^1,36^, ACE and NME termini patching, membrane and solvation box building were performed using Charmm-GUI,^37,38^ webserver. The final system consisted of one tetrameric channel, 168 POPC molecules, 11,134 water molecules, 160 K^+^ ions, 152 Cl^-^ ions, corresponding to 0.8 m/L salt concentration.

#### 4.1.2. TRAAK

The crystal structure 4wfe^39^ from OPM was used for TRAAK. 6 water behind the SF was preserved. The I159C and R284C mutations were introduced based on experimental results indicating that this disulfide bond can prevent the channel from closing.^39^ In the previously reported WT simulation, a lipid can enter the pore from this fenestration site and possibly close the channel.^8^ 246 POPC molecules were added using Charmm-GUI.^37,38^ ACE and NME were added to cap the termini. The final system consisted of one channel, 246 POPC molecules, 27,973 water molecules, 403 K^+^ ions, 389 Cl^-^ ions, corresponding to 0.8 m/L salt concentration.

#### 4.1.3. MthK

For MthK the 3ldc^33^ crystal structure was used. This structure was modified using the PyMOL mutagenesis tool to perform H68S and C77V mutations to match the WT amino acid sequence.^40^ This structure was aligned using OPM. The Charmm-GUI Bilayer Builder was used to add ACE and NME capping groups and to create the simulation box. Crystal water and potassium ions were kept during this procedure. The final system contained 169 POPC molecules and 244 K^+^ ions, 232 Cl^-^ and 16,900 water molecules, corresponding to 0.8 m/L salt concentration.

#### 4.1.4. Kv1.2-2.1

The crystal structure of the Kv1.2-2.1 paddle chimera (2r9r^41^) from OPM was used. 8 water behind SF was preserved. The pore and voltage-sensing domains (E144 to T417) were embedded in a POPC membrane using Charmm-GUI. The functionally nonessential^42^ tetramerization domain (T1) and the regulatory β subunit were omitted. ACE and NME capping were added in Charmm-GUI. The final system contained 313 POPC molecules and 460 K^+^ ions, 452 Cl^-^ and 31,908 water molecules, corresponding to 0.8 m/L salt concentration.

#### 4.1.5. KcsA

A pre-equilibrated structure of the E71A mutant (that removes C-type inactivation), including 17 water molecules behind and on top of the SF, based on the 5vk6^43^ crystal structure as used in previous studies^44,45^ was embedded in a POPC membrane using Charmm-GUI. ACE and NME capping groups were used for the N- and C-terminus respectively. GLU118 and GLU120 were protonated to mimic the state at pH 4. The final system contained 153 POPC molecules, 234 K^+^ ions, 250 Cl^-^ and 16,196 water molecules, corresponding to 0.8 m/L salt concentration.

### 4.2. Force field-specific parameters

#### 4.2.1. Charmm36m

For the Charmm36m^46^ force field, Charmm lipid^47^, Charmm TIP3P^48^, and Charmm ions^49^ were used. A cutoff of 1.2 nm was used for van der Waals (vdW) interactions, and the forces were switched smoothly to zero between 1.0 to 1.2 nm. The particle mesh Ewald (PME) method with a 1.2 nm cutoff was used for long-range electrostatic interactions.

#### 4.2.2. Amber14sb

For the Amber14sb^50^ force field, lipid21^51^, TIP3P water^52^, and Joung and Cheatham ion parameters^53^ were used. A cutoff of 1.0 nm was used for van der Waals (vdW) interactions, and a long-range analytical dispersion correction was applied to the energy and pressure. The PME method with a 1.0 nm cutoff was used for long-range electrostatic interactions.

#### 4.2.3. Rescaled force fields

In one simulation box, the partial charges for all the charged residues including ions and charged amino acids were adjusted with the same scaling factor, resulting in overall neutral simulation systems. The partial charges for the charged amino acids were adjusted as follows. In Charmm36m the partial charges of the charge group contributing to the net charge were multiplied by the selected scaling factor. For example, in ASP the -CH_2_-CO_2_ group has a charge of -1. When charge scaling is applied to ASP, the charge of the -CH_2_-CO_2_ group is scaled with the scaling factor (Fig. S21). This approach could not be used in Amber, because the partial charges are also distributed over the backbone atoms. Therefore, the scaled charges were obtained by taking a weighted average (scaling factor as the weight) over the protonated and deprotonated charges (Fig. S21). The deprotonated arginine, that was not available in the original amber14sb distribution, was optimized at MP2/6-311G(d) using Gaussian 09 and RESP charge was fitted to HF/6-31G(d) with antechamber.^54,55^ All the partial charges used in this work at every scaling factors are provided in the github repository: https://github.com/deGrootLab/Charge_Scaling_in_Potassium_Channel_Simulations_paper/

#### 4.2.4. Radius corrected vdW for charge scaled ion

The radius corrected vdW for the charge scaled ion was obtained and validated in the following 3 steps.

**Step 1**: Sigma optimization with grid search.

At each 0.05 charge from 1.00 (1.00, 0.95, 0.90 …) the sigma parameter of the ion decreased to the point when the radius of the first solvation shell reproduced the original radius in 1.00 charge. This sigma optimization was done by a grid search (a grid with (sigma, charge) pair). The setting and results of this grid search can be found in Fig. S8, and Fig. S9. The first peak in the ion-water oxygen RDF was used as the radius of the first solvation shell. Epsilon was kept the same as the original paper in the respective force field.^49,53^ The sigma parameter between 0.05 charge (0.99, 0.98, 0.97, 0.96, 0.94 …) was linearly interpolated. The optimized vdW for K^+^, Na^+^, Cl^-^ is listed in Table S5.

In this step a simulation box with 2163 water molecules and 1 ion was used. Each (sigma, charge) pair on the grid was simulated for 40 ns, and trajectory was saved every 0.2 ps. Each grid (K^+^, Na^+^, Cl^-^) was run in a Hamiltonian replica exchange simulation, and an exchange is attempted every 10 ps. RDF between ion and water oxygen is computed with “gmx rdf”, using a bin width of 0.002 nm. The highest 7 points in the RDF were fitted with a Gaussian function (illustrated in Fig. S11). The same non-bonded settings for the corresponding force fields mentioned in Sections 4.2.1 and

#### 4.2.2 were used

**Step 2**: Solvation free energy validation

The ion solvation free energy calculation was performed using Gromacs 2022.5. The non-bonded parameters were set the same as previously mentioned in 4.2.1 and 4.2.2. Temperature was controlled by langevin integrator at 298 K and pressure was set at 1 bar using Parrinello-Rahman. The Coulomb interaction was gradually turned off in 10 lambda windows (0.0, 0.1, 0.2, 0.32, 0.44, 0.56, 0.68, 0.8, 0.9, 1.0, window 1-10), and vdw interaction was then turned off in the next 11 windows (0.0, 0.1, 0.2, 0.3, 0.4, 0.5, 0.6, 0.7, 0.8, 0.9, 1.0, window 10-20). HRE was used to speed up the sampling with an exchange attempt every 100 steps (0.2 ps). Each lambda window was simulated for 3 ns. The ∂H/∂λ and energy were saved every 2 ps. At least 1000 uncorrelated samples were recorded in each lambda window. “Alchemical-analysis” tool^56^ was used and the reported value was estimated by the Multistate Bennett Acceptance Ratio (MBAR) method. A soft-core potential^57^ was employed for the van der Waals interactions. The solvation free energy is shown in Fig. 4 and Fig. S10.

**Step 3**: RDF validation in dilute solution and 4M solution.

The RDF was validated under two conditions: (1) an infinitely diluted system, where the same simulation box as in step 1 was used, and (2) a 4M KCl solution, where the concentration matched that of the experiment.^58^

In the infinitely diluted system, each condition (charge with optimized sigma) was simulated for 100 ns and the trajectory is saved every 2 ps. The highest 7 points in the RDF were fitted with a Gaussian function (Fig. S11, Fig. S12). The radius is shown in Fig. 4 and Fig. S10.

A 4M system was constructed to compare with the neutron scattering experiment. The simulation box consisted of 6661 water, 120 K^+^ ions, and 120 Cl^-^ ions. Each condition (charge with optimized sigma) was simulated for 20 ns, and the trajectory was saved every 1 ps. The experimental RDF was provided by the authors of the publication.^58^ The RDF comparison is shown in Fig. S13.

All the initial structures, topology, and MD parameter files are provided in the github repository: https://github.com/deGrootLab/Charge_Scaling_in_Potassium_Channel_Simulations_paper/

### 4.3. MD simulation

#### 4.3.1. Equilibration and general setting

Gromacs 2022.5 was used except for the charge-scaled HRE.^59,60^ The initial structure was minimized using the steepest descent until the maximum force was lower than 1000 kJ/(mol*nm). 100 ps of NVT with a harmonic restraint of 1000 kJ/(mol*nm^2^) on all heavy atoms of protein and POPC. The temperature in this equilibration was controlled by either the Berendsen (Charmm36m) or v-rescale (Amber14sb) thermostat at 310 K. An additional 110 ns NPT equilibration was performed to relax the membrane with harmonic restraints of 1000 kJ/(mol*nm^2^) on all heavy atoms of the protein. The temperature was controlled by either the Berendsen (Charmm36m) or V-rescale (Amber14sb) thermostat at 310 K. The semi-isotropic pressure coupling was controlled by the Berendsen barostat at 1 bar. The averaged box z dimension in the last 100 ns was used to set the electric field. In the later production run, we switched to the Nose-Hoover (for Chamm36m) or V-rescale (for Amber14sb) thermostat and the Parrinello-Rahman (for both Charmm36m and Amber14sb) barostat. A time step of 2 fs was used, and bonds with Hydrogen atoms were constrained by LINCS.^61^

#### 4.3.2. Non-equilibrium simulation under voltage

To simulate the channel under voltage, an electric field is applied.^62^ In Charmm36m and Amber14sb, the simulation was started after the equilibration, while in Amber14sb-S3, they were started from the ensemble generated by HRE with randomly generated velocity at 310 K. For every condition (including channel, force field, scaling factors, and voltage), 10 replicas of 500 ns were performed.

When counting the K^+^ current crossing the channel, we assume that 1 K^+^ ion contributes to 1 elementary charge even if the partial charge is less than 1. The scaling of the partial charge is a mean field treatment to approximate the polarization of the electron cloud between the nuclei, thus the electric field (voltage) in the simulation is set the same for the 1.00 charge force field and a charge scaled force field, and transporting 1 K^+^ ion would generate the current of 1 elementary charge regardless of the parameter.

#### 4.3.3. Hamiltonian replica exchange (HRE)

The HRE simulations were performed on Gromacs 2021.7 with plumed 2.8.2.^63–65^ Although Gromacs has its own HRE implementation, it only works with the free energy code and can only interpolate states using the lambda definition. The actual interpolation can be found in Gromacs documentation ( https://manual.gromacs.org/2022.5/reference-manual/functions/free-energy-interactions.html).^59,60^ In the Plumed Hamiltonian Replica Exchange (HRE) setup, each replica’s Hamiltonian can be defined independently using separate topology files. We opted for Plumed HRE because this implementation allows every replica to have physically meaningful parameters, enabling us to assign distinct charges to each replica. Replica exchange is meaningful only in equilibrium conditions, so no voltage is applied. 36 replicas were set up with charge scaling factors decreasing from 1.00 to 0.65 in steps of 0.01, using the same simulation boxes as mentioned in the system preparation section. Every 100 MD steps (0.2 ps), an exchange attempt was tested. Each replica was simulated for 80 ns and the trajectory was saved every 20 ps. In our simulation practice, we found the HRE approach to be an efficient method to probe the scaling factor space. The conductance-scaling bar plot in Fig. 2(a) was obtained from 40 μs simulations with a scaling factor from 1.00 to 0.65 and a step of 0.05 (8 scaling factors, 10 replicas per scale, 500 ns every replica), while the SF state population plot in Fig. 2(d) was obtained from 2.88 μs simulation on the same simulation box with a scaling factor from 1.00 to 0.65 and a step of 0.01 (36 replicas, 80 ns each). Using HRE, with a fraction of the computation power, the high conductance parameters can be quickly located with fine resolution.

#### 4.3.4. Comparison of solvation free energy with experiment

The solvation free energy calculated from the scaled ion cannot be directly compared with the experiment, because part of the energy from electric polarization (E_ele_) is removed from the force field (E_ff_) and treated in the mean field approximation. (Eq. 1) The actual solvation free energy can be splitted into 2 terms, ΔG_nuc_ which can be computed from MD solvation free energy calculation, and ΔG_ele_ which comes from the mean field approximation.

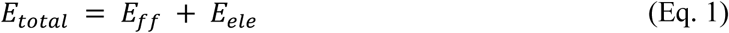

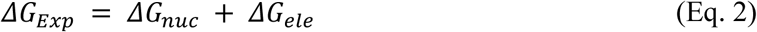

Assuming the approximated medium between MD particles has a relative dielectric constant of ε_r_, the ΔG_ele_ can be estimated with Born model (Eq. 3)

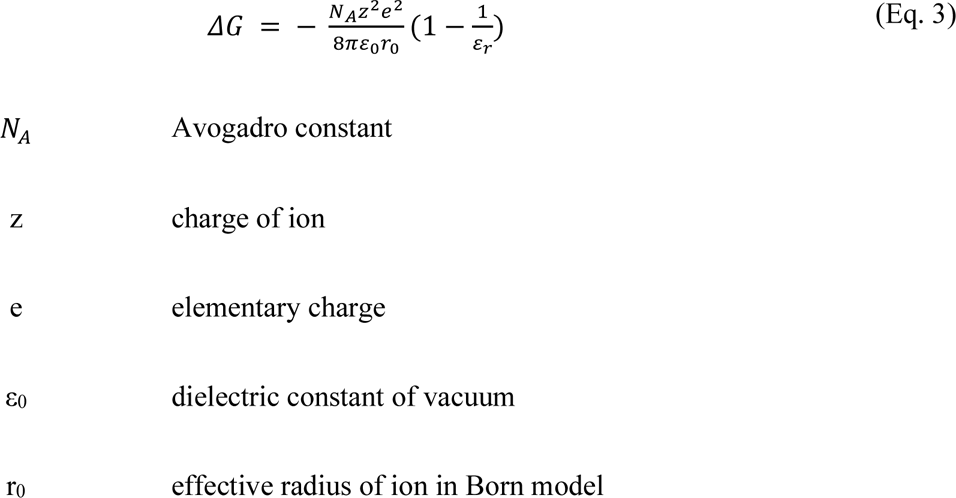

If we take scaling factor q as 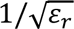 , Eq. 3 becomes Eq. 4.

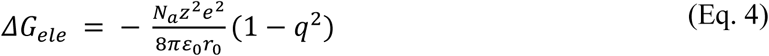

The relative dielectric constant of water is significantly greater than 1 (ε_r_ for water = 78.4). We approximate the experimental solvation free energy using the Born model, as shown in Eq. 5.

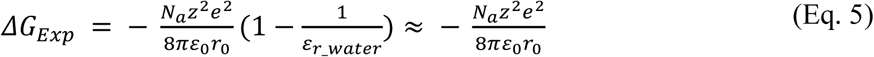

So that the MD simulated solvation free energy should target the rest of the solvation free energy, which is Eq. 6.

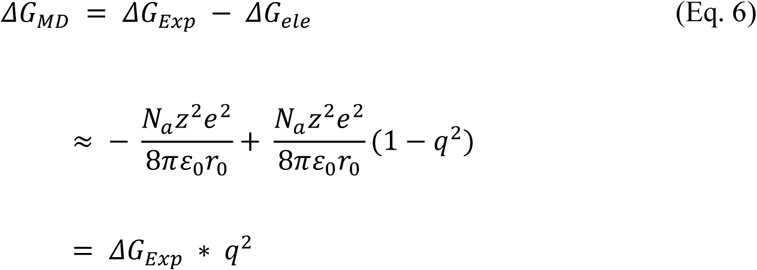

The comparing of experimental solvation free energy with ECC ion was discussed in several other references.^24,66^

### 4.4. Mechanism Graphs

The simulation was saved every 0.2 ps. The trajectory was discretized by the SF state. The rate between two states was defined as the inverse mean first passage time. All of the rates were first listed (every state pair, forward and backward), and the mechanism graph was progressively lumped from the state pair which had the fastest transition. If at least one of the forward or backward rates between a pair of states exceeded the cutoff rate, this pair of states would be lumped together. The cutoff rate when the lumping stopped is listed under each graph.

## 5. Acknowledgments

We would like to thank Youxing Jiang, Thomas Boulin, Dawon Kang, and Brad Rothberg for kindly sharing the experimental IV curves of NaK2K^1^, TRAAK^30^, and MthK^10^. We also thank Carter J. Wilson for providing the RESP charge of deprotonated arginine and Andrei Mironenko for providing the KcsA structure. R.d.V., W.K. and B.L.d.G. acknowledge funding from the Leibniz Collaborative Excellence Project “Ion Selectivity and Conduction Mechanism of Cation Channels” K305/2020. W.K and B.L.d.G. acknowledge funding from the German Research Foundation (DFG) through FOR2518 ’Dynion’, Project P5.

## 6. Author contributions

W.K. and B.L.d.G. conceived and supervised the project. C.H. and R.d.V. performed all computational work and analyzed the data. C.H., R.d.V., W.K. and B.L.d.G. wrote the manuscript.

