## Supplementary information file for "Effective Polarization in Potassium Channel Simulations: Ion Conductance, Occupancy, Voltage Response, and Selectivity"

| **NaK2K** | | | | |
| --- | --- | --- | --- | --- |
| | Charmm36m | | Amber14sb | |
| | state | population (%) | state | population (%) |
| 3 ion | KK0K | 58 | K0KK | 21.2 |
| | K0KK | 12 | KK0K | 9.6 |
| 2 ion | WK0K | 12.8 | 0KKW | 36.8 |
| | 0K0K | 5.7 | 0KK0 | 11.9 |
| | K0K0 | 4.3 | WKKW | 8.3 |
| | 0KKW | 1.9 | K0KW | 4.2 |
| | KWK0 | 1.6 | WK0K | 3.6 |
| | K0KW | 1.5 | WKK0 | 3.5 |
| | 0KK0 | 1 | | |
| **TRAAK** | | | | |
| | Charmm36m | | Amber14sb | |
| | state | population (%) | state | population (%) |
| 3 ion | KK0K | 77 | KK0K | 31.2 |
| | K0KK | 3.6 | K0KK | 12.2 |
| 2 ion | 0K0K | 8.9 | 0KK0 | 35.9 |
| | WK0K | 4.2 | WKK0 | 8.7 |
| | 0KKW | 2.2 | 0KKW | 5.5 |
| | 0KK0 | 1.7 | WK0K | 2.8 |
| | K0KW | 1.3 | WKKW | 2 |
| MthK | | | | |
| | Charmm36m | | Amber14sb | |
| | state | population (%) | state | population (%) |
| 3 ion | KK0K | 45.2 | KK0K | 13.9 |
| | K0KK | 4.1 | K0KK | 9.7 |
| 2 ion | 0K0K | 28.9 | 0KK0 | 50.7 |
| | K0K0 | 17 | WKK0 | 14.7 |
| | WK0K | 3.5 | 0KKW | 3.8 |
| | | | 0K0K | 3 |
| | | | K0KW | 2.6 |

Table S1: SF state population in NaK2K, TRAAK, MthK with Charmm36m and Amber14sb in 150 mV simulations. SF states with a population above 1% are listed.

Fig. S1: SF oxygen charge in Charmm36m, Amber14sb, and our modified Amber14sb-S3

| | Charmm36m | Amber14sb | Amber14sb-S3 |
|---|---|---|---|
| O-Tyr | -0.51 | -0.5679 | -0.5679 |
| O-Gly | -0.51 | -0.5679 | -0.5679 |
| O-Val | -0.51 | -0.5679 | -0.53 |
| O-Thr | -0.51 | -0.5679 | -0.53 |
| O-Thr | -0.66 | -0.6761 | -0.6761 |

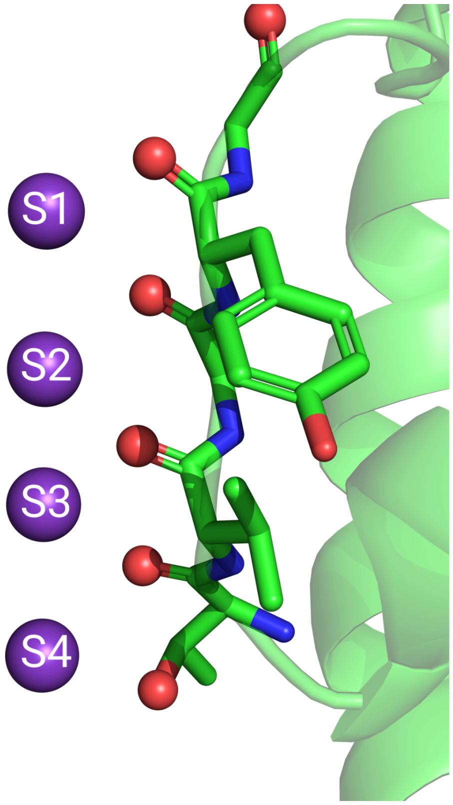

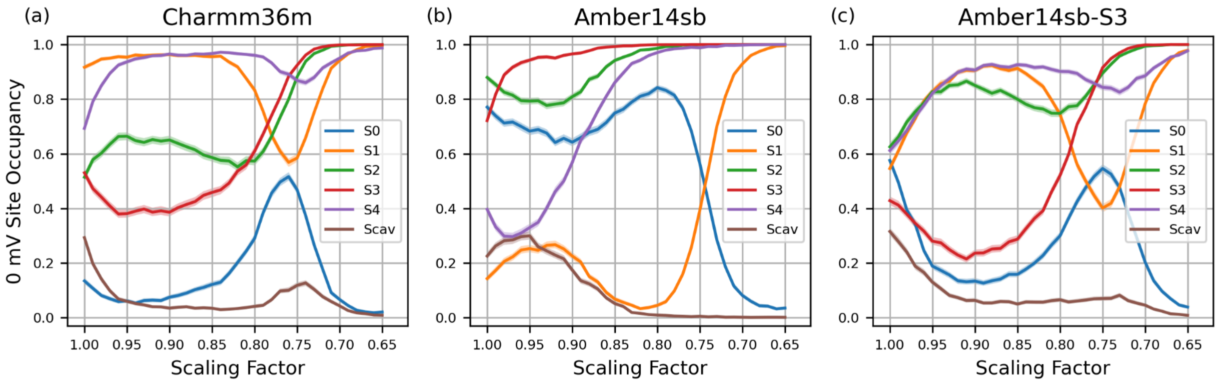

Fig. S2: SF state distribution at different scaling factors in force fields in NaK2K. **(a-c)** Occupancy of individual ion binding sites by a K+ ion from HRE simulation. The error bar is the 95 % confidence interval when bootstrapping frames.

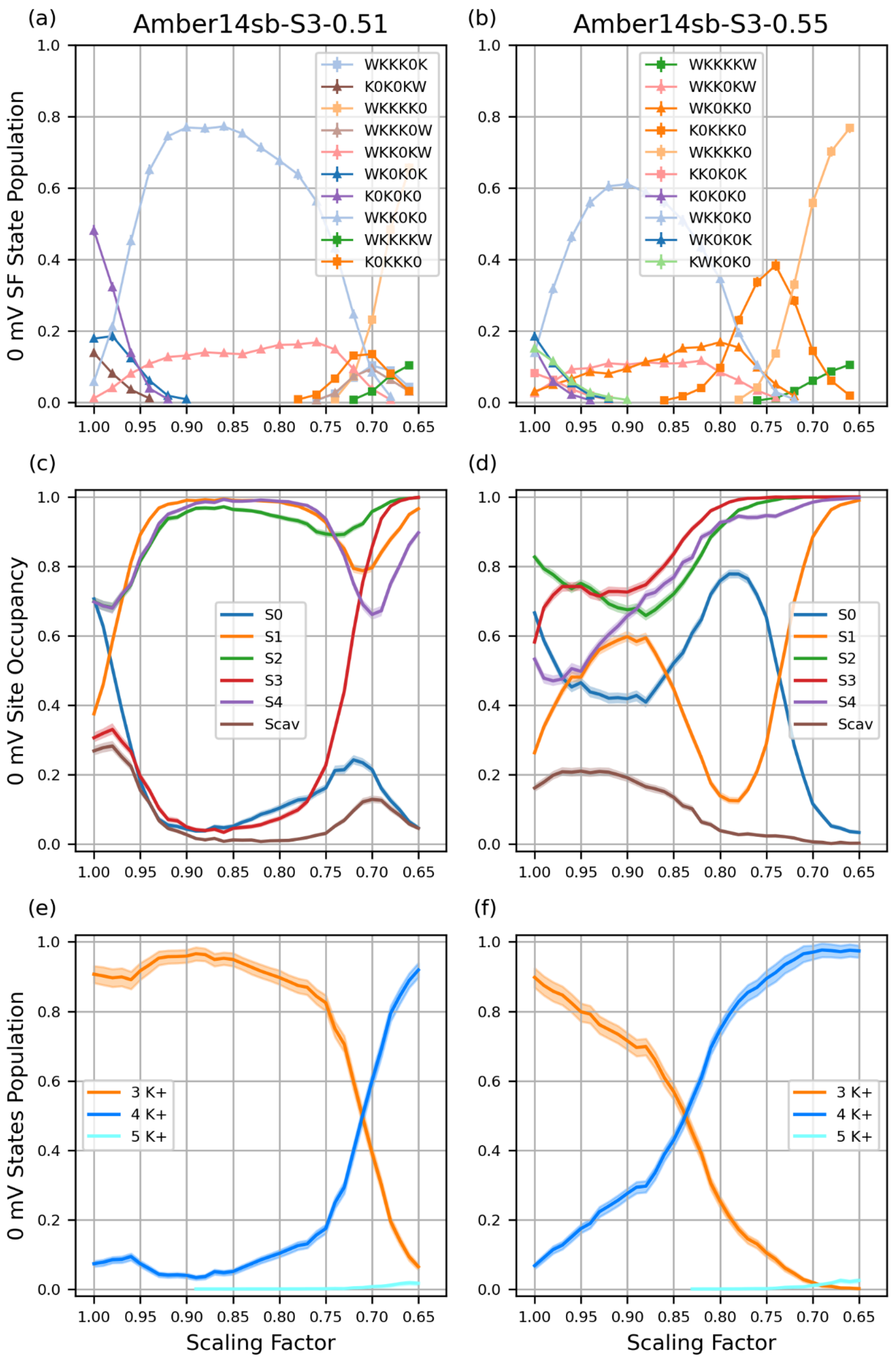

(a)
Amber14sb-S3-0.51
(b)
Amber14sb-S3-0.55
0 mV SF State Population
WKKK0K
K0K0KW
WKKKK0
WKKK0W
WKK0KW
WK0K0K
K0K0K0
WKK0K0
WKKKKW
K0KKK0
WKKKKW
WKK0KW
WK0KK0
K0KKK0
WKKKK0
KK0K0K
K0K0K0
WKK0K0
WK0K0K
KWK0K0
(c)
(d)
0 mV Site Occupancy
S0
S1
S2
S3
S4
Scav
(e)
(f)
0 mV States Population
3 K+
4 K+
5 K+
Scaling Factor

Fig. S3 : SF state distribution at different scaling factors in Amber14sb with different S3 modification in NaK2K. **(a-b)** Equilibrated populations of the SF states recorded in HRE in NaK2K, using a 6-letter code showing the occupancy at binding sites S0 to Scav: 'K' for a K+ ion, 'W' for water, and '0' for an unoccupied site. For clarity, only the even replicas are plotted (replica 0, 2, 4 etc at scale 1.00, 0.98, 0.96 etc). The odd replicas are not shown (replica 1, 3, 5 etc at scale 0.99, 0.97, 0.95 etc). **(c-d)** Occupancy of individual ion binding sites by a K+ ion from HRE simulation. **(j-l)** Ion occupation state distribution from HRE simulation, similar to (c-d), but the states with the same number of ions in the SF are lumped together. In (a-f) The error bar is the 95 % confidence interval when bootstrapping frames.

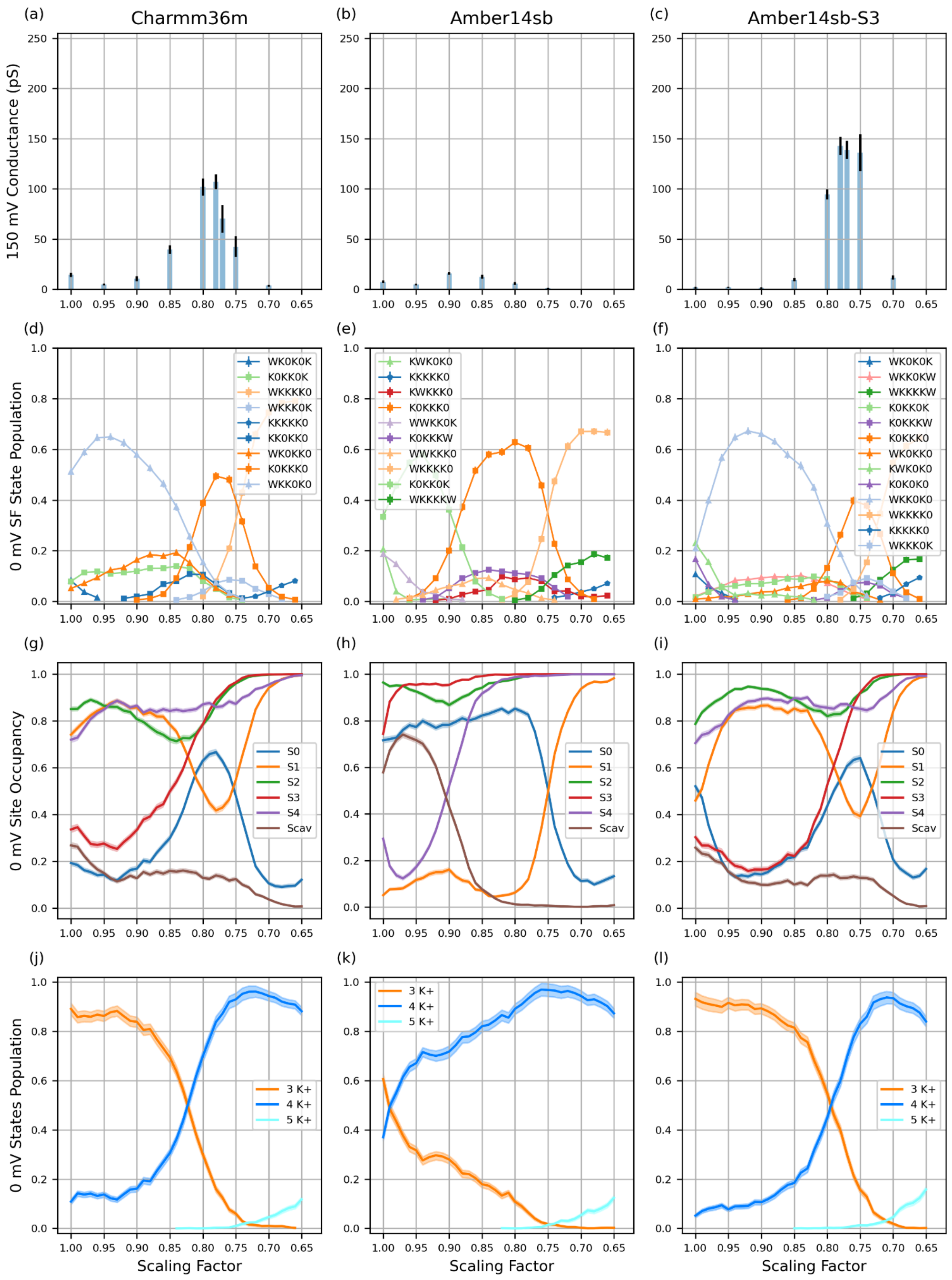
(a)
Charmm36m
(b)
Amber14sb
(c)
Amber14sb-S3
150 mV Conductance (pS)
(d)
(e)
(f)
0 mV SF State Population
WK0K0K
K0KK0K
WKKKK0
WKKK0K
KKKKK0
KK0KK0
WK0KK0
K0KKK0
WKK0K0
KWK0K0
KKKKK0
KWKKK0
K0KKK0
WWKK0K
K0KKKW
WWKKK0
WKKKK0
K0KK0K
WKKKKW
WK0K0K
WKK0KW
WKKKKW
K0KK0K
K0KKKW
K0KKK0
WK0KK0
KWK0K0
K0K0K0
WKK0K0
WKKKK0
KKKKK0
WKKK0K
(g)
(h)
(i)
0 mV Site Occupancy
S0
S1
S2
S3
S4
Scav
(j)
(k)
(l)
0 mV States Population
3 K+
4 K+
5 K+
Scaling Factor

Fig. S4: Conductance and SF state distribution at different scaling factors in force fields in TRAAK. (**a-c**) Average conductances at different scaling factors in MD simulations using Charmm36m, Amber14sb, and Amber14sb-S3 (see text for explanation) force fields under +150 mV in NaK2K. The error bar is the standard error of the mean (N=10). **(d-f)** Equilibrated populations of the SF states recorded in HRE in NaK2K, using a 6-letter code showing the occupancy at binding sites S0 to Scav: 'K' for a K+ ion, 'W' for water, and '0' for an unoccupied site. For clarity, only the even replicas are plotted (replica 0, 2, 4 etc at scale 1.00, 0.98, 0.96 etc). The odd replicas are not shown (replica 1, 3, 5 etc at scale 0.99, 0.97, 0.95 etc). **(g-i)** Occupancy of individual ion binding sites by a K+ ion from HRE simulation. **(j-l)** Ion occupation state distribution from HRE simulation, similar to **(d-f)**, but the states with the same number of ions in the SF are lumped together. In **(d-l)** The error bar is the 95 % confidence interval when bootstrapping frames.

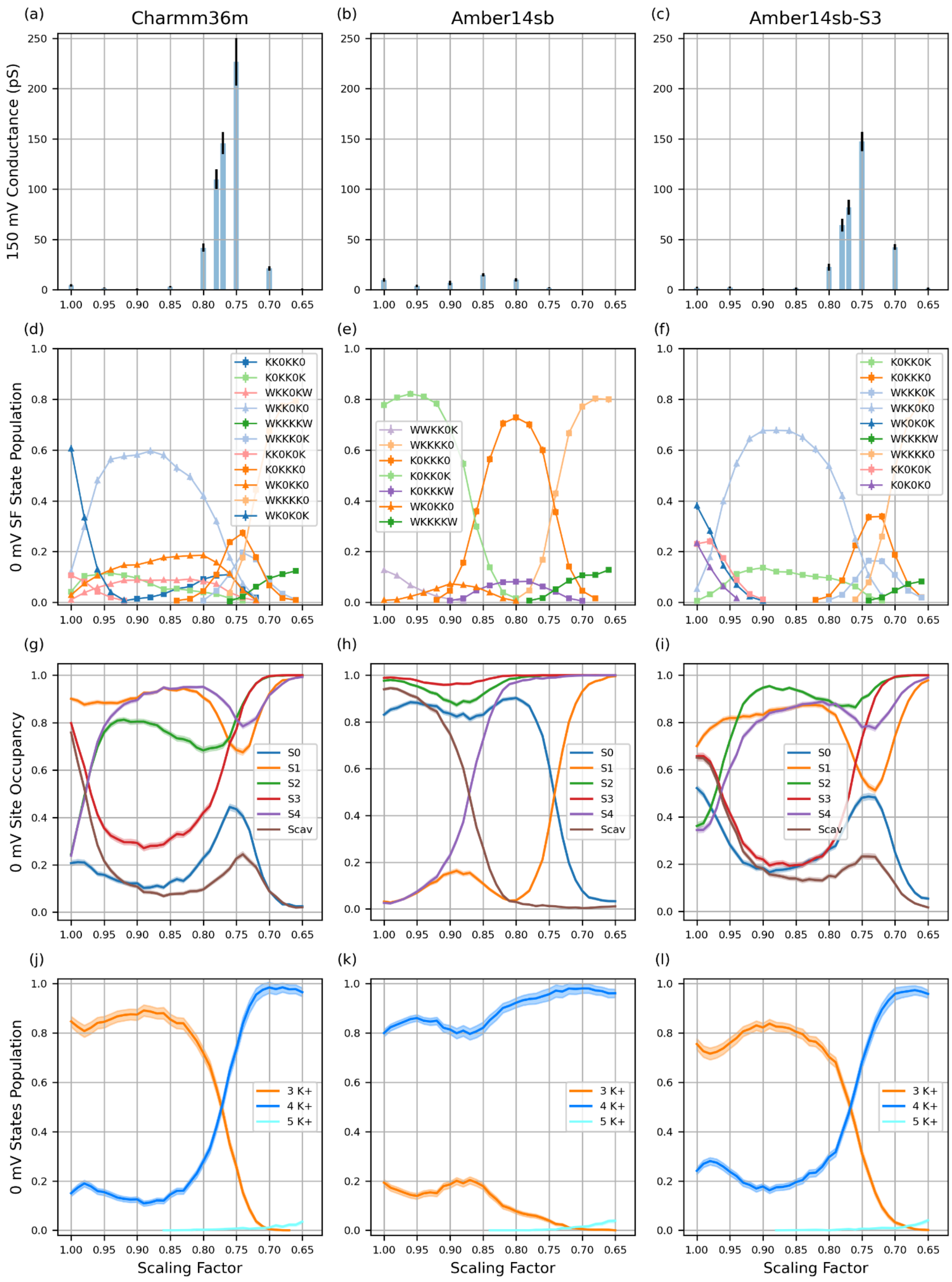

(a)
Charmm36m
(b)
Amber14sb
(c)
Amber14sb-S3
150 mV Conductance (pS)
(d)
(e)
(f)
0 mV SF State Population
KK0KK0
K0KK0K
WKK0KW
WKK0K0
WKKKKW
WKKK0K
KK0K0K
K0KKK0
WK0KK0
WKKKK0
WK0K0K
WWKK0K
WKKKK0
K0KKK0
K0KK0K
K0KKKW
WK0KK0
WKKKKW
K0KK0K
K0KKK0
WKKK0K
WKK0K0
WK0K0K
WKKKKW
WKKKK0
KK0K0K
K0K0K0
(g)
(h)
(i)
0 mV Site Occupancy
S0
S1
S2
S3
S4
Scav
(j)
(k)
(l)
0 mV States Population
3 K+
4 K+
5 K+
Scaling Factor

Fig. S5: Conductance and SF state distribution at different scaling factors in force fields in MthK. (**a-c**) Average conductances at different scaling factors in MD simulations using Charmm36m, Amber14sb, and Amber14sb-S3 (see text for explanation) force fields under +150 mV in NaK2K. The error bar is the standard error of the mean (N=10). **(d-f)** Equilibrated populations of the SF states recorded in HRE in NaK2K, using a 6-letter code showing the occupancy at binding sites S0 to Scav: 'K' for a K+ ion, 'W' for water, and '0' for an unoccupied site. For clarity, only the even replicas are plotted (replica 0, 2, 4 etc at scale 1.00, 0.98, 0.96 etc). The odd replicas are not shown (replica 1, 3, 5 etc at scale 0.99, 0.97, 0.95 etc). **(g-i)** Occupancy of individual ion binding sites by a K+ ion from HRE simulation. **(j-l)** Ion occupation state distribution from HRE simulation, similar to **(d-f)**, but the states with the same number of ions in the SF are lumped together. In **(d-l)** The error bar is the 95 % confidence interval when bootstrapping frames.

### 1. Additional tests in Kv1.2-2.1 and KcsA

To test the generalizability of the force field, we further tested 2 more channels, Kv1.2-2.1 and KcsA with the optimized parameters we obtained in simulating NaK2K, TRAAK, MthK. The same protocol of HRE and conductance simulation in other channels was applied.

For simulations under voltage (Fig. S7) we used the optimal parameters found for other channels, i.e. the 0.78 charge scaling factor and the radius-corrected LJ parameters for ions in Charmm36m. Both channels, simulated at ±150 mV, display conductances in a qualitative agreement with experimental ones (approximate values from experiments for Kv1.2: ±4 pA at ±90 mV, for KcsA: +35 pA at +150 mV, -10 pA at -150 mV).[1,2] Notably this means that we also reproduce the strong outward rectification of KcsA at high potassium concentrations.

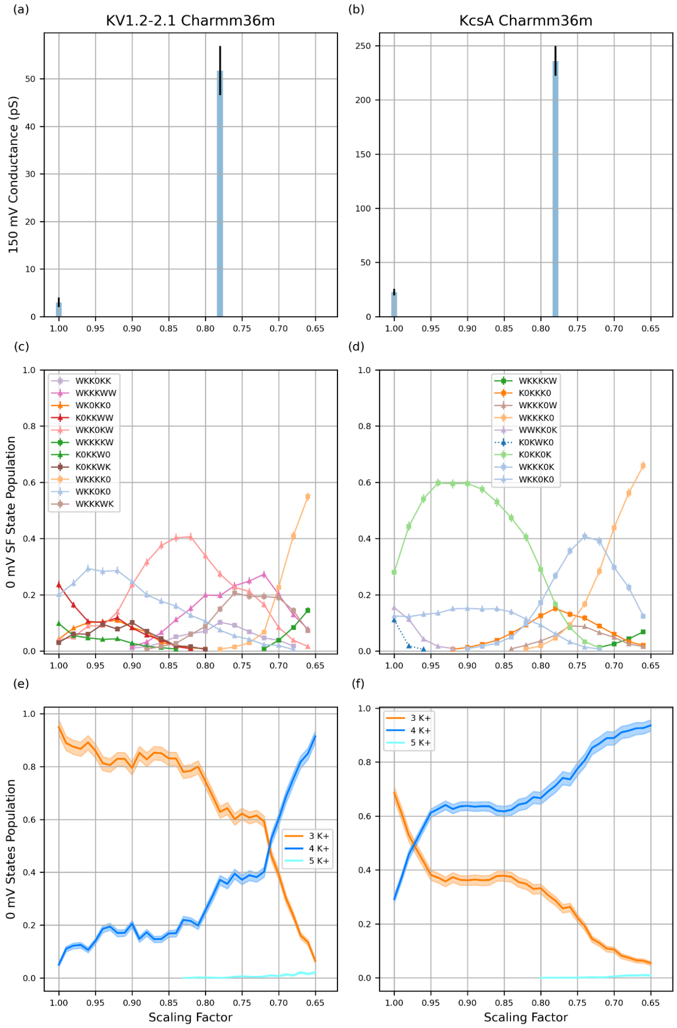

Fig. S6: Conductance and SF state distribution at different scaling factors in Charmm36m in KV1.2-2.1 and KcsA. **(a-b)** Average conductances at different scaling factors in MD simulations using Charmm36m, force fields under +150 mV in KV1.2-2.1 and KcsA. The error bar is the standard error of the mean (N=10). The y axis range is not the same in (a) and (b). **(c-d)** Equilibrated populations of the SF states recorded in HRE in KV1.2-2.1 and KcsA, using a 6-letter code showing the occupancy at binding sites S0 to Scav: 'K' for a $K^+$ ion, 'W' for water, and '0' for an unoccupied site. For clarity, only the even replicas are plotted (replica 0, 2, 4 etc at scale 1.00, 0.98, 0.96 etc). The odd replicas are not shown (replica 1, 3, 5 etc at scale 0.99, 0.97, 0.95 etc). **(e-f)** Ion occupation state distribution from HRE simulation, the states with the same number of ions in the SF are lumped together. In **(c-f)** The error bar is the 95 % confidence interval when bootstrapping frames.

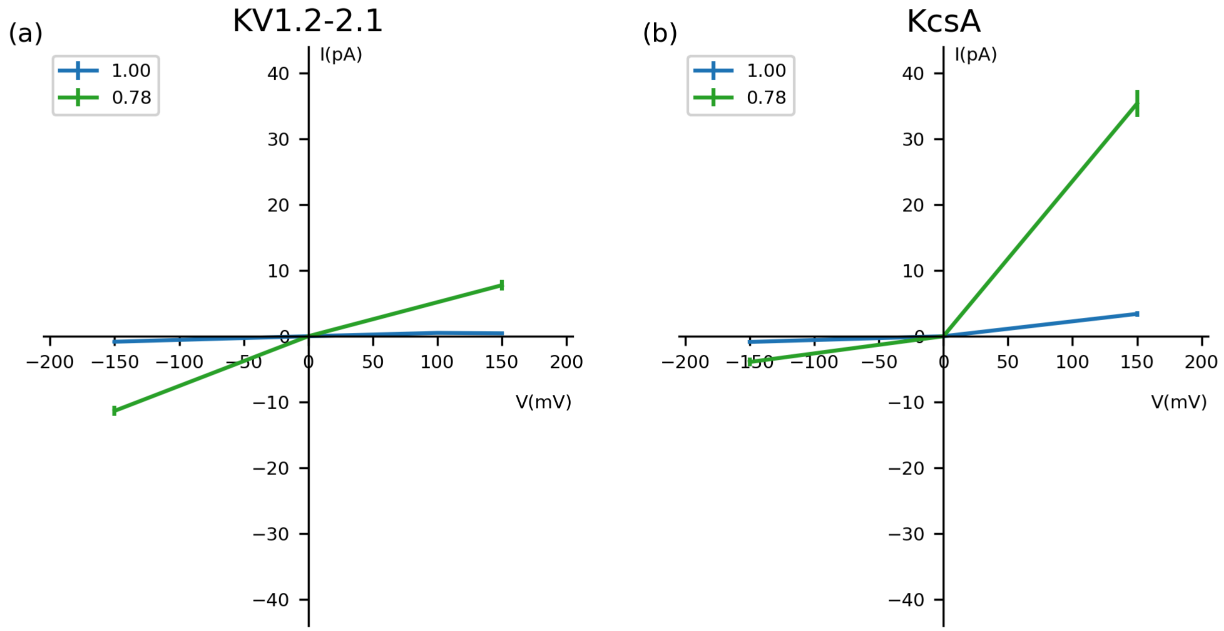

Fig. S7 Simulated current-voltage (I-V) curves for KV1.2-2.1 and KcsA channels at different scaling factors (1.00, and 0.78) in Charmm36m. Simulations were performed at ± 150 mV.

### 2. Simulation results in digital form

| NaK2K | | | | | | | | | |
|---|---|---|---|---|---|---|---|---|---|
| | -150 mV | | | | 150 mV | | | | |
| charge | G_ave | G_SEM | K perm | W perm | G_ave | G_SEM | K perm | W perm | force field |
| 1 | 7.5 | 1.9 | 35 | 0 | 9.2 | 1.7 | 43 | 1 | Charmm36m |
| 0.95 | 6.6 | 1 | 31 | 1 | 2.1 | 0.8 | 10 | 0 | Charmm36m |
| 0.9 | 5.6 | 1.2 | 26 | 0 | 1.5 | 0.6 | 7 | 0 | Charmm36m |
| 0.85 | 17.5 | 1.9 | 82 | 0 | 6.6 | 1.9 | 31 | 0 | Charmm36m |
| 0.8 | 113.4 | 8 | 531 | 0 | 62 | 6.7 | 290 | 0 | Charmm36m |
| 0.78 | 154.9 | 18 | 725 | 0 | 104.7 | 4.4 | 490 | 0 | Charmm36m |
| 0.77 | 155.5 | 9.7 | 728 | 0 | 128.4 | 5.6 | 601 | 0 | Charmm36m |
| 0.75 | 129.2 | 7.3 | 605 | 0 | 122.4 | 10.1 | 573 | 0 | Charmm36m |
| 0.7 | 8.8 | 1.2 | 41 | 0 | 10.7 | 2.8 | 50 | 0 | Charmm36m |
| 0.65 | 0.4 | 0.3 | 2 | 0 | 0 | 0.3 | 0 | 0 | Charmm36m |
| TRAAK | | | | | | | | | |
| | -150 mV | | | | 150 mV | | | | |
| charge | G_ave | G_SEM | K perm | W perm | G_ave | G_SEM | K perm | W perm | force field |
| 1 | 12.4 | 2.3 | 58 | 3 | 14.5 | 2.2 | 68 | 0 | Charmm36m |
| 0.95 | 9 | 1.3 | 42 | 3 | 4.9 | 1 | 23 | 0 | Charmm36m |
| 0.9 | 13.2 | 2 | 62 | 3 | 10.5 | 2.5 | 49 | 0 | Charmm36m |
| 0.85 | 50.2 | 7.4 | 235 | 0 | 39.7 | 4.3 | 186 | 0 | Charmm36m |
| 0.8 | 131 | 16.2 | 613 | 1 | 101.9 | 8.4 | 477 | 0 | Charmm36m |
| 0.78 | 112.4 | 18.2 | 526 | 0 | 92.1 | 12.1 | 431 | 0 | Charmm36m |
| 0.77 | 78 | 12 | 365 | 0 | 70.3 | 13.8 | 329 | 0 | Charmm36m |
| 0.75 | 38.2 | 4.9 | 179 | 0 | 42.5 | 10.2 | 199 | 0 | Charmm36m |
| 0.7 | 3.8 | 1 | 18 | 0 | 3.6 | 1 | 17 | 0 | Charmm36m |

| | | | | | | | | | |
|---|---|---|---|---|---|---|---|---|---|
| 0.65 | 0.2 | 0.2 | 1 | 0 | 0 | 0 | 0 | 0 | Charmm36m |
| MthK | | | | | | | | | |
| | -150 mV | | | | 150 mV | | | | |
| charge | G_ave | G_SEM | K perm | W perm | G_ave | G_SEM | K perm | W perm | force field |
| 1 | 2.8 | 0.6 | 13 | 0 | 4.5 | 1.2 | 21 | 0 | Charmm36m |
| 0.95 | 3.6 | 1 | 17 | 0 | 1.7 | 0.6 | 8 | 0 | Charmm36m |
| 0.9 | 1.5 | 0.5 | 7 | 0 | 0.6 | 0.5 | 3 | 0 | Charmm36m |
| 0.85 | 12.6 | 2 | 59 | 0 | 3 | 0.7 | 14 | 0 | Charmm36m |
| 0.8 | 114.5 | 10.6 | 536 | 0 | 41.9 | 4 | 196 | 0 | Charmm36m |
| 0.78 | 207.2 | 9.4 | 970 | 0 | 110 | 9.7 | 515 | 0 | Charmm36m |
| 0.77 | 219.8 | 8.3 | 1029 | 0 | 145.7 | 10.9 | 682 | 0 | Charmm36m |
| 0.75 | 237.3 | 11.4 | 1111 | 0 | 226.7 | 23.6 | 1061 | 0 | Charmm36m |
| 0.7 | 29.5 | 3.2 | 138 | 0 | 21.4 | 2.1 | 100 | 0 | Charmm36m |
| 0.65 | 0.2 | 0.2 | 1 | 0 | 0.6 | 0.5 | 3 | 0 | Charmm36m |
| NaK2K | | | | | | | | | |
| | | | | | 150 mV | | | | |
| charge | | | | | G_ave | G_SEM | K perm | W perm | force field |
| 1 | | | | | 20.1 | 1.8 | 94 | 0 | Amber14sb |
| 0.95 | | | | | 10.9 | 1.8 | 51 | -1 | Amber14sb |
| 0.9 | | | | | 21.4 | 3.5 | 100 | 0 | Amber14sb |
| 0.85 | | | | | 20.7 | 2.7 | 97 | -1 | Amber14sb |
| 0.8 | | | | | 10.5 | 2.2 | 49 | 0 | Amber14sb |
| 0.75 | | | | | 5.6 | 0.8 | 26 | 0 | Amber14sb |
| 0.7 | | | | | 0 | 0 | 0 | 0 | Amber14sb |
| 0.65 | | | | | 0 | 0 | 0 | 0 | Amber14sb |
| TRAAK | | | | | | | | | |
| | | | | | 150 mV | | | | |
| charge | | | | | G_ave | G_SEM | K perm | W perm | force field |
| 1 | | | | | 7.5 | 1.2 | 35 | 0 | Amber14sb |

| | | | | | | | | | |
|---|---|---|---|---|---|---|---|---|---|
| 0.95 | | | | | 4.9 | 0.7 | 23 | 0 | Amber14sb |
| 0.9 | | | | | 15.8 | 1.3 | 74 | 0 | Amber14sb |
| 0.85 | | | | | 12.4 | 2 | 58 | 0 | Amber14sb |
| 0.8 | | | | | 5.8 | 1.5 | 27 | 0 | Amber14sb |
| 0.75 | | | | | 1.1 | 0.6 | 5 | 0 | Amber14sb |
| 0.7 | | | | | 0 | 0 | 0 | 0 | Amber14sb |
| 0.65 | | | | | 0 | 0 | 0 | 0 | Amber14sb |
| MthK | | | | | | | | | |
| | | | | | 150 mV | | | | |
| charge | | | | | G_ave | G_SEM | K perm | W perm | force field |
| 1 | | | | | 9.8 | 1.7 | 46 | 0 | Amber14sb |
| 0.95 | | | | | 3.8 | 1.3 | 18 | 0 | Amber14sb |
| 0.9 | | | | | 6.8 | 2.3 | 32 | 0 | Amber14sb |
| 0.85 | | | | | 15 | 1.6 | 70 | 0 | Amber14sb |
| 0.8 | | | | | 10 | 1.6 | 47 | 0 | Amber14sb |
| 0.75 | | | | | 2.1 | 0.6 | 10 | 0 | Amber14sb |
| 0.7 | | | | | 0 | 0 | 0 | 0 | Amber14sb |
| 0.65 | | | | | 0 | 0 | 0 | 0 | Amber14sb |
| NaK2K | | | | | | | | | |
| | -150 mV | | | | 150 mV | | | | |
| charge | G_ave | G_SEM | K perm | W perm | G_ave | G_SEM | K perm | W perm | force field |
| 1 | 4.1 | 1.8 | 19 | 0 | 9.6 | 2.5 | 45 | 0 | Amber14sb-S3 |
| 0.95 | 4.5 | 0.9 | 21 | 0 | 5.1 | 0.9 | 24 | 0 | Amber14sb-S3 |
| 0.9 | 7.9 | 1.1 | 37 | 1 | 4.7 | 1.4 | 22 | 1 | Amber14sb-S3 |
| 0.85 | 22.9 | 3.6 | 107 | 1 | 11.5 | 2.4 | 54 | 1 | Amber14sb-S3 |
| 0.8 | 60.9 | 4.2 | 285 | 0 | 35.2 | 3.3 | 165 | -1 | Amber14sb-S3 |
| 0.78 | 76.3 | 8.9 | 357 | 2 | 58.5 | 3.5 | 274 | -2 | Amber14sb-S3 |
| 0.77 | 97.8 | 7.8 | 458 | -3 | 61.3 | 5.3 | 287 | 2 | Amber14sb-S3 |
| 0.75 | 81.8 | 11.1 | 383 | -1 | 88 | 3.2 | 412 | -1 | Amber14sb-S3 |
| 0.7 | 23.9 | 3 | 112 | 0 | 39.5 | 7 | 185 | 0 | Amber14sb-S3 |
| 0.65 | 1.5 | 0.8 | 7 | 0 | 1.1 | 0.4 | 5 | 0 | Amber14sb-S3 |

| | | | | | | | | | |
|---|---|---|---|---|---|---|---|---|---|
| TRAAK | | | | | | | | | |
| | -150 mV | | | | 150 mV | | | | |
| charge | G_ave | G_SEM | K perm | W perm | G_ave | G_SEM | K perm | W perm | force field |
| 1 | 1.9 | 0.7 | 9 | 0 | 1.9 | 0.9 | 9 | 0 | Amber14sb-S3 |
| 0.95 | 3 | 0.7 | 14 | 0 | 2.1 | 0.7 | 10 | 0 | Amber14sb-S3 |
| 0.9 | 4.9 | 1.2 | 23 | 2 | 1.5 | 0.6 | 7 | 0 | Amber14sb-S3 |
| 0.85 | 19.7 | 3.7 | 92 | 5 | 9.6 | 1.6 | 45 | 0 | Amber14sb-S3 |
| 0.8 | 140.6 | 12.9 | 658 | 1 | 94.4 | 5.1 | 442 | 0 | Amber14sb-S3 |
| 0.78 | 176.9 | 8.4 | 828 | 0 | 140.4 | 7.3 | 657 | 0 | Amber14sb-S3 |
| 0.77 | 110.4 | 18.6 | 517 | 3 | 138.9 | 9 | 650 | 0 | Amber14sb-S3 |
| 0.75 | 132.7 | 11.8 | 621 | 0 | 135.9 | 18.3 | 636 | 0 | Amber14sb-S3 |
| 0.7 | 11.3 | 1.8 | 53 | 0 | 11.7 | 2 | 55 | 0 | Amber14sb-S3 |
| 0.65 | 0.2 | 0.2 | 1 | 0 | 0.2 | 0.2 | 1 | 0 | Amber14sb-S3 |
| MthK | | | | | | | | | |
| | -150 mV | | | | 150 mV | | | | |
| charge | G_ave | G_SEM | K perm | W perm | G_ave | G_SEM | K perm | W perm | force field |
| 1 | 3.2 | 0.7 | 15 | 0 | 2.1 | 0.9 | 10 | 0 | Amber14sb-S3 |
| 0.95 | 2.6 | 0.6 | 12 | 0 | 2.3 | 0.9 | 11 | 0 | Amber14sb-S3 |
| 0.9 | 1.7 | 0.5 | 8 | 0 | 0.9 | 0.5 | 4 | 0 | Amber14sb-S3 |
| 0.85 | 7.9 | 1.4 | 37 | 0 | 1.7 | 0.7 | 8 | 0 | Amber14sb-S3 |
| 0.8 | 98.3 | 6.8 | 460 | 0 | 22.6 | 3.5 | 106 | 0 | Amber14sb-S3 |
| 0.78 | 132.4 | 9.2 | 620 | 0 | 57.7 | 2.4 | 270 | 0 | Amber14sb-S3 |
| 0.77 | 159.4 | 9.4 | 746 | 1 | 82 | 7.3 | 384 | 0 | Amber14sb-S3 |
| 0.75 | 162.6 | 18.4 | 761 | 1 | 147.4 | 9.7 | 690 | 0 | Amber14sb-S3 |
| 0.7 | 40.8 | 1.9 | 191 | 0 | 42.5 | 2.7 | 199 | 0 | Amber14sb-S3 |
| 0.65 | 0.2 | 0.2 | 1 | 0 | 1.3 | 0.9 | 6 | 0 | Amber14sb-S3 |

Table S2: Simulated conductance and number of permeation events. G_ave : Average conductance in 10 replicas. G_SEM : Standard error of the mean of conductance in 10 replicas. K perm : Total number of $K^+$ ion permeation events. W perm : Total number water permeation.

| NaK2K | | | | | | |
|---|---|---|---|---|---|---|
| | | -150 mV | | 150 mV | | |
| charge | site | K | Water | K | Water | force field |
| 1 | S0 | 0.19±0.02 | 1.46±0.03 | 0.12±0.01 | 1.34±0.01 | Charmm36m |
| | S1 | 0.84±0.01 | 0.02±0.00 | 0.89±0.01 | 0.00±0.00 | |
| | S2 | 0.30±0.01 | 0 | 0.78±0.01 | 0 | |
| | S3 | 0.86±0.01 | 0 | 0.25±0.01 | 0 | |
| | S4 | 0.34±0.01 | 0.02±0.01 | 0.89±0.01 | 0.05±0.01 | |
| | S5 | 0.66±0.01 | 0.04±0.00 | 0.07±0.01 | 0.08±0.01 | |
| 0.95 | S0 | 0.39±0.04 | 1.15±0.05 | 0.01±0.00 | 1.52±0.00 | Charmm36m |
| | S1 | 0.63±0.04 | 0.01±0.00 | 0.99±0.00 | 0.00±0.00 | |
| | S2 | 0.65±0.02 | 0.00±0.00 | 0.81±0.00 | 0 | |
| | S3 | 0.71±0.03 | 0.04±0.04 | 0.19±0.00 | 0 | |
| | S4 | 0.58±0.04 | 0.01±0.00 | 0.99±0.00 | 0.00±0.00 | |
| | S5 | 0.41±0.04 | 0.06±0.01 | 0.01±0.00 | 0.06±0.00 | |
| 0.9 | S0 | 0.31±0.01 | 1.31±0.02 | 0.02±0.00 | 1.61±0.00 | Charmm36m |
| | S1 | 0.73±0.02 | 0.00±0.00 | 1.00±0.00 | 0 | |
| | S2 | 0.59±0.01 | 0 | 0.77±0.00 | 0 | |
| | S3 | 0.68±0.01 | 0 | 0.23±0.00 | 0 | |
| | S4 | 0.73±0.02 | 0.00±0.00 | 1.00±0.00 | 0.00±0.00 | |
| | S5 | 0.27±0.02 | 0.05±0.00 | 0.00±0.00 | 0.07±0.00 | |
| 0.85 | S0 | 0.34±0.01 | 1.35±0.02 | 0.06±0.01 | 1.64±0.01 | Charmm36m |
| | S1 | 0.75±0.01 | 0.00±0.00 | 0.99±0.00 | 0.00±0.00 | |
| | S2 | 0.54±0.01 | 0 | 0.75±0.03 | 0 | |
| | S3 | 0.71±0.01 | 0 | 0.27±0.03 | 0 | |
| | S4 | 0.79±0.01 | 0.00±0.00 | 0.99±0.00 | 0.01±0.01 | |
| | S5 | 0.21±0.01 | 0.06±0.01 | 0.01±0.00 | 0.13±0.03 | |
| 0.8 | S0 | 0.41±0.01 | 1.26±0.01 | 0.17±0.02 | 1.53±0.02 | Charmm36m |
| | S1 | 0.68±0.01 | 0.00±0.00 | 0.95±0.01 | 0.00±0.00 | |
| | S2 | 0.60±0.01 | 0 | 0.70±0.03 | 0 | |
| | S3 | 0.78±0.02 | 0 | 0.39±0.02 | 0 | |
| | S4 | 0.87±0.00 | 0.00±0.00 | 0.95±0.02 | 0.05±0.03 | |
| | S5 | 0.14±0.01 | 0.08±0.01 | 0.03±0.01 | 0.20±0.05 | |
| 0.78 | S0 | | | 0.33±0.02 | 1.33±0.02 | Charmm36m |

| | | | | | | |
|---|---|---|---|---|---|---|
| | S1 | | | 0.84±0.01 | 0.00±0.00 | |
| | S2 | | | 0.66±0.01 | 0 | |
| | S3 | | | 0.55±0.02 | 0 | |
| | S4 | | | 0.95±0.01 | 0.03±0.01 | |
| | S5 | | | 0.03±0.00 | 0.16±0.02 | |
| 0.77 | S0 | 0.37±0.02 | 1.28±0.03 | 0.30±0.05 | 1.33±0.05 | Charmm36m |
| | S1 | 0.66±0.02 | 0.01±0.00 | 0.83±0.03 | 0.00±0.00 | |
| | S2 | 0.74±0.01 | 0 | 0.79±0.04 | 0 | |
| | S3 | 0.86±0.02 | 0 | 0.62±0.03 | 0 | |
| | S4 | 0.82±0.01 | 0.01±0.00 | 0.77±0.09 | 0.19±0.09 | |
| | S5 | 0.18±0.01 | 0.10±0.02 | 0.04±0.00 | 0.43±0.15 | |
| 0.75 | S0 | 0.24±0.04 | 1.34±0.04 | 0.49±0.07 | 1.08±0.07 | Charmm36m |
| | S1 | 0.77±0.04 | 0.00±0.00 | 0.63±0.05 | 0.00±0.00 | |
| | S2 | 0.87±0.00 | 0 | 0.83±0.03 | 0 | |
| | S3 | 0.89±0.03 | 0 | 0.84±0.01 | 0 | |
| | S4 | 0.74±0.02 | 0.04±0.03 | 0.84±0.08 | 0.12±0.08 | |
| | S5 | 0.23±0.02 | 0.14±0.05 | 0.06±0.01 | 0.22±0.09 | |
| 0.7 | S0 | 0.04±0.00 | 1.59±0.01 | 0.10±0.00 | 1.49±0.01 | Charmm36m |
| | S1 | 0.97±0.00 | 0.00±0.00 | 0.91±0.00 | 0 | |
| | S2 | 0.99±0.00 | 0 | 0.99±0.00 | 0 | |
| | S3 | 1.00±0.00 | 0 | 1.00±0.00 | 0 | |
| | S4 | 0.93±0.01 | 0.01±0.00 | 0.97±0.00 | 0.01±0.00 | |
| | S5 | 0.07±0.00 | 0.08±0.01 | 0.02±0.00 | 0.07±0.01 | |
| 0.65 | S0 | 0.03±0.00 | 1.69±0.01 | 0.02±0.00 | 1.63±0.03 | Charmm36m |
| | S1 | 1.00±0.00 | 0 | 1.00±0.00 | 0 | |
| | S2 | 1.00±0.00 | 0 | 1.00±0.00 | 0 | |
| | S3 | 1.00±0.00 | 0 | 1.00±0.00 | 0 | |
| | S4 | 0.99±0.00 | 0.00±0.00 | 1.00±0.00 | 0.00±0.00 | |
| | S5 | 0.01±0.00 | 0.09±0.00 | 0.00±0.00 | 0.11±0.02 | |
| TRAAK | | | | | | |
| | | -150 mV | | 150 mV | | |
| charge | site | K | Water | K | Water | force field |
| 1 | S0 | 0.61±0.07 | 0.87±0.08 | 0.21±0.08 | 1.11±0.06 | Charmm36m |
| | S1 | 0.32±0.05 | 0.21±0.05 | 0.79±0.08 | 0.09±0.08 | |

| | | | | | | |
|---|---|---|---|---|---|---|
| | S2 | 0.85±0.02 | 0.01±0.01 | 0.94±0.01 | 0 | |
| | S3 | 0.63±0.13 | 0.16±0.16 | 0.10±0.01 | 0 | |
| | S4 | 0.41±0.12 | 0.02±0.01 | 0.94±0.02 | 0.03±0.01 | |
| | S5 | 0.58±0.12 | 0.05±0.01 | 0.04±0.01 | 0.07±0.01 | |
| 0.95 | S0 | 0.72±0.02 | 0.73±0.06 | 0.04±0.00 | 1.31±0.02 | Charmm36m |
| | S1 | 0.26±0.02 | 0.12±0.03 | 0.97±0.00 | 0.00±0.00 | |
| | S2 | 0.95±0.01 | 0.00±0.00 | 0.94±0.01 | 0 | |
| | S3 | 0.60±0.16 | 0.25±0.18 | 0.09±0.01 | 0 | |
| | S4 | 0.44±0.15 | 0.01±0.00 | 0.97±0.00 | 0.01±0.00 | |
| | S5 | 0.55±0.14 | 0.04±0.01 | 0.02±0.00 | 0.06±0.00 | |
| 0.9 | S0 | 0.55±0.06 | 0.92±0.08 | 0.06±0.01 | 1.37±0.03 | Charmm36m |
| | S1 | 0.45±0.07 | 0.08±0.03 | 0.97±0.01 | 0.00±0.00 | |
| | S2 | 0.90±0.03 | 0.01±0.01 | 0.90±0.01 | 0 | |
| | S3 | 0.55±0.13 | 0.29±0.18 | 0.12±0.01 | 0 | |
| | S4 | 0.55±0.11 | 0.00±0.00 | 0.97±0.01 | 0.01±0.00 | |
| | S5 | 0.45±0.11 | 0.06±0.01 | 0.02±0.00 | 0.09±0.00 | |
| 0.85 | S0 | 0.69±0.03 | 0.75±0.03 | 0.15±0.02 | 1.35±0.01 | Charmm36m |
| | S1 | 0.35±0.03 | 0.02±0.00 | 0.93±0.01 | 0.00±0.00 | |
| | S2 | 0.87±0.01 | 0 | 0.84±0.01 | 0 | |
| | S3 | 0.83±0.02 | 0 | 0.24±0.02 | 0 | |
| | S4 | 0.48±0.03 | 0.00±0.00 | 0.96±0.00 | 0.01±0.00 | |
| | S5 | 0.52±0.03 | 0.05±0.00 | 0.03±0.00 | 0.11±0.00 | |
| 0.8 | S0 | 0.52±0.08 | 0.92±0.12 | 0.52±0.03 | 0.93±0.05 | Charmm36m |
| | S1 | 0.43±0.01 | 0.11±0.09 | 0.64±0.03 | 0.01±0.00 | |
| | S2 | 0.87±0.05 | 0.00±0.00 | 0.76±0.06 | 0 | |
| | S3 | 0.87±0.04 | 0.05±0.05 | 0.67±0.03 | 0 | |
| | S4 | 0.72±0.04 | 0.00±0.00 | 0.94±0.01 | 0.02±0.00 | |
| | S5 | 0.28±0.04 | 0.07±0.01 | 0.05±0.00 | 0.09±0.01 | |
| 0.78 | S0 | 0.48±0.02 | 0.90±0.02 | 0.54±0.06 | 0.65±0.10 | Charmm36m |
| | S1 | 0.55±0.02 | 0.01±0.00 | 0.53±0.05 | 0.01±0.00 | |
| | S2 | 0.95±0.00 | 0 | 0.93±0.01 | 0 | |
| | S3 | 0.94±0.01 | 0 | 0.90±0.02 | 0 | |
| | S4 | 0.69±0.02 | 0.00±0.00 | 0.94±0.01 | 0.01±0.00 | |
| | S5 | 0.31±0.02 | 0.05±0.00 | 0.05±0.01 | 0.07±0.01 | |

| | | | | | | |
|---|---|---|---|---|---|---|
| 0.77 | S0 | 0.41±0.02 | 0.95±0.02 | 0.50±0.07 | 0.89±0.11 | Charmm36m |
| | S1 | 0.61±0.02 | 0.01±0.00 | 0.59±0.07 | 0.08±0.06 | |
| | S2 | 0.96±0.00 | 0 | 0.69±0.15 | 0.00±0.00 | |
| | S3 | 0.97±0.00 | 0 | 0.93±0.01 | 0 | |
| | S4 | 0.71±0.03 | 0.00±0.00 | 0.95±0.02 | 0.01±0.00 | |
| | S5 | 0.29±0.03 | 0.06±0.01 | 0.04±0.01 | 0.07±0.01 | |
| 0.75 | S0 | 0.25±0.03 | 1.11±0.04 | 0.43±0.02 | 0.92±0.02 | Charmm36m |
| | S1 | 0.77±0.02 | 0.01±0.00 | 0.60±0.01 | 0.01±0.01 | |
| | S2 | 0.99±0.00 | 0 | 0.97±0.00 | 0 | |
| | S3 | 0.99±0.00 | 0 | 0.98±0.00 | 0 | |
| | S4 | 0.81±0.01 | 0.00±0.00 | 0.94±0.01 | 0.01±0.00 | |
| | S5 | 0.19±0.01 | 0.04±0.00 | 0.05±0.01 | 0.06±0.01 | |
| 0.7 | S0 | 0.08±0.01 | 1.37±0.03 | 0.12±0.01 | 1.29±0.03 | Charmm36m |
| | S1 | 0.94±0.04 | 0.04±0.04 | 0.81±0.14 | 0.13±0.13 | |
| | S2 | 1.00±0.00 | 0 | 0.97±0.03 | 0 | |
| | S3 | 1.00±0.00 | 0 | 1.00±0.00 | 0 | |
| | S4 | 0.97±0.00 | 0.00±0.00 | 0.99±0.00 | 0.00±0.00 | |
| | S5 | 0.03±0.00 | 0.05±0.00 | 0.01±0.00 | 0.06±0.00 | |
| 0.65 | S0 | 0.15±0.00 | 1.38±0.02 | 0.13±0.01 | 1.42±0.02 | Charmm36m |
| | S1 | 1.00±0.00 | 0 | 1.00±0.00 | 0 | |
| | S2 | 1.00±0.00 | 0 | 1.00±0.00 | 0 | |
| | S3 | 1.00±0.00 | 0 | 1.00±0.00 | 0 | |
| | S4 | 1.00±0.00 | 0 | 1.00±0.00 | 0 | |
| | S5 | 0.01±0.00 | 0.08±0.00 | 0.00±0.00 | 0.08±0.00 | |
| MthK | | | | | | |
| | | -150 mV | | 150 mV | | |
| charge | site | K | Water | K | Water | force field |
| 1 | S0 | 0.14±0.02 | 1.37±0.03 | 0.36±0.06 | 0.99±0.05 | Charmm36m |
| | S1 | 0.93±0.01 | 0.00±0.00 | 0.66±0.06 | 0.07±0.07 | |
| | S2 | 0.10±0.01 | 0 | 0.79±0.02 | 0 | |
| | S3 | 0.97±0.00 | 0 | 0.22±0.02 | 0 | |
| | S4 | 0.05±0.00 | 0.01±0.00 | 0.83±0.02 | 0.00±0.00 | |
| | S5 | 0.95±0.00 | 0.06±0.01 | 0.17±0.02 | 0.06±0.00 | |
| 0.95 | S0 | 0.44±0.04 | 0.99±0.04 | 0.02±0.00 | 1.36±0.00 | Charmm36m |

| | | | | | | |
|---|---|---|---|---|---|---|
| | S1 | 0.62±0.04 | 0.00±0.00 | 0.99±0.00 | 0 | |
| | S2 | 0.64±0.03 | 0 | 0.91±0.00 | 0 | |
| | S3 | 0.75±0.01 | 0 | 0.10±0.00 | 0 | |
| | S4 | 0.39±0.03 | 0.00±0.00 | 0.99±0.00 | 0.00±0.00 | |
| | S5 | 0.61±0.03 | 0.06±0.01 | 0.01±0.00 | 0.08±0.01 | |
| 0.9 | S0 | 0.43±0.05 | 1.02±0.07 | 0.01±0.00 | 1.46±0.01 | Charmm36m |
| | S1 | 0.60±0.06 | 0.01±0.01 | 1.00±0.00 | 0 | |
| | S2 | 0.78±0.02 | 0 | 0.89±0.01 | 0 | |
| | S3 | 0.62±0.04 | 0 | 0.11±0.01 | 0 | |
| | S4 | 0.58±0.06 | 0.00±0.00 | 1.00±0.00 | 0 | |
| | S5 | 0.42±0.05 | 0.04±0.00 | 0.01±0.00 | 0.08±0.02 | |
| 0.85 | S0 | 0.34±0.02 | 1.19±0.02 | 0.03±0.00 | 1.51±0.01 | Charmm36m |
| | S1 | 0.73±0.02 | 0.00±0.00 | 1.00±0.00 | 0 | |
| | S2 | 0.69±0.01 | 0 | 0.86±0.00 | 0 | |
| | S3 | 0.58±0.01 | 0 | 0.15±0.00 | 0 | |
| | S4 | 0.73±0.02 | 0.00±0.00 | 1.00±0.00 | 0.00±0.00 | |
| | S5 | 0.28±0.02 | 0.07±0.01 | 0.03±0.00 | 0.10±0.01 | |
| 0.8 | S0 | 0.27±0.01 | 1.35±0.02 | 0.11±0.01 | 1.51±0.01 | Charmm36m |
| | S1 | 0.83±0.02 | 0.00±0.00 | 0.99±0.00 | 0 | |
| | S2 | 0.63±0.02 | 0 | 0.78±0.01 | 0 | |
| | S3 | 0.57±0.01 | 0 | 0.23±0.01 | 0 | |
| | S4 | 0.86±0.02 | 0.00±0.00 | 0.99±0.00 | 0.00±0.00 | |
| | S5 | 0.18±0.02 | 0.09±0.01 | 0.07±0.00 | 0.11±0.01 | |
| 0.78 | S0 | | | 0.19±0.00 | 1.42±0.01 | Charmm36m |
| | S1 | | | 0.95±0.00 | 0.00±0.00 | |
| | S2 | | | 0.77±0.01 | 0 | |
| | S3 | | | 0.30±0.00 | 0 | |
| | S4 | | | 0.98±0.00 | 0.00±0.00 | |
| | S5 | | | 0.10±0.01 | 0.13±0.01 | |
| 0.77 | S0 | 0.22±0.02 | 1.40±0.02 | 0.25±0.01 | 1.37±0.01 | Charmm36m |
| | S1 | 0.83±0.02 | 0.00±0.00 | 0.91±0.01 | 0.00±0.00 | |
| | S2 | 0.69±0.01 | 0 | 0.74±0.01 | 0 | |
| | S3 | 0.63±0.03 | 0 | 0.37±0.01 | 0 | |
| | S4 | 0.83±0.02 | 0.00±0.00 | 0.98±0.00 | 0.00±0.00 | |
| | S5 | 0.22±0.01 | 0.10±0.01 | 0.12±0.01 | 0.13±0.01 | |

| | | | | | | |
|---|---|---|---|---|---|---|
| 0.75 | S0 | 0.21±0.02 | 1.40±0.03 | 0.40±0.06 | 1.17±0.06 | Charmm36m |
| | S1 | 0.82±0.02 | 0.00±0.00 | 0.78±0.03 | 0.00±0.00 | |
| | S2 | 0.77±0.01 | 0 | 0.66±0.11 | 0 | |
| | S3 | 0.74±0.03 | 0 | 0.66±0.06 | 0 | |
| | S4 | 0.74±0.01 | 0.00±0.00 | 0.95±0.01 | 0.00±0.00 | |
| | S5 | 0.30±0.01 | 0.09±0.01 | 0.11±0.01 | 0.10±0.01 | |
| 0.7 | S0 | 0.06±0.00 | 1.51±0.01 | 0.15±0.01 | 1.40±0.02 | Charmm36m |
| | S1 | 0.95±0.00 | 0.00±0.00 | 0.87±0.01 | 0.00±0.00 | |
| | S2 | 0.98±0.00 | 0 | 0.99±0.00 | 0 | |
| | S3 | 0.99±0.00 | 0 | 0.99±0.00 | 0 | |
| | S4 | 0.85±0.01 | 0.00±0.00 | 0.96±0.00 | 0.00±0.00 | |
| | S5 | 0.15±0.01 | 0.06±0.01 | 0.04±0.00 | 0.07±0.01 | |
| 0.65 | S0 | 0.03±0.00 | 1.63±0.01 | 0.03±0.00 | 1.60±0.01 | Charmm36m |
| | S1 | 1.00±0.00 | 0 | 0.99±0.00 | 0 | |
| | S2 | 1.00±0.00 | 0 | 1.00±0.00 | 0 | |
| | S3 | 1.00±0.00 | 0 | 1.00±0.00 | 0 | |
| | S4 | 0.99±0.00 | 0.00±0.00 | 0.99±0.00 | 0 | |
| | S5 | 0.02±0.00 | 0.09±0.01 | 0.01±0.00 | 0.09±0.00 | |
| NaK2K | | | | | | |
| | | -150 mV | | 150 mV | | |
| charge | site | K | Water | K | Water | force field |
| 1 | S0 | | | 0.55±0.04 | 0.95±0.06 | Amber14sb |
| | S1 | | | 0.40±0.05 | 0.15±0.02 | |
| | S2 | | | 0.71±0.04 | 0 | |
| | S3 | | | 0.85±0.01 | 0 | |
| | S4 | | | 0.40±0.04 | 0.44±0.05 | |
| | S5 | | | 0.14±0.01 | 0.36±0.02 | |
| 0.95 | S0 | | | 0.39±0.03 | 1.24±0.05 | Amber14sb |
| | S1 | | | 0.60±0.04 | 0.12±0.01 | |
| | S2 | | | 0.59±0.03 | 0 | |
| | S3 | | | 0.81±0.01 | 0 | |
| | S4 | | | 0.64±0.04 | 0.24±0.04 | |
| | S5 | | | 0.11±0.01 | 0.25±0.02 | |
| 0.9 | S0 | | | 0.60±0.04 | 0.92±0.07 | Amber14sb |

| | | | | | | |
|---|---|---|---|---|---|---|
| | S1 | | | 0.35±0.04 | 0.15±0.02 | |
| | S2 | | | 0.74±0.03 | 0 | |
| | S3 | | | 0.91±0.01 | 0 | |
| | S4 | | | 0.58±0.06 | 0.35±0.06 | |
| | S5 | | | 0.11±0.01 | 0.27±0.04 | |
| 0.85 | S0 | | | 0.79±0.04 | 0.66±0.05 | Amber14sb |
| | S1 | | | 0.09±0.01 | 0.23±0.02 | |
| | S2 | | | 0.93±0.01 | 0 | |
| | S3 | | | 0.98±0.00 | 0 | |
| | S4 | | | 0.51±0.07 | 0.43±0.05 | |
| | S5 | | | 0.10±0.01 | 0.32±0.07 | |
| 0.8 | S0 | | | 0.80±0.04 | 0.67±0.05 | Amber14sb |
| | S1 | | | 0.12±0.05 | 0.14±0.02 | |
| | S2 | | | 0.98±0.00 | 0 | |
| | S3 | | | 0.99±0.00 | 0 | |
| | S4 | | | 0.78±0.06 | 0.21±0.06 | |
| | S5 | | | 0.05±0.01 | 0.23±0.05 | |
| 0.75 | S0 | | | 0.61±0.01 | 0.89±0.08 | Amber14sb |
| | S1 | | | 0.34±0.01 | 0.17±0.02 | |
| | S2 | | | 1.00±0.00 | 0 | |
| | S3 | | | 1.00±0.00 | 0 | |
| | S4 | | | 0.94±0.01 | 0.06±0.01 | |
| | S5 | | | 0.01±0.00 | 0.16±0.01 | |
| 0.7 | S0 | | | 0.11±0.01 | 1.61±0.01 | Amber14sb |
| | S1 | | | 0.89±0.01 | 0.04±0.01 | |
| | S2 | | | 1.00±0.00 | 0 | |
| | S3 | | | 1.00±0.00 | 0 | |
| | S4 | | | 1.00±0.00 | 0 | |
| | S5 | | | 0.00±0.00 | 0.12±0.01 | |
| 0.65 | S0 | | | 0.03±0.00 | 1.72±0.05 | Amber14sb |
| | S1 | | | 1.00±0.00 | 0 | |
| | S2 | | | 1.00±0.00 | 0 | |
| | S3 | | | 1.00±0.00 | 0 | |
| | S4 | | | 1.00±0.00 | 0 | |
| | S5 | | | 0.00±0.00 | 0.18±0.00 | |

| | | | | | | |
|---|---|---|---|---|---|---|
| TRAAK | | | | | | |
| | | -150 mV | | 150 mV | | |
| charge | site | K | Water | K | Water | force field |
| 1 | S0 | | | 0.51±0.09 | 0.80±0.10 | Amber14sb |
| | S1 | | | 0.42±0.08 | 0.15±0.04 | |
| | S2 | | | 0.90±0.02 | 0 | |
| | S3 | | | 0.63±0.06 | 0 | |
| | S4 | | | 0.48±0.08 | 0.06±0.01 | |
| | S5 | | | 0.46±0.08 | 0.10±0.01 | |
| 0.95 | S0 | | | 0.37±0.06 | 1.00±0.06 | Amber14sb |
| | S1 | | | 0.60±0.06 | 0.07±0.01 | |
| | S2 | | | 0.78±0.03 | 0 | |
| | S3 | | | 0.62±0.05 | 0 | |
| | S4 | | | 0.64±0.05 | 0.05±0.01 | |
| | S5 | | | 0.32±0.04 | 0.11±0.01 | |
| 0.9 | S0 | | | 0.56±0.05 | 0.80±0.08 | Amber14sb |
| | S1 | | | 0.42±0.04 | 0.11±0.02 | |
| | S2 | | | 0.79±0.02 | 0 | |
| | S3 | | | 0.80±0.03 | 0 | |
| | S4 | | | 0.79±0.02 | 0.04±0.01 | |
| | S5 | | | 0.18±0.02 | 0.12±0.01 | |
| 0.85 | S0 | | | 0.89±0.02 | 0.37±0.04 | Amber14sb |
| | S1 | | | 0.08±0.01 | 0.10±0.02 | |
| | S2 | | | 0.94±0.02 | 0 | |
| | S3 | | | 0.98±0.00 | 0 | |
| | S4 | | | 0.94±0.01 | 0.03±0.01 | |
| | S5 | | | 0.04±0.00 | 0.15±0.02 | |
| 0.8 | S0 | | | 0.71±0.12 | 0.91±0.34 | Amber14sb |
| | S1 | | | 0.09±0.02 | 0.28±0.10 | |
| | S2 | | | 0.99±0.00 | 0 | |
| | S3 | | | 1.00±0.00 | 0 | |
| | S4 | | | 0.97±0.01 | 0.02±0.01 | |
| | S5 | | | 0.01±0.00 | 0.15±0.02 | |
| 0.75 | S0 | | | 0.39±0.03 | 0.98±0.17 | Amber14sb |

| | | | | | | |
|---|---|---|---|---|---|---|
| | S1 | | | 0.57±0.03 | 0.09±0.04 | |
| | S2 | | | 1.00±0.00 | 0 | |
| | S3 | | | 1.00±0.00 | 0 | |
| | S4 | | | 1.00±0.00 | 0.00±0.00 | |
| | S5 | | | 0.00±0.00 | 0.15±0.02 | |
| 0.7 | S0 | | | 0.08±0.01 | 1.44±0.13 | Amber14sb |
| | S1 | | | 0.97±0.00 | 0.01±0.01 | |
| | S2 | | | 1.00±0.00 | 0 | |
| | S3 | | | 1.00±0.00 | 0 | |
| | S4 | | | 1.00±0.00 | 0.00±0.00 | |
| | S5 | | | 0.00±0.00 | 0.18±0.03 | |
| 0.65 | S0 | | | 0.19±0.07 | 1.30±0.09 | Amber14sb |
| | S1 | | | 0.99±0.01 | 0.01±0.00 | |
| | S2 | | | 1.00±0.00 | 0 | |
| | S3 | | | 1.00±0.00 | 0 | |
| | S4 | | | 1.00±0.00 | 0 | |
| | S5 | | | 0.01±0.00 | 0.17±0.01 | |
| MthK | | | | | | |
| | | -150 mV | | 150 mV | | |
| charge | site | K | Water | K | Water | force field |
| 1 | S0 | | | 0.64±0.07 | 0.77±0.10 | Amber14sb |
| | S1 | | | 0.26±0.04 | 0.14±0.04 | |
| | S2 | | | 0.87±0.02 | 0 | |
| | S3 | | | 0.83±0.03 | 0 | |
| | S4 | | | 0.27±0.05 | 0.06±0.02 | |
| | S5 | | | 0.70±0.05 | 0.06±0.01 | |
| 0.95 | S0 | | | 0.53±0.05 | 0.92±0.06 | Amber14sb |
| | S1 | | | 0.47±0.04 | 0.04±0.01 | |
| | S2 | | | 0.82±0.02 | 0 | |
| | S3 | | | 0.71±0.03 | 0 | |
| | S4 | | | 0.47±0.04 | 0.03±0.01 | |
| | S5 | | | 0.50±0.04 | 0.08±0.01 | |
| 0.9 | S0 | | | 0.45±0.03 | 1.10±0.05 | Amber14sb |
| | S1 | | | 0.59±0.03 | 0.06±0.01 | |

| | | | | | | |
|---|---|---|---|---|---|---|
| | S2 | | | 0.70±0.03 | 0 | |
| | S3 | | | 0.71±0.01 | 0 | |
| | S4 | | | 0.68±0.02 | 0.04±0.01 | |
| | S5 | | | 0.29±0.01 | 0.08±0.02 | |
| 0.85 | S0 | | | 0.65±0.03 | 0.82±0.05 | Amber14sb |
| | S1 | | | 0.31±0.02 | 0.12±0.01 | |
| | S2 | | | 0.79±0.01 | 0 | |
| | S3 | | | 0.90±0.01 | 0 | |
| | S4 | | | 0.90±0.01 | 0.01±0.01 | |
| | S5 | | | 0.09±0.01 | 0.11±0.04 | |
| 0.8 | S0 | | | 0.86±0.03 | 0.57±0.04 | Amber14sb |
| | S1 | | | 0.05±0.00 | 0.11±0.02 | |
| | S2 | | | 0.97±0.00 | 0 | |
| | S3 | | | 0.99±0.00 | 0 | |
| | S4 | | | 0.96±0.01 | 0.03±0.01 | |
| | S5 | | | 0.02±0.00 | 0.08±0.01 | |
| 0.75 | S0 | | | 0.58±0.11 | 0.94±0.12 | Amber14sb |
| | S1 | | | 0.22±0.05 | 0.22±0.12 | |
| | S2 | | | 1.00±0.00 | 0 | |
| | S3 | | | 1.00±0.00 | 0 | |
| | S4 | | | 1.00±0.00 | 0.00±0.00 | |
| | S5 | | | 0.00±0.00 | 0.08±0.01 | |
| 0.7 | S0 | | | 0.11±0.01 | 1.53±0.01 | Amber14sb |
| | S1 | | | 0.90±0.01 | 0.01±0.00 | |
| | S2 | | | 1.00±0.00 | 0 | |
| | S3 | | | 1.00±0.00 | 0 | |
| | S4 | | | 1.00±0.00 | 0 | |
| | S5 | | | 0.00±0.00 | 0.08±0.01 | |
| 0.65 | S0 | | | 0.03±0.00 | 1.69±0.01 | Amber14sb |
| | S1 | | | 0.99±0.00 | 0 | |
| | S2 | | | 1.00±0.00 | 0 | |
| | S3 | | | 1.00±0.00 | 0 | |
| | S4 | | | 1.00±0.00 | 0 | |
| | S5 | | | 0.01±0.00 | 0.11±0.01 | |

| NaK2K | | | | | | |
|---|---|---|---|---|---|---|
| | | -150 mV | | 150 mV | | |
| charge | site | K | Water | K | Water | force field |
| 1 | S0 | 0.79±0.13 | 0.69±0.21 | 0.78±0.09 | 0.66±0.14 | Amber14sb-S3 |
| | S1 | 0.22±0.15 | 0.77±0.15 | 0.27±0.11 | 0.54±0.19 | |
| | S2 | 0.80±0.14 | 0 | 0.78±0.08 | 0 | |
| | S3 | 0.25±0.17 | 0 | 0.24±0.09 | 0 | |
| | S4 | 0.77±0.15 | 0.02±0.01 | 0.76±0.09 | 0.21±0.09 | |
| | S5 | 0.22±0.15 | 0.18±0.04 | 0.08±0.04 | 0.27±0.03 | |
| 0.95 | S0 | 0.86±0.05 | 0.59±0.08 | 0.54±0.12 | 1.00±0.15 | Amber14sb-S3 |
| | S1 | 0.12±0.05 | 0.83±0.06 | 0.47±0.12 | 0.46±0.12 | |
| | S2 | 0.93±0.03 | 0 | 0.90±0.02 | 0 | |
| | S3 | 0.17±0.05 | 0 | 0.16±0.04 | 0 | |
| | S4 | 0.86±0.04 | 0.03±0.01 | 0.86±0.04 | 0.11±0.02 | |
| | S5 | 0.11±0.04 | 0.17±0.02 | 0.03±0.01 | 0.24±0.05 | |
| 0.9 | S0 | 0.52±0.09 | 1.10±0.13 | 0.11±0.04 | 1.63±0.06 | Amber14sb-S3 |
| | S1 | 0.47±0.10 | 0.38±0.10 | 0.92±0.04 | 0.04±0.03 | |
| | S2 | 0.86±0.02 | 0.00±0.00 | 0.92±0.00 | 0 | |
| | S3 | 0.34±0.05 | 0 | 0.12±0.02 | 0 | |
| | S4 | 0.78±0.04 | 0.04±0.01 | 0.94±0.03 | 0.05±0.02 | |
| | S5 | 0.18±0.03 | 0.20±0.01 | 0.02±0.00 | 0.18±0.01 | |
| 0.85 | S0 | 0.44±0.02 | 1.20±0.04 | 0.14±0.03 | 1.63±0.05 | Amber14sb-S3 |
| | S1 | 0.57±0.02 | 0.19±0.02 | 0.90±0.03 | 0.02±0.01 | |
| | S2 | 0.78±0.01 | 0.00±0.00 | 0.94±0.01 | 0.00±0.00 | |
| | S3 | 0.57±0.04 | 0.00±0.00 | 0.16±0.04 | 0.15±0.15 | |
| | S4 | 0.70±0.03 | 0.05±0.00 | 0.90±0.03 | 0.09±0.02 | |
| | S5 | 0.25±0.03 | 0.17±0.02 | 0.01±0.00 | 0.27±0.02 | |
| 0.8 | S0 | 0.56±0.01 | 1.08±0.02 | 0.27±0.02 | 1.50±0.02 | Amber14sb-S3 |
| | S1 | 0.41±0.02 | 0.22±0.02 | 0.80±0.02 | 0.04±0.01 | |
| | S2 | 0.79±0.01 | 0 | 0.86±0.02 | 0 | |
| | S3 | 0.83±0.01 | 0 | 0.38±0.01 | 0 | |
| | S4 | 0.72±0.01 | 0.04±0.01 | 0.79±0.02 | 0.18±0.02 | |
| | S5 | 0.23±0.01 | 0.15±0.01 | 0.03±0.00 | 0.29±0.06 | |
| 0.78 | S0 | 0.54±0.01 | 1.11±0.02 | 0.37±0.03 | 1.32±0.04 | Amber14sb-S3 |
| | S1 | 0.37±0.01 | 0.20±0.01 | 0.67±0.03 | 0.07±0.01 | |

| | | | | | | |
|---|---|---|---|---|---|---|
| | S2 | 0.82±0.01 | 0 | 0.87±0.01 | 0.00±0.00 | |
| | S3 | 0.89±0.01 | 0 | 0.56±0.02 | 0 | |
| | S4 | 0.79±0.01 | 0.02±0.01 | 0.69±0.02 | 0.28±0.03 | |
| | S5 | 0.19±0.01 | 0.12±0.01 | 0.05±0.01 | 0.29±0.01 | |
| 0.77 | S0 | 0.51±0.02 | 1.17±0.03 | 0.29±0.05 | 1.42±0.07 | Amber14sb-S3 |
| | S1 | 0.40±0.02 | 0.22±0.02 | 0.75±0.05 | 0.04±0.01 | |
| | S2 | 0.87±0.01 | 0 | 0.93±0.02 | 0 | |
| | S3 | 0.87±0.01 | 0 | 0.64±0.03 | 0 | |
| | S4 | 0.78±0.01 | 0.05±0.01 | 0.51±0.05 | 0.43±0.06 | |
| | S5 | 0.17±0.01 | 0.15±0.01 | 0.06±0.01 | 0.59±0.13 | |
| 0.75 | S0 | 0.53±0.01 | 1.12±0.03 | 0.44±0.01 | 1.23±0.01 | Amber14sb-S3 |
| | S1 | 0.36±0.02 | 0.22±0.02 | 0.58±0.01 | 0.06±0.01 | |
| | S2 | 0.92±0.01 | 0 | 0.91±0.01 | 0 | |
| | S3 | 0.96±0.01 | 0 | 0.84±0.01 | 0 | |
| | S4 | 0.80±0.02 | 0.03±0.01 | 0.55±0.02 | 0.37±0.03 | |
| | S5 | 0.16±0.00 | 0.18±0.04 | 0.06±0.01 | 0.33±0.02 | |
| 0.7 | S0 | 0.14±0.01 | 1.49±0.12 | 0.22±0.02 | 1.48±0.02 | Amber14sb-S3 |
| | S1 | 0.84±0.01 | 0.07±0.01 | 0.79±0.02 | 0.03±0.01 | |
| | S2 | 1.00±0.00 | 0 | 0.99±0.00 | 0 | |
| | S3 | 1.00±0.00 | 0 | 0.99±0.00 | 0 | |
| | S4 | 0.84±0.04 | 0.08±0.03 | 0.74±0.03 | 0.21±0.02 | |
| | S5 | 0.09±0.01 | 0.16±0.02 | 0.05±0.01 | 0.23±0.01 | |
| 0.65 | S0 | 0.04±0.00 | 1.76±0.02 | 0.04±0.00 | 1.73±0.02 | Amber14sb-S3 |
| | S1 | 0.99±0.01 | 0.01±0.00 | 0.98±0.00 | 0.00±0.00 | |
| | S2 | 1.00±0.00 | 0 | 1.00±0.00 | 0 | |
| | S3 | 1.00±0.00 | 0 | 1.00±0.00 | 0 | |
| | S4 | 0.96±0.01 | 0.02±0.01 | 0.97±0.01 | 0.02±0.01 | |
| | S5 | 0.02±0.00 | 0.18±0.02 | 0.01±0.00 | 0.16±0.03 | |
| TRAAK | | | | | | |
| | | -150 mV | | 150 mV | | |
| charge | site | K | Water | K | Water | force field |
| 1 | S0 | 0.87±0.06 | 0.46±0.08 | 0.78±0.13 | 0.49±0.13 | Amber14sb-S3 |
| | S1 | 0.09±0.04 | 0.84±0.07 | 0.22±0.13 | 0.63±0.18 | |
| | S2 | 0.94±0.03 | 0 | 0.99±0.01 | 0 | |

| | | | | | | |
|---|---|---|---|---|---|---|
| | S3 | 0.20±0.10 | 0 | 0.03±0.02 | 0 | |
| | S4 | 0.80±0.10 | 0.01±0.00 | 0.97±0.02 | 0.02±0.01 | |
| | S5 | 0.19±0.10 | 0.12±0.01 | 0.01±0.01 | 0.10±0.01 | |
| 0.95 | S0 | 0.88±0.06 | 0.46±0.06 | 0.03±0.00 | 1.36±0.04 | Amber14sb-S3 |
| | S1 | 0.07±0.04 | 0.89±0.07 | 0.97±0.00 | 0.00±0.00 | |
| | S2 | 0.98±0.01 | 0 | 0.99±0.00 | 0 | |
| | S3 | 0.12±0.05 | 0 | 0.02±0.00 | 0 | |
| | S4 | 0.89±0.05 | 0.00±0.00 | 0.99±0.00 | 0.00±0.00 | |
| | S5 | 0.11±0.05 | 0.11±0.01 | 0.01±0.00 | 0.09±0.01 | |
| 0.9 | S0 | 0.77±0.02 | 0.58±0.03 | 0.03±0.01 | 1.33±0.02 | Amber14sb-S3 |
| | S1 | 0.22±0.02 | 0.45±0.11 | 0.99±0.00 | 0.00±0.00 | |
| | S2 | 0.98±0.00 | 0.00±0.00 | 0.98±0.00 | 0 | |
| | S3 | 0.41±0.10 | 0.08±0.08 | 0.02±0.00 | 0 | |
| | S4 | 0.61±0.10 | 0.03±0.01 | 0.99±0.00 | 0.00±0.00 | |
| | S5 | 0.35±0.09 | 0.13±0.02 | 0.01±0.00 | 0.12±0.01 | |
| 0.85 | S0 | 0.44±0.10 | 0.97±0.11 | 0.11±0.02 | 1.33±0.05 | Amber14sb-S3 |
| | S1 | 0.57±0.11 | 0.17±0.08 | 0.98±0.00 | 0.00±0.00 | |
| | S2 | 0.87±0.06 | 0.07±0.05 | 0.96±0.00 | 0 | |
| | S3 | 0.45±0.11 | 0.30±0.17 | 0.06±0.01 | 0 | |
| | S4 | 0.69±0.10 | 0.01±0.00 | 0.98±0.01 | 0.01±0.00 | |
| | S5 | 0.30±0.10 | 0.16±0.03 | 0.02±0.00 | 0.15±0.02 | |
| 0.8 | S0 | 0.53±0.04 | 0.86±0.04 | 0.34±0.01 | 1.10±0.02 | Amber14sb-S3 |
| | S1 | 0.50±0.03 | 0.11±0.01 | 0.87±0.01 | 0.01±0.00 | |
| | S2 | 0.90±0.01 | 0 | 0.90±0.01 | 0 | |
| | S3 | 0.70±0.03 | 0 | 0.25±0.02 | 0 | |
| | S4 | 0.64±0.02 | 0.01±0.00 | 0.93±0.01 | 0.04±0.00 | |
| | S5 | 0.36±0.02 | 0.12±0.01 | 0.05±0.00 | 0.21±0.03 | |
| 0.78 | S0 | 0.47±0.04 | 0.97±0.04 | 0.49±0.01 | 0.90±0.03 | Amber14sb-S3 |
| | S1 | 0.52±0.04 | 0.14±0.01 | 0.74±0.02 | 0.03±0.00 | |
| | S2 | 0.82±0.07 | 0.06±0.06 | 0.89±0.01 | 0 | |
| | S3 | 0.78±0.03 | 0 | 0.44±0.02 | 0 | |
| | S4 | 0.73±0.03 | 0.01±0.00 | 0.89±0.01 | 0.07±0.01 | |
| | S5 | 0.27±0.03 | 0.19±0.04 | 0.07±0.00 | 0.23±0.04 | |
| 0.77 | S0 | 0.44±0.03 | 1.08±0.11 | 0.59±0.02 | 0.77±0.03 | Amber14sb-S3 |
| | S1 | 0.51±0.09 | 0.27±0.06 | 0.64±0.02 | 0.03±0.00 | |

| | | | | | | |
|---|---|---|---|---|---|---|
| | S2 | 0.71±0.14 | 0.21±0.13 | 0.88±0.03 | 0 | |
| | S3 | 0.84±0.05 | 0.07±0.05 | 0.58±0.03 | 0 | |
| | S4 | 0.81±0.06 | 0.01±0.00 | 0.89±0.01 | 0.07±0.01 | |
| | S5 | 0.19±0.06 | 0.17±0.02 | 0.07±0.01 | 0.16±0.01 | |
| 0.75 | S0 | 0.40±0.02 | 1.00±0.02 | 0.59±0.06 | 0.86±0.18 | Amber14sb-S3 |
| | S1 | 0.61±0.02 | 0.09±0.02 | 0.44±0.05 | 0.13±0.10 | |
| | S2 | 0.97±0.00 | 0 | 0.93±0.01 | 0 | |
| | S3 | 0.95±0.01 | 0 | 0.85±0.03 | 0 | |
| | S4 | 0.66±0.02 | 0.02±0.00 | 0.89±0.02 | 0.05±0.01 | |
| | S5 | 0.33±0.02 | 0.14±0.01 | 0.07±0.01 | 0.14±0.01 | |
| 0.7 | S0 | 0.14±0.02 | 1.42±0.15 | 0.20±0.03 | 1.06±0.13 | Amber14sb-S3 |
| | S1 | 0.90±0.01 | 0.03±0.02 | 0.86±0.02 | 0.02±0.01 | |
| | S2 | 1.00±0.00 | 0 | 0.99±0.01 | 0 | |
| | S3 | 1.00±0.00 | 0 | 1.00±0.00 | 0 | |
| | S4 | 0.90±0.01 | 0.01±0.00 | 0.97±0.00 | 0.01±0.00 | |
| | S5 | 0.09±0.01 | 0.15±0.02 | 0.02±0.00 | 0.15±0.02 | |
| 0.65 | S0 | 0.13±0.01 | 1.39±0.03 | 0.16±0.03 | 1.31±0.05 | Amber14sb-S3 |
| | S1 | 0.99±0.00 | 0.00±0.00 | 0.99±0.00 | 0.00±0.00 | |
| | S2 | 1.00±0.00 | 0 | 1.00±0.00 | 0 | |
| | S3 | 1.00±0.00 | 0 | 1.00±0.00 | 0 | |
| | S4 | 0.99±0.00 | 0.00±0.00 | 1.00±0.00 | 0 | |
| | S5 | 0.01±0.00 | 0.19±0.01 | 0.00±0.00 | 0.18±0.02 | |
| MthK | | | | | | |
| | | -150 mV | | 150 mV | | |
| charge | site | K | Water | K | Water | force field |
| 1 | S0 | 0.36±0.02 | 1.32±0.03 | 0.83±0.03 | 0.48±0.04 | Amber14sb-S3 |
| | S1 | 0.89±0.01 | 0.06±0.01 | 0.19±0.03 | 0.12±0.11 | |
| | S2 | 0.12±0.01 | 0 | 0.88±0.03 | 0 | |
| | S3 | 0.97±0.01 | 0 | 0.12±0.03 | 0 | |
| | S4 | 0.04±0.01 | 0.02±0.00 | 0.88±0.03 | 0.00±0.00 | |
| | S5 | 0.95±0.01 | 0.03±0.01 | 0.12±0.03 | 0.09±0.02 | |
| 0.95 | S0 | 0.60±0.07 | 0.90±0.11 | 0.19±0.03 | 1.32±0.04 | Amber14sb-S3 |
| | S1 | 0.41±0.06 | 0.37±0.10 | 0.82±0.03 | 0.05±0.03 | |
| | S2 | 0.68±0.05 | 0 | 0.94±0.01 | 0 | |

| | | | | | | |
|---|---|---|---|---|---|---|
| | S3 | 0.69±0.13 | 0 | 0.08±0.01 | 0 | |
| | S4 | 0.32±0.13 | 0.02±0.00 | 0.93±0.01 | 0.01±0.01 | |
| | S5 | 0.66±0.14 | 0.04±0.02 | 0.06±0.00 | 0.11±0.03 | |
| 0.9 | S0 | 0.76±0.08 | 0.67±0.09 | 0.03±0.00 | 1.61±0.01 | Amber14sb-S3 |
| | S1 | 0.22±0.08 | 0.43±0.15 | 0.99±0.00 | 0.00±0.00 | |
| | S2 | 0.96±0.01 | 0 | 0.98±0.00 | 0 | |
| | S3 | 0.43±0.11 | 0 | 0.03±0.00 | 0 | |
| | S4 | 0.59±0.10 | 0.01±0.00 | 0.99±0.00 | 0 | |
| | S5 | 0.40±0.10 | 0.11±0.04 | 0.02±0.00 | 0.07±0.00 | |
| 0.85 | S0 | 0.64±0.05 | 0.85±0.05 | 0.05±0.00 | 1.64±0.01 | Amber14sb-S3 |
| | S1 | 0.37±0.05 | 0.16±0.09 | 0.99±0.00 | 0.00±0.00 | |
| | S2 | 0.94±0.01 | 0 | 0.97±0.00 | 0 | |
| | S3 | 0.59±0.05 | 0 | 0.04±0.01 | 0 | |
| | S4 | 0.48±0.05 | 0.01±0.00 | 0.99±0.00 | 0.00±0.00 | |
| | S5 | 0.51±0.04 | 0.07±0.01 | 0.02±0.00 | 0.11±0.02 | |
| 0.8 | S0 | 0.49±0.03 | 1.12±0.05 | 0.10±0.01 | 1.65±0.01 | Amber14sb-S3 |
| | S1 | 0.52±0.03 | 0.14±0.01 | 0.98±0.00 | 0.00±0.00 | |
| | S2 | 0.88±0.01 | 0 | 0.93±0.01 | 0 | |
| | S3 | 0.63±0.03 | 0 | 0.09±0.01 | 0 | |
| | S4 | 0.56±0.04 | 0.01±0.00 | 0.98±0.00 | 0.01±0.00 | |
| | S5 | 0.44±0.04 | 0.06±0.01 | 0.05±0.00 | 0.11±0.01 | |
| 0.78 | S0 | 0.40±0.04 | 1.26±0.06 | 0.19±0.01 | 1.55±0.02 | Amber14sb-S3 |
| | S1 | 0.59±0.05 | 0.11±0.00 | 0.93±0.00 | 0.01±0.00 | |
| | S2 | 0.77±0.08 | 0.09±0.09 | 0.90±0.01 | 0 | |
| | S3 | 0.74±0.03 | 0.00±0.00 | 0.18±0.00 | 0 | |
| | S4 | 0.63±0.04 | 0.01±0.00 | 0.96±0.01 | 0.02±0.01 | |
| | S5 | 0.38±0.04 | 0.06±0.01 | 0.08±0.00 | 0.12±0.01 | |
| 0.77 | S0 | 0.38±0.01 | 1.27±0.02 | 0.22±0.01 | 1.50±0.02 | Amber14sb-S3 |
| | S1 | 0.57±0.01 | 0.12±0.01 | 0.90±0.01 | 0.02±0.00 | |
| | S2 | 0.86±0.01 | 0 | 0.90±0.01 | 0 | |
| | S3 | 0.74±0.02 | 0 | 0.22±0.01 | 0 | |
| | S4 | 0.62±0.03 | 0.01±0.00 | 0.96±0.01 | 0.01±0.00 | |
| | S5 | 0.39±0.03 | 0.09±0.02 | 0.08±0.01 | 0.18±0.05 | |
| 0.75 | S0 | 0.28±0.05 | 1.40±0.08 | 0.37±0.01 | 1.30±0.02 | Amber14sb-S3 |
| | S1 | 0.66±0.07 | 0.11±0.03 | 0.73±0.01 | 0.03±0.01 | |

| | | | | | | |
|---|---|---|---|---|---|---|
| | S2 | 0.91±0.01 | 0.01±0.01 | 0.89±0.02 | 0 | |
| | S3 | 0.70±0.14 | 0.17±0.17 | 0.48±0.01 | 0 | |
| | S4 | 0.68±0.08 | 0.01±0.00 | 0.87±0.04 | 0.08±0.04 | |
| | S5 | 0.33±0.08 | 0.09±0.03 | 0.10±0.01 | 0.14±0.02 | |
| 0.7 | S0 | 0.16±0.01 | 1.52±0.01 | 0.33±0.02 | 1.29±0.03 | Amber14sb-S3 |
| | S1 | 0.81±0.01 | 0.06±0.01 | 0.68±0.03 | 0.03±0.00 | |
| | S2 | 0.99±0.00 | 0 | 0.97±0.01 | 0 | |
| | S3 | 1.00±0.00 | 0 | 0.98±0.00 | 0 | |
| | S4 | 0.81±0.01 | 0.02±0.00 | 0.92±0.01 | 0.03±0.01 | |
| | S5 | 0.17±0.01 | 0.09±0.00 | 0.06±0.00 | 0.14±0.04 | |
| 0.65 | S0 | 0.04±0.00 | 1.71±0.00 | 0.06±0.01 | 1.66±0.01 | Amber14sb-S3 |
| | S1 | 0.98±0.00 | 0.00±0.00 | 0.96±0.01 | 0.00±0.00 | |
| | S2 | 1.00±0.00 | 0 | 1.00±0.00 | 0 | |
| | S3 | 1.00±0.00 | 0 | 1.00±0.00 | 0 | |
| | S4 | 0.98±0.00 | 0.00±0.00 | 0.99±0.00 | 0.00±0.00 | |
| | S5 | 0.02±0.00 | 0.08±0.01 | 0.01±0.00 | 0.09±0.02 | |

Table S3: $K^+$ ion and water occupancy in SF binding site in NaK2K, TRAAK, and MthK channels with Charmm36m, Amber14sb, and Amber14sb-S3 force fields, at different scaling factors.

| +150 | | | | | | -150 | | | | | |
|---|---|---|---|---|---|---|---|---|---|---|---|
| **i_name** | **j_name** | **flux_ij** | **flux_ji** | rate_ij | rate_ji | **i_name** | **j_name** | **flux_ij** | **flux_ji** | rate_ij | rate_ji |
| WKK0KK | WKKK0K | 454 | 62 | 0.318 | 9.815 | WKKK0K | WKK0KK | 1023 | 197 | 0.912 | 1.084 |
| WKK0KK | 0KKK0K | 15 | 1 | 0.220 | 6.683 | K0KKK0 | W0KKK0 | 164 | 185 | 1.035 | 0.566 |
| 0KK0KK | WKKK0K | 15 | 0 | 0.307 | 2.481 | K0KKK0 | WK0KK0 | 31 | 19 | 1.216 | 0.596 |
| WKK0K0 | WKKK0K | 12 | 2 | 0.299 | 16.923 | 0KKK0K | WKK0KK | 26 | 8 | 0.923 | 1.080 |
| WKK0KW | WKKK0K | 10 | 4 | 0.301 | 12.118 | WKKK0K | 0KK0KK | 25 | 5 | 0.629 | 0.531 |
| W0KKK0 | K0KKK0 | 9 | 9 | 0.464 | 0.311 | WKKK0K | WKK0K0 | 21 | 2 | 0.982 | 0.525 |
| WK0KK0 | K0KKK0 | 7 | 4 | 0.614 | 2.444 | K0KKKW | W0KKK0 | 15 | 10 | 0.659 | 0.465 |
| W0KKKW | K0KKK0 | 3 | 1 | 0.463 | 0.132 | K0KKKW | W0KKKW | 11 | 11 | 0.417 | 0.472 |
| 0KK0KK | 0KKK0K | 1 | 0 | 0.221 | 1.792 | K0KKK0 | W0KKKW | 11 | 12 | 0.454 | 0.516 |
| 0KK0K0 | 0KKK0K | 1 | 0 | 0.209 | 6.588 | WKKK0K | WKK0KW | 11 | 4 | 0.885 | 0.479 |
| WK0KK0 | K0KKKW | 1 | 0 | 0.543 | 2.395 | 0KKK0K | 0KK0KK | 7 | 0 | 0.635 | 0.512 |
| 0KK0K0 | WKKK0K | 1 | 0 | 0.294 | 8.114 | K0KK0K | W0KKK0 | 4 | 0 | 0.594 | 0.446 |
| | | | | | | KKKK0K | WKK0KK | 2 | 0 | 0.944 | 0.337 |
| | | | | | | K0KKK0 | 0K0KK0 | 1 | 0 | 0.852 | 0.501 |
| | | | | | | K0KKKW | WK0KKW | 1 | 1 | 0.877 | 0.478 |
| | | | | | | K0KKKW | WK0KK0 | 1 | 3 | 0.989 | 0.532 |
| | | | | | | 0KKK0K | WKK0K0 | 1 | 0 | 0.999 | 0.515 |

Table S4: Flux and rate between states in the rate-limiting step in MthK -150 mV and +150 mV simulations. State pairs that have no flux observed are not listed. The rate is computed as inversed mean first passage time, unit in ns$^{-1}$.

### 3. Ion Reparameterization (Radius corrected vdW)

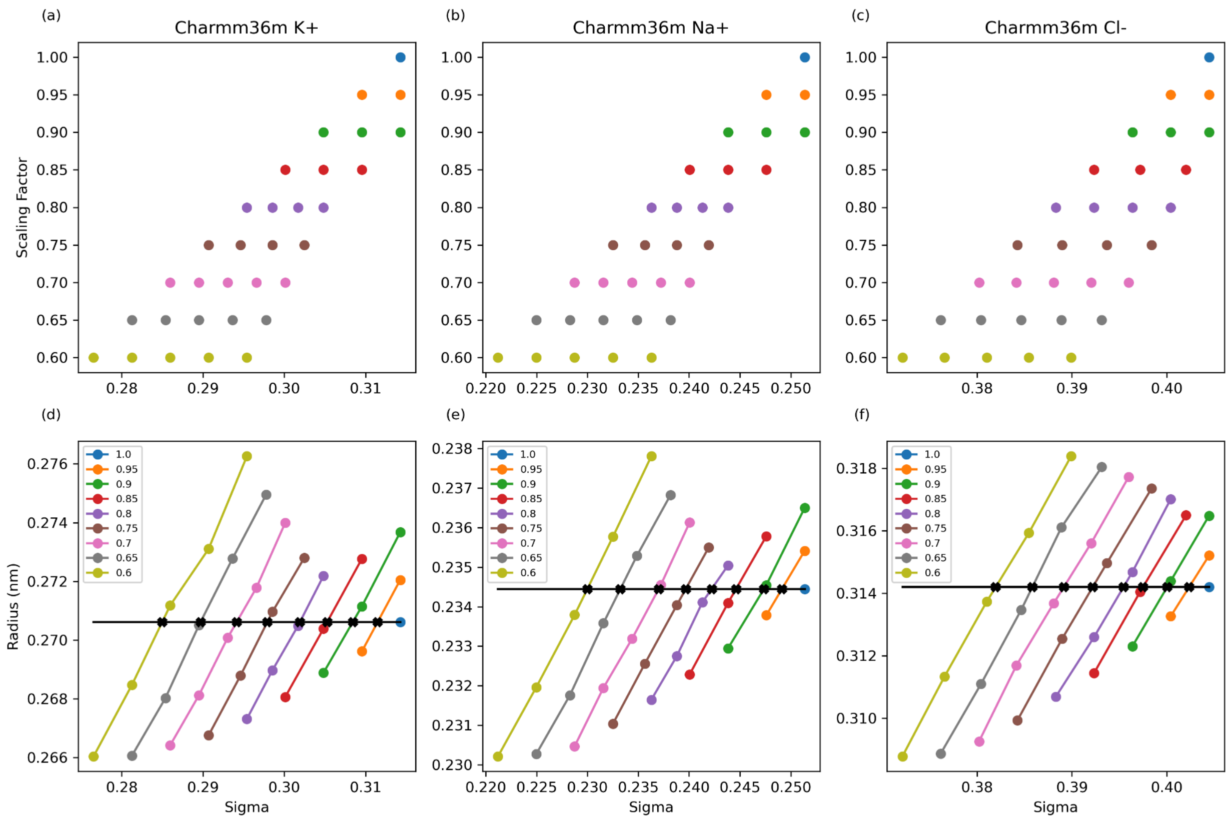

Fig. S8: Ion parameter grid search in Charmm36m. (a)(b)(c) Search grids with (sigma, scaling factor) pairs for $K^+$, $Na^+$, $Cl^-$ ions. (d)(e)(f) The radius of all the points on the grids. The radius from the original radius at scaling factor 1.00 was chosen as the targeted radius. The optimized sigma values for every scaling factor point at a step of 0.05 (1.00, 0.95, 0.90, …) are plotted as x. The sigma parameters for the scaling factor between 0.05 (0.99~0.96, 0.94~0.91, …) are interpolated. All the optimized ion parameters are listed in Table S5.

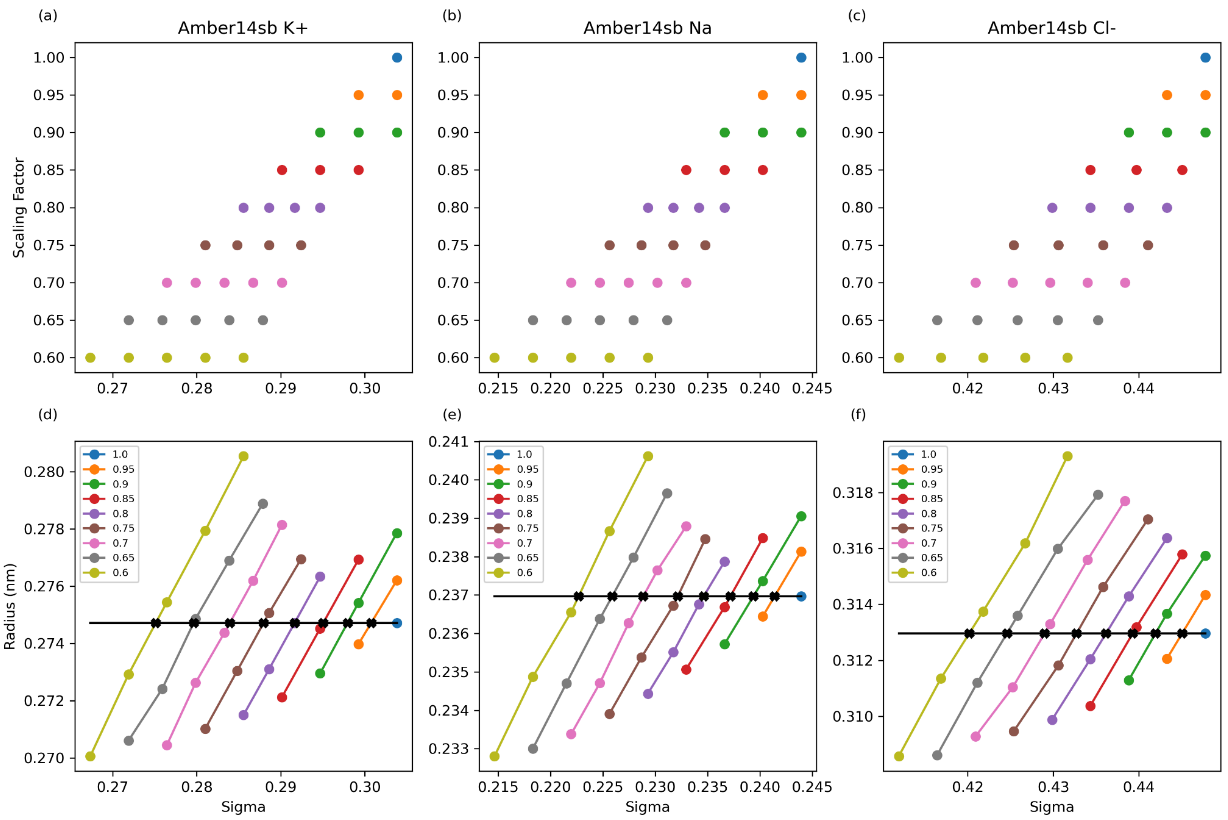

Fig. S9: Ion parameter grid search in Amber14sb. (a)(b)(c) Search grids with (sigma, scaling factor) pairs for $K^+$, $Na^+$, $Cl^-$ ions. (d)(e)(f) The radius of all the points on the grids. The radius from the original radius at scaling factor 1.00 was chosen as the targeted radius. The optimized sigma values for every scaling factor point at a step of 0.05 (1.00, 0.95, 0.90, …) are plotted as x. The sigma parameters for the scaling factor between 0.05 (0.99~0.96, 0.94~0.91, …) are interpolated.

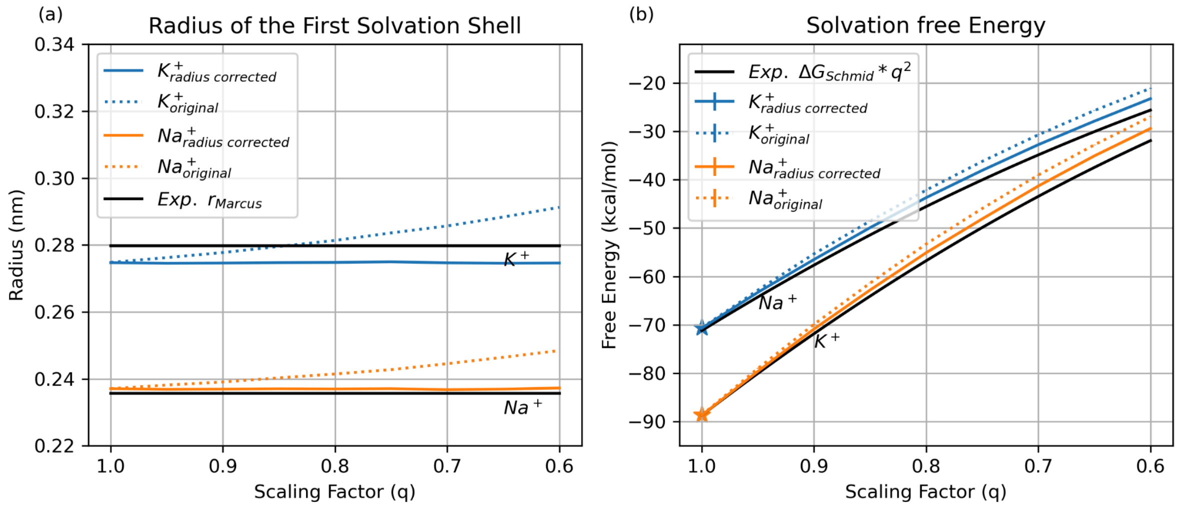

Fig. S10: Comparison of the radius corrected vdW for ECC ion in this work (solid line) and the original vdW (dotted line) in Amber14sb. (a) The radius of the first solvation shell for $K^+$ and $Na^+$ with two sets of vdW parameters. (b) The solvation free energy for $K^+$ and $Na^+$ with two sets of vdW parameters.

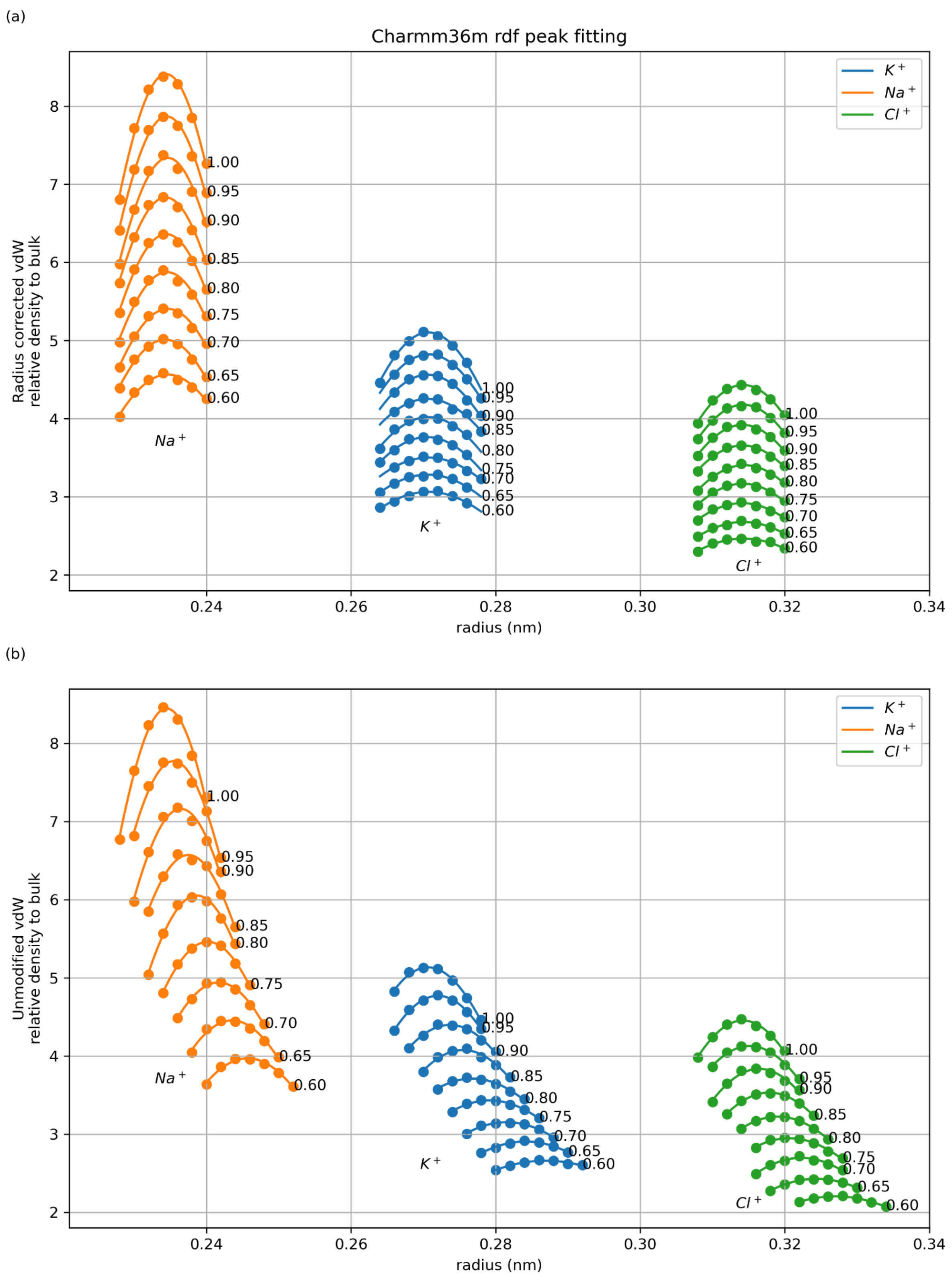

Fig. S11: First solvation shell rdf peak fitting in Charmm36m with radius corrected vdW (a) and unmodified vdW (b).

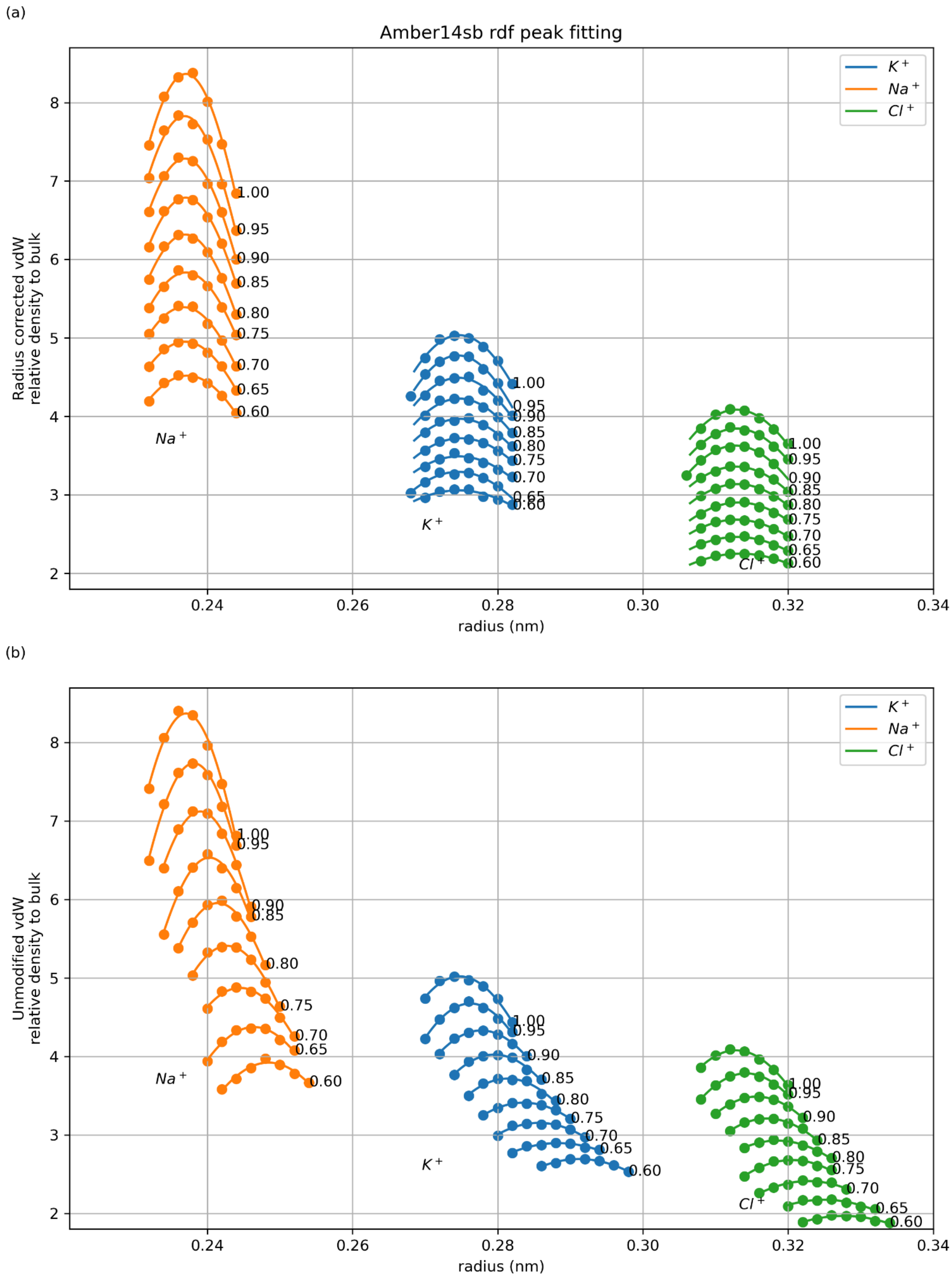

Fig. S12: First solvation shell rdf peak fitting in Amber14sb with radius corrected vdW (a) and unmodified vdW (b).

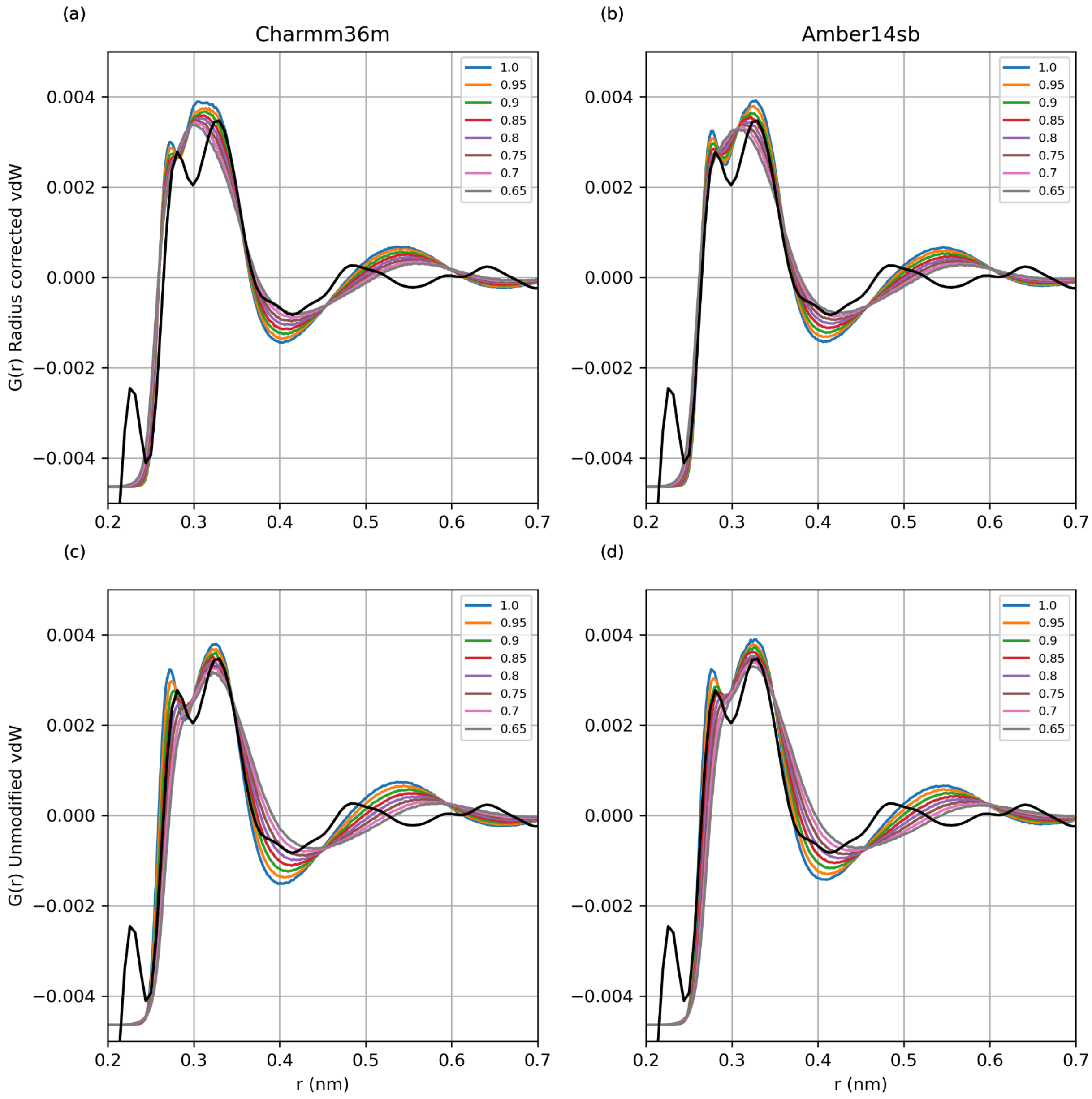

Fig. S13: Comparison of MD simulated RDF with experimental RDF. The experimental RDF is the sum of K-O, K-H(D), K-Cl, and K-K. The simulated RDF is normalized as the experimental publication.[3]

In the parameter sets with radius corrected vdW, the radius of the first peak at 0.28 nm (K-O), and the radius of the second leak at 3.3 nm (K-H) are close to the experimental peaks. The comparison with the experimental RDF cannot distinguish the quality of the parameters with different charges.

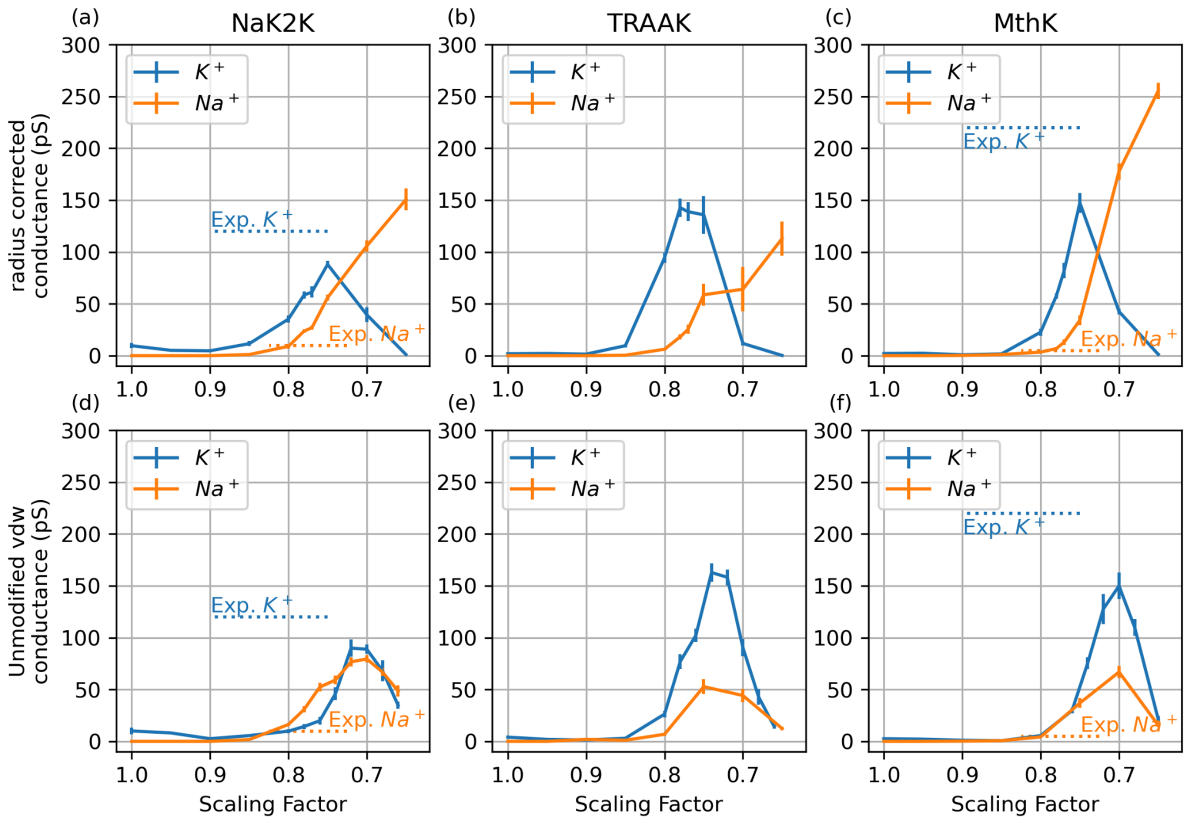

Fig. S14: K+/Na+ conductance with 2 different vdW treatments in Amber14sb. (a-c) Simulated conductance with the vdW that keeps the radii of the ion constant. (d-f) Simulated conductance with the unmodified Amber14sb vdW. All simulations were performed under 150 mV.

| | Charmm36m | | | | | | Amber14sb | | | | | |
|---|---|---|---|---|---|---|---|---|---|---|---|---|
| | K+ | | Na+ | | Cl- | | K+ | | Na+ | | Cl- | |
| charge | sigma(nm) | epsilon (kJ/mol) | sigma(nm) | epsilon (kJ/mol) | sigma(nm) | epsilon (kJ/mol) | sigma(nm) | epsilon (kJ/mol) | sigma(nm) | epsilon (kJ/mol) | sigma(nm) | epsilon (kJ/mol) |
| 1.00 | 0.31426452 | 0.36400800 | 0.25136707 | 0.19622960 | 0.40446802 | 0.62760000 | 0.30379600 | 0.81036900 | 0.24392800 | 0.36584600 | 0.44776600 | 0.14891300 |
| 0.99 | 0.31370893 | | 0.25092105 | | 0.40404701 | | 0.30318805 | | 0.24342284 | | 0.44722335 | |
| 0.98 | 0.31315333 | | 0.25047503 | | 0.40362600 | | 0.30258011 | | 0.24291768 | | 0.44668070 | |
| 0.97 | 0.31259774 | | 0.25002901 | | 0.40320499 | | 0.30197216 | | 0.24241252 | | 0.44613805 | |
| 0.96 | 0.31204214 | | 0.24958299 | | 0.40278398 | | 0.30136422 | | 0.24190736 | | 0.44559540 | |
| 0.95 | 0.31148655 | | 0.24913697 | | 0.40236297 | | 0.30075627 | | 0.24140220 | | 0.44505275 | |
| 0.94 | 0.31087542 | | 0.24878324 | | 0.40189908 | | 0.30019342 | | 0.24099730 | | 0.44443139 | |
| 0.93 | 0.31026430 | | 0.24842952 | | 0.40143519 | | 0.29963056 | | 0.24059239 | | 0.44381004 | |
| 0.92 | 0.30965317 | | 0.24807579 | | 0.40097131 | | 0.29906771 | | 0.24018749 | | 0.44318868 | |
| 0.91 | 0.30904205 | | 0.24772207 | | 0.40050742 | | 0.29850485 | | 0.23978258 | | 0.44256733 | |
| 0.90 | 0.30843092 | | 0.24736834 | | 0.40004353 | | 0.29794200 | | 0.23937768 | | 0.44194597 | |
| 0.89 | 0.30780065 | | 0.24681900 | | 0.39953266 | | 0.29736392 | | 0.23893917 | | 0.44141226 | |
| 0.88 | 0.30717038 | | 0.24626966 | | 0.39902179 | | 0.29678585 | | 0.23850067 | | 0.44087856 | |
| 0.87 | 0.30654011 | | 0.24572031 | | 0.39851092 | | 0.29620777 | | 0.23806216 | | 0.44034485 | |
| 0.86 | 0.30590984 | | 0.24517097 | | 0.39800005 | | 0.29562970 | | 0.23762366 | | 0.43981115 | |
| 0.85 | 0.30527957 | | 0.24462163 | | 0.39748918 | | 0.29505162 | | 0.23718515 | | 0.43927744 | |
| 0.84 | 0.30460951 | | 0.24414149 | | 0.39707871 | | 0.29437078 | | 0.23667342 | | 0.43865405 | |
| 0.83 | 0.30393944 | | 0.24366134 | | 0.39666824 | | 0.29368994 | | 0.23616168 | | 0.43803066 | |
| 0.82 | 0.30326938 | | 0.24318120 | | 0.39625778 | | 0.29300910 | | 0.23564995 | | 0.43740727 | |
| 0.81 | 0.30259931 | | 0.24270105 | | 0.39584731 | | 0.29232826 | | 0.23513821 | | 0.43678388 | |
| 0.80 | 0.30192925 | | 0.24222091 | | 0.39543684 | | 0.29164742 | | 0.23462648 | | 0.43616049 | |
| 0.79 | 0.30112750 | | 0.24171205 | | 0.39478710 | | 0.29091019 | | 0.23413197 | | 0.43547221 | |
| 0.78 | 0.30032574 | | 0.24120320 | | 0.39413736 | | 0.29017297 | | 0.23363746 | | 0.43478392 | |
| 0.77 | 0.29952399 | | 0.24069434 | | 0.39348762 | | 0.28943574 | | 0.23314294 | | 0.43409564 | |
| 0.76 | 0.29872223 | | 0.24018549 | | 0.39283788 | | 0.28869852 | | 0.23264843 | | 0.43340735 | |
| 0.75 | 0.29792048 | | 0.23967663 | | 0.39218814 | | 0.28796129 | | 0.23215392 | | 0.43271907 | |
| 0.74 | 0.29716759 | | 0.23914351 | | 0.39157835 | | 0.28715762 | | 0.23149487 | | 0.43197204 | |
| 0.73 | 0.29641469 | | 0.23861039 | | 0.39096856 | | 0.28635394 | | 0.23083582 | | 0.43122501 | |
| 0.72 | 0.29566180 | | 0.23807726 | | 0.39035877 | | 0.28555027 | | 0.23017676 | | 0.43047799 | |
| 0.71 | 0.29490890 | | 0.23754414 | | 0.38974898 | | 0.28474659 | | 0.22951771 | | 0.42973096 | |
| 0.70 | 0.29415601 | | 0.23701102 | | 0.38913919 | | 0.28394292 | | 0.22885866 | | 0.42898393 | |
| 0.69 | 0.29326681 | | 0.23625808 | | 0.38847510 | | 0.28308173 | | 0.22826536 | | 0.42810378 | |
| 0.68 | 0.29237760 | | 0.23550514 | | 0.38781101 | | 0.28222053 | | 0.22767205 | | 0.42722363 | |
| 0.67 | 0.29148840 | | 0.23475221 | | 0.38714693 | | 0.28135934 | | 0.22707875 | | 0.42634349 | |

| | | | | | | | | | | | |
|---|---|---|---|---|---|---|---|---|---|---|---|
| 0.66 | 0.29059919 | | 0.23399927 | | 0.38648284 | | 0.28049814 | | 0.22648544 | | 0.42546334 |
| 0.65 | 0.28970999 | | 0.23324633 | | 0.38581875 | | 0.27963695 | | 0.22589214 | | 0.42458319 |
| 0.64 | 0.28876672 | | 0.23259615 | | 0.38504512 | | 0.27873764 | | 0.22525181 | | 0.42370099 |
| 0.63 | 0.28782345 | | 0.23194596 | | 0.38427148 | | 0.27783834 | | 0.22461149 | | 0.42281879 |
| 0.62 | 0.28688017 | | 0.23129578 | | 0.38349785 | | 0.27693903 | | 0.22397116 | | 0.42193659 |
| 0.61 | 0.28593690 | | 0.23064559 | | 0.38272421 | | 0.27603973 | | 0.22333084 | | 0.42105439 |
| 0.60 | 0.28499363 | | 0.22999541 | | 0.38195058 | | 0.27514042 | | 0.22269051 | | 0.42017219 |

Table S5: The radius corrected vdW parameters for ions at different scaling factors in Charmm36m and Amber14sb.

### 4. Permeation definitions

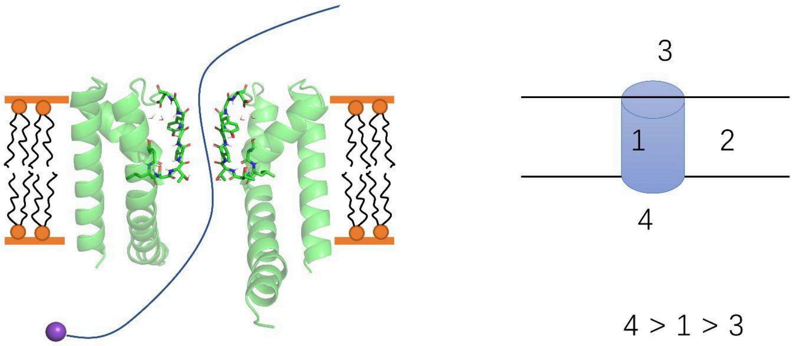

Fig. S15: Illustration of permeation counting.

For counting of permeation events and selectivity filter states a custom python script was used, the counting procedure and SF site definitions are briefly explained below.
The trajectory was first centered on the protein. The simulation box was separated into 4 compartments: (1) the selectivity filter, (2) the middle compartment, (3) the upper compartment, (4) the lower compartment.
(1) The selectivity filter compartment was defined as the cylinder which centered on selectivity filter oxygens. The top z limit was defined as the backbone oxygen on top of S1. The bottom z limit was defined as the Threonine hydroxyl oxygen. The radius of the cylinder was set as 3.5 Å.
(2) The middle compartment was defined as the space with the z coordinate as compartment (1) but outside the cylinder.
(3) The upper compartment was defined as the space above the compartment (1)
(4) The lower compartment was defined.
An ion up-permeation event was defined as an ion being in 4, 1, 3 compartment consecutively. An ion down-permeation event was defined as an ion being in 3, 1, 4 compartment consecutively.

### 5. Binding site definitions

All binding sites were defined as the cylinder region center by SF oxygens. The z position was defined as such (we take NaK2K as an example which has the SF sequence of TVGYG):

S0 : z coordinates below the carbonyl oxygen of Gly67, and above the carbonyl oxygen of Tyr66.
S1 : z coordinates below the carbonyl oxygen of Tyr66, and above the carbonyl oxygen of Gly65.
S2 : z coordinates below the carbonyl oxygen of Gly65, and above the carbonyl oxygen of Val64.
S3 : z coordinates below the carbonyl oxygen of Val64, and above the carbonyl oxygen of Thr63.
S4 : z coordinates below the carbonyl oxygen of Thr63, and above the hydroxyl oxygen of Thr63.
S5 : z coordinate below the hydroxyl oxygen of Thr63, until 1.8 Å below the hydroxyl oxygen of Thr63.

The diameter of S0 was defined as the distance between the carbonyl oxygen of Gly67 in the opposite chain. The radius of S1-S4 was defined as 3.5 Å. The radius of S5 was defined as 9 Å.

### 6. Protein stability analysis across different scaling factors

#### 6.1. Principal component analysis (PCA) of SF

PCA is applied to examine if there are potential artifacts in the charge scaled force fields. Multiple trajectories simulated at different scaling factors (1.00, 0.95, 0.90 … , 0.65) are concatenated together as one data set for PCA. The same analysis is applied for SF in this section, and protein backbone in 4.2, in NaK2K, TRAAK, and MthK.

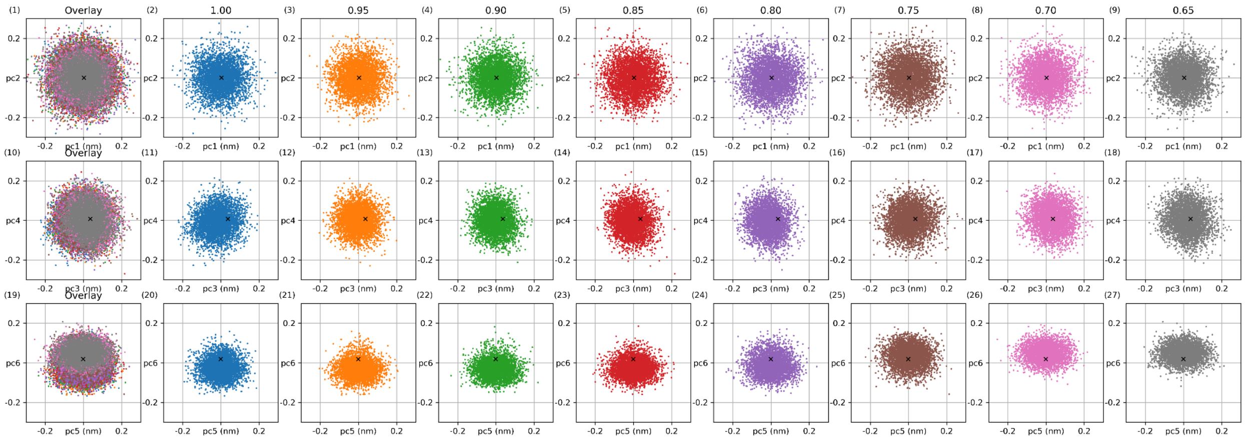

Fig. S15.1: SF PCA in NaK2K with Charmm36m in different scaling factors. PCA is applied to the expanded ensemble trajectory obtained from Hamiltonian replica exchange simulations with different scaling factors. Each column (except the first) corresponds to a different scaling factor, while the first column overlays all trajectories. Each row represents a projection onto different principal components (PCs): PC1 (x)–PC2 (y) in the first row, PC3 (x)–PC4 (y) in the second row, and PC5 (x)–PC6 (y) in the third row. The X-ray structure is marked as x in each plot.

In PC1–PC5, no significant differences are observed across scaling factors. In PC6, different scaling factors sample different projected values. The motion corresponding to PC6 is shown in Fig. S14.2. At a lower scaling factor (0.70, 0.65), the SF is elongated along the Z-axis, and the distance between the VAL oxygen (between S2 and S3) decreases. This small motion is reasonable given the increase in ion occupancy from 3 to 4 (Fig. 2(g)).

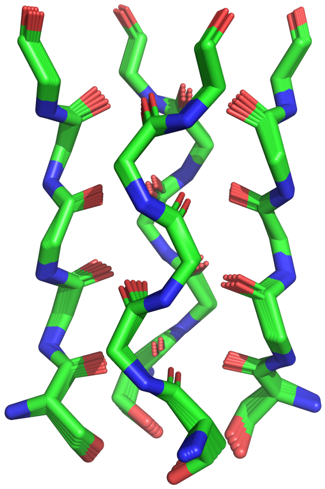

Fig. S15.2: NaK2K SF motion in PC6 in Charmm36m.

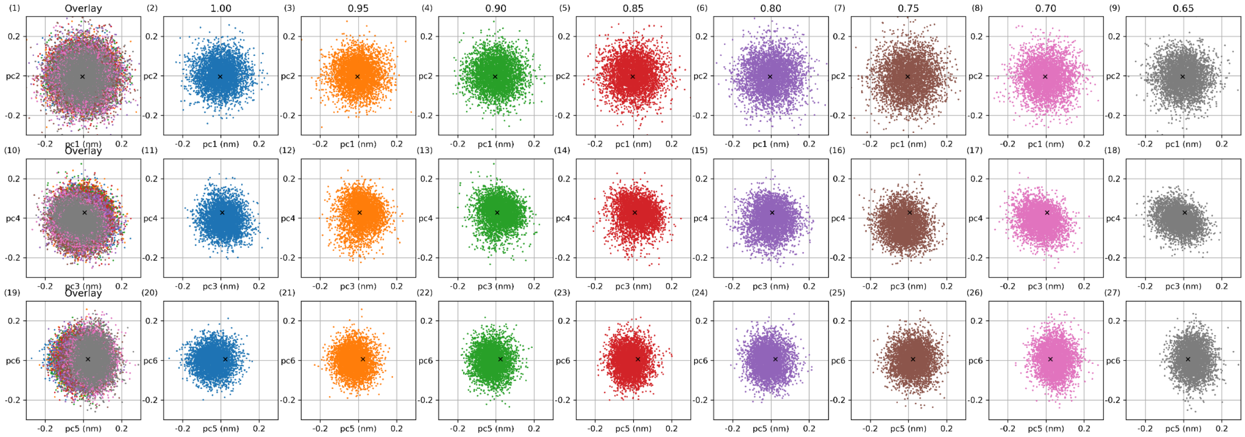

Fig. S15.3: SF PCA in NaK2K with Amber14sb-S3 in different scaling factors. PCA is applied to the expanded ensemble trajectory obtained from Hamiltonian replica exchange simulations with different scaling factors. Each column (except the first) corresponds to a different scaling factor, while the first column overlays all trajectories. Each row represents a projection onto different principal components (PCs): PC1 (x)–PC2 (y) in the first row, PC3 (x)–PC4 (y) in the second row, and PC5 (x)–PC6 (y) in the third row. The X-ray structure is marked as x in each plot.

In PC1–PC4, and PC6, no significant differences are observed across scaling factors in NaK2K with Amber14sb-S3 (Fig. S14.3). The motion of PC5 is shown in Fig. S14.4. Comparing 2 force fields (Charmm36m and Amber14sb-S3), PC6 in Charmm36m is very similar to PC5 in Amber14sb-S3, both represent the elongation along the Z-axis and the decrease of the radius in the middle of the SF with the increase of occupancy from 3 to 4 (Fig. 2(i)).

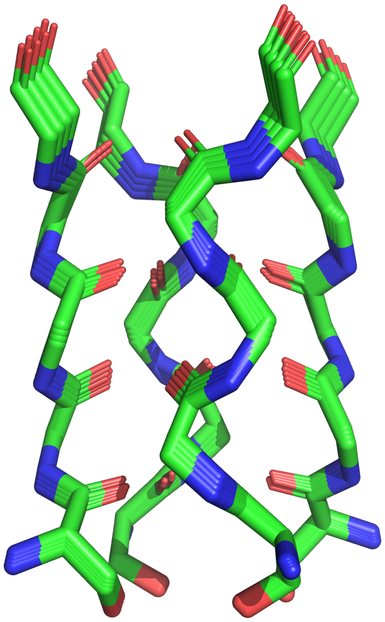

Fig. S15.4: NaK2K SF motion in PC5 in Amber14sb-S3.

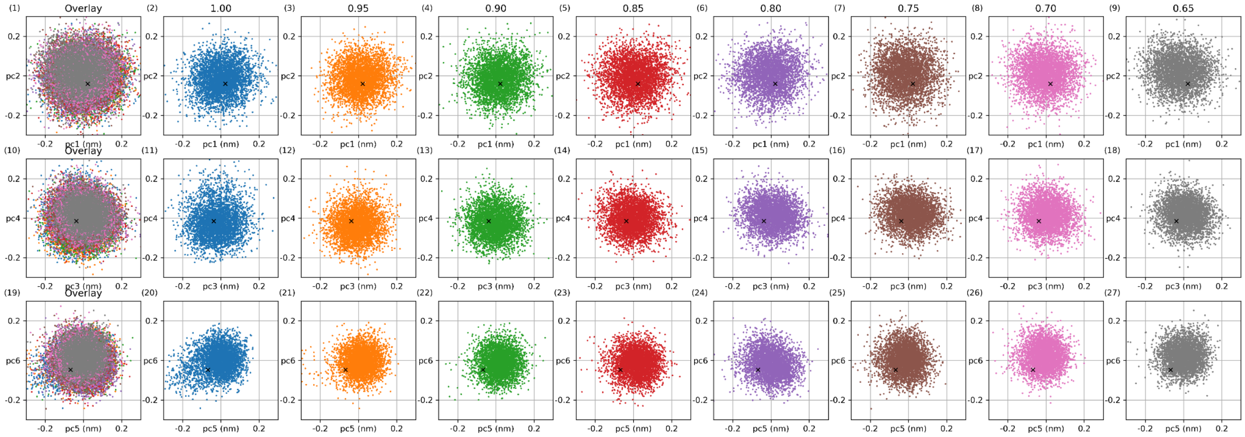

Fig. S16.1: SF PCA in TRAAK with Charmm36m in different scaling factors. PCA is applied to the expanded ensemble trajectory obtained from Hamiltonian replica exchange simulations with different scaling factors. Each column (except the first) corresponds to a different scaling factor, while the first column overlays all trajectories. Each row represents a projection onto different principal components (PCs): PC1 (x)–PC2 (y) in the first row, PC3 (x)–PC4 (y) in the second row, and PC5 (x)–PC6 (y) in the third row. The X-ray structure is marked as x in each plot.

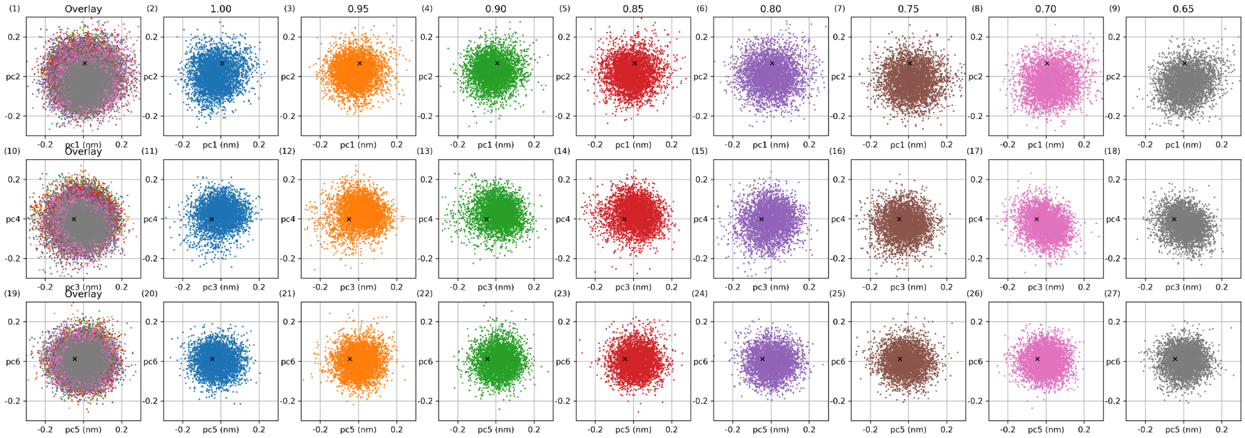

Fig. S16.2: SF PCA in TRAAK with Amber14sb-S3 in different scaling factors. PCA is applied to the expanded ensemble trajectory obtained from Hamiltonian replica exchange simulations with different scaling factors. Each column (except the first) corresponds to a different scaling factor, while the first column overlays all trajectories. Each row represents a projection onto different principal components (PCs): PC1 (x)–PC2 (y) in the first row, PC3 (x)–PC4 (y) in the second row, and PC5 (x)–PC6 (y) in the third row. The X-ray structure is marked as x in each plot.

The same PCA is applied to SF of TRAAK simulation in Charmm36m and Amber14sb-S3 (Fig. S15). Different scaling factors sample similar values on PC1 to PC6.

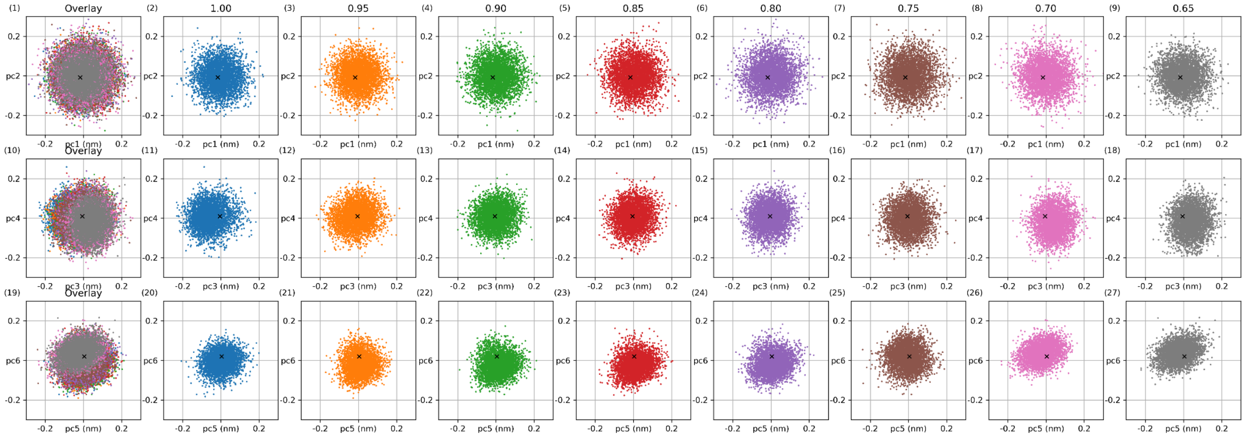

Fig. S17.1: SF PCA in MthK with Charmm36m in different scaling factors. PCA is applied to the expanded ensemble trajectory obtained from Hamiltonian replica exchange simulations with different scaling factors. Each column (except the first) corresponds to a different scaling factor, while the first column overlays all trajectories. Each row represents a projection onto different principal components (PCs): PC1 (x)–PC2 (y) in the first row, PC3 (x)–PC4 (y) in the second row, and PC5 (x)–PC6 (y) in the third row. The X-ray structure is marked as x in each plot.

In the PCA of SF trajectory in MthK with Charmm36m. Different scaling factors give different projected values in PC6, and the motion of PC6 is shown in Fig. S16.2. The motion is highly similar to what is explained in PCA of SF in NaK2K.

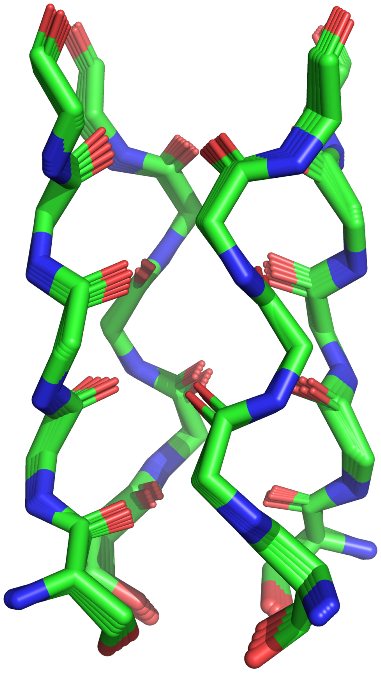

Fig. S17.2: MthK SF motion in PC6 in Charmm36m.

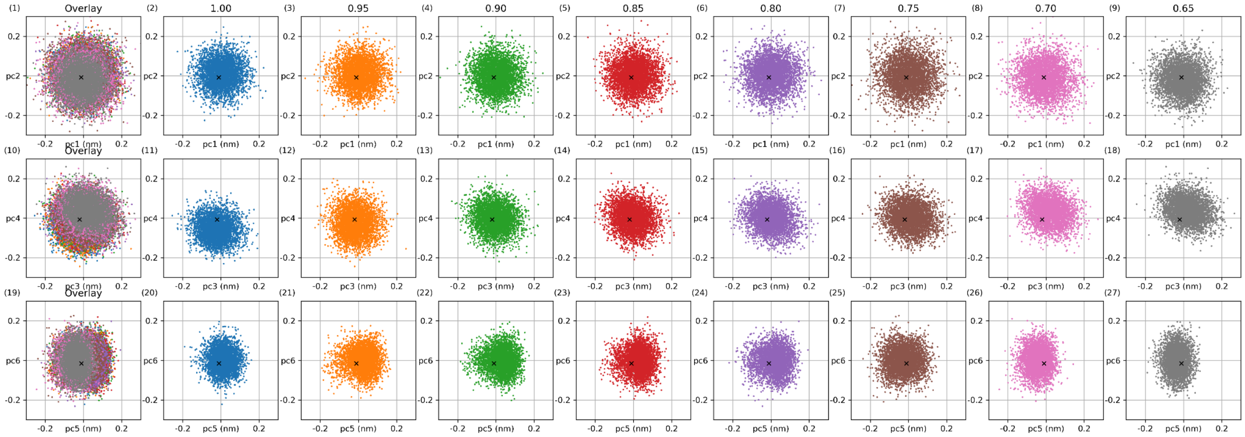

Fig. S17.3: SF PCA in MthK with Amber14sb-S3 in different scaling factors. PCA is applied to the expanded ensemble trajectory obtained from Hamiltonian replica exchange simulations with different scaling factors. Each column (except the first) corresponds to a different scaling factor, while the first column overlays all trajectories. Each row represents a projection onto different principal components (PCs): PC1 (x)–PC2 (y) in the first row, PC3 (x)–PC4 (y) in the second row, and PC5 (x)–PC6 (y) in the third row. The X-ray structure is marked as x in each plot.

In the PCA of SF trajectory in MthK with Amber14sb-S3. Different scaling factors give different projected values in PC5, and the motion of PC5 is shown in Fig. S16.2. The motion is highly similar to PC6 in the PCA of SF trajectory in MthK with Charmm36m and similar to what is explained in PCA of SF in NaK2K.

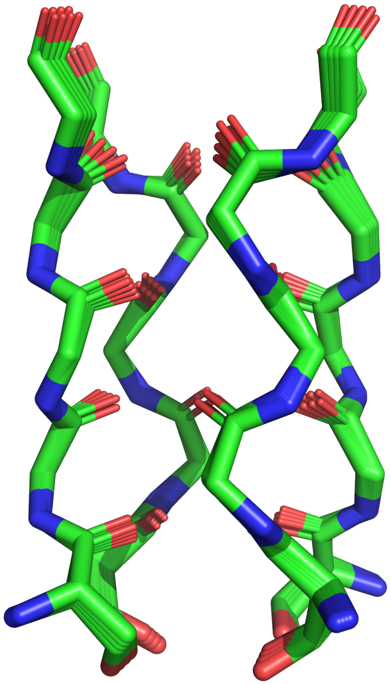

Fig. S17.4: MthK SF motion in PC5 in Amber14sb-S3.

##### 6.2. PCA of backbone

PCA is also applied to the expanded ensemble trajectory with different scaling factors to check the overall structure of protein. The backbone trajectories from the Hamiltonian replica exchange simulation are concatenated together for PCA. Each trajectories are later projected on PCs and visualized in Fig. S17 to Fig. S19. No significant difference can be found in the distribution of the projected value in different scaling factors.

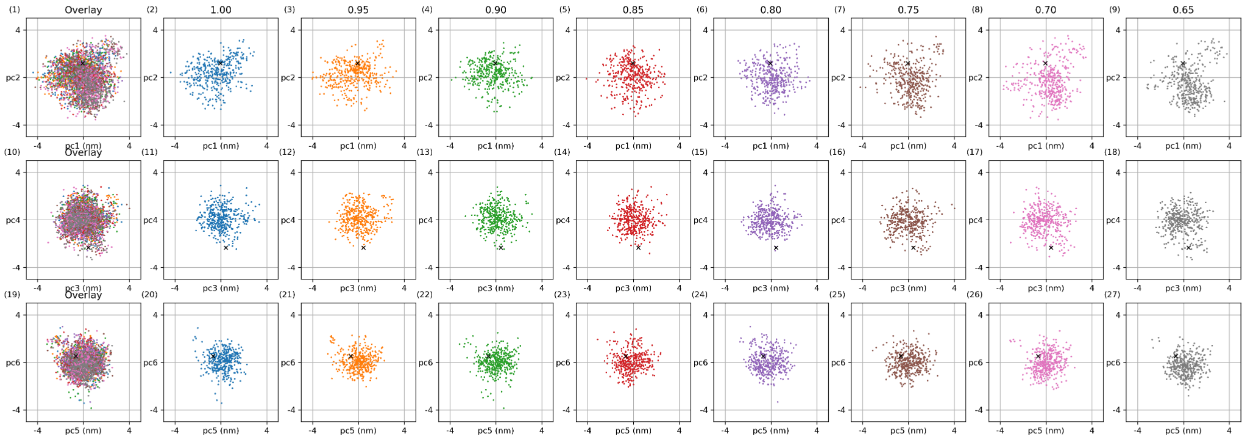

Fig. S18.1: Backbone PCA in NaK2K with Charmm36m in different scaling factors. PCA is applied to the expanded ensemble trajectory obtained from Hamiltonian replica exchange simulations with different scaling factors. Each column (except the first) corresponds to a different scaling factor, while the first column overlays all trajectories. Each row represents a projection onto different principal components (PCs)

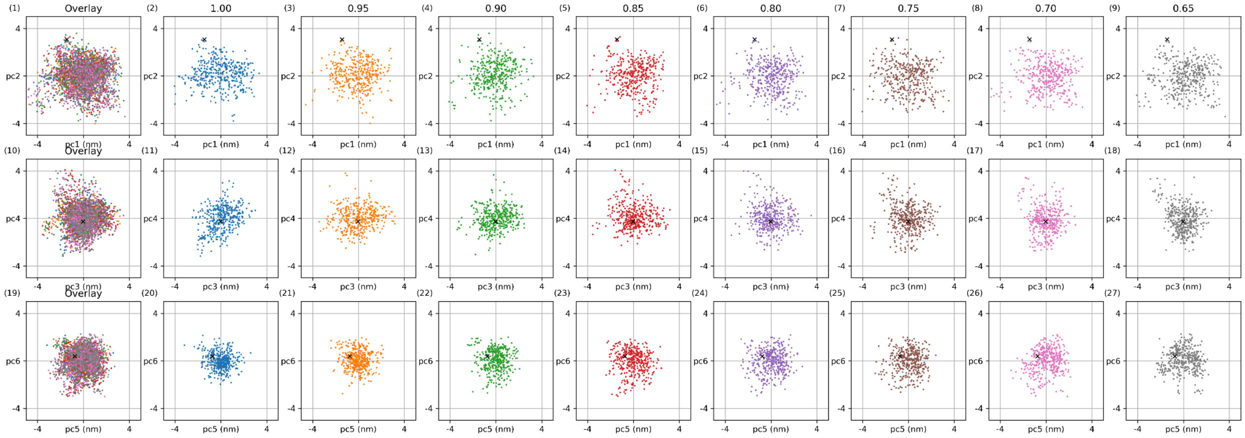

Fig. S18.2: Backbone PCA in NaK2K with Amber14sb-S3 in different scaling factors. PCA is applied to the expanded ensemble trajectory obtained from Hamiltonian replica exchange simulations with different scaling factors. Each column (except the first) corresponds to a different scaling factor, while the first column overlays all trajectories. Each row represents a projection onto different principal components (PCs)

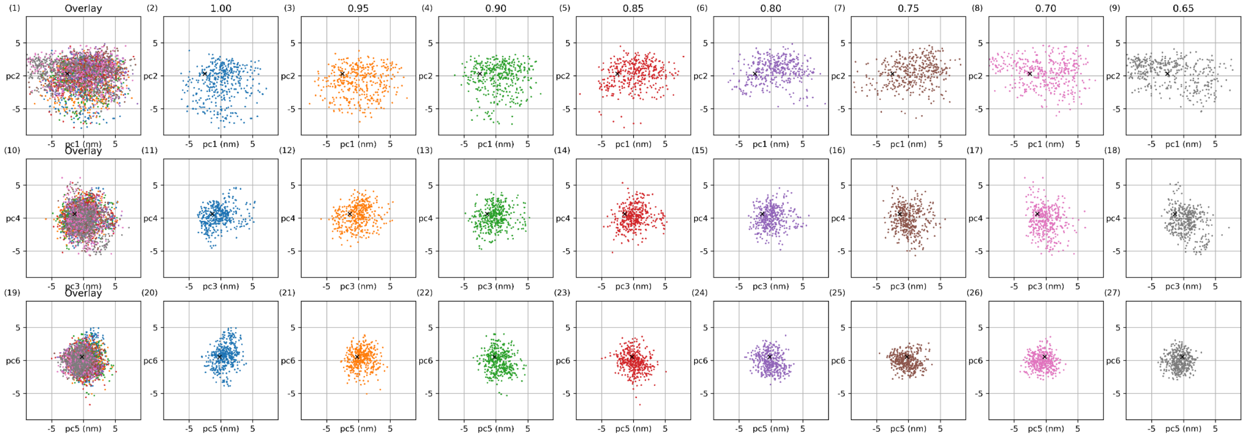

Fig. S19.1: Backbone PCA in TRAAK with Charmm36m in different scaling factors. PCA is applied to the expanded ensemble trajectory obtained from Hamiltonian replica exchange simulations with different scaling factors. Each column (except the first) corresponds to a different scaling factor, while the first column overlays all trajectories. Each row represents a projection onto different principal components (PCs)

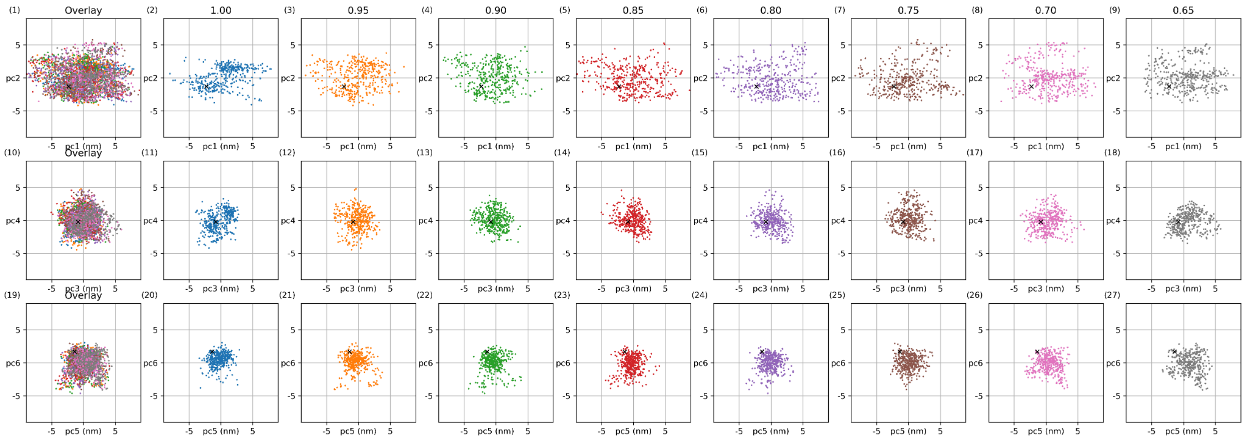

Fig. S19.2: Backbone PCA in TRAAK with Amber14sb-S3 in different scaling factors. PCA is applied to the expanded ensemble trajectory obtained from Hamiltonian replica exchange simulations with different scaling factors. Each column (except the first) corresponds to a different scaling factor, while the first column overlays all trajectories. Each row represents a projection onto different principal components (PCs)

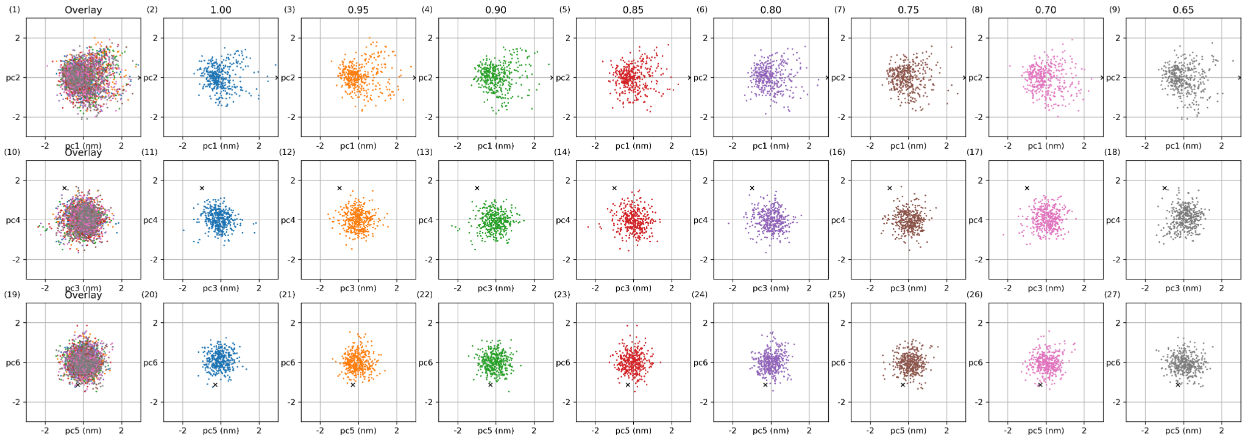

Fig. S20.1: Backbone PCA in MthK with Charmm36m in different scaling factors. PCA is applied to the expanded ensemble trajectory obtained from Hamiltonian replica exchange simulations with different scaling factors. Each column (except the first) corresponds to a different scaling factor, while the first column overlays all trajectories. Each row represents a projection onto different principal components (PCs)

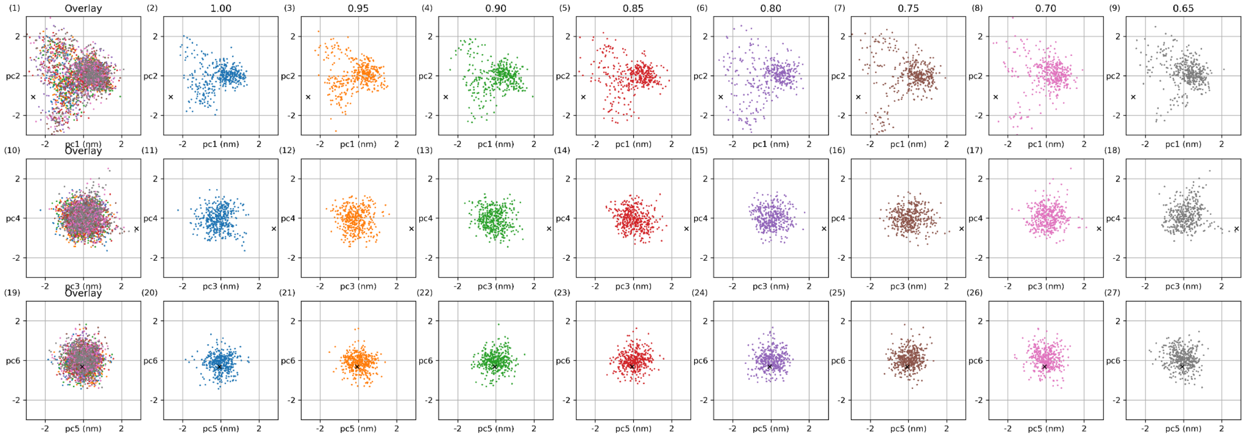

Fig. S20.2: Backbone PCA in MthK with Amber14sb-S3 in different scaling factors. PCA is applied to the expanded ensemble trajectory obtained from Hamiltonian replica exchange simulations with different scaling factors. Each column (except the first) corresponds to a different scaling factor, while the first column overlays all trajectories. Each row represents a projection onto different principal components (PCs)

### 7. Amino acid charge scaling example

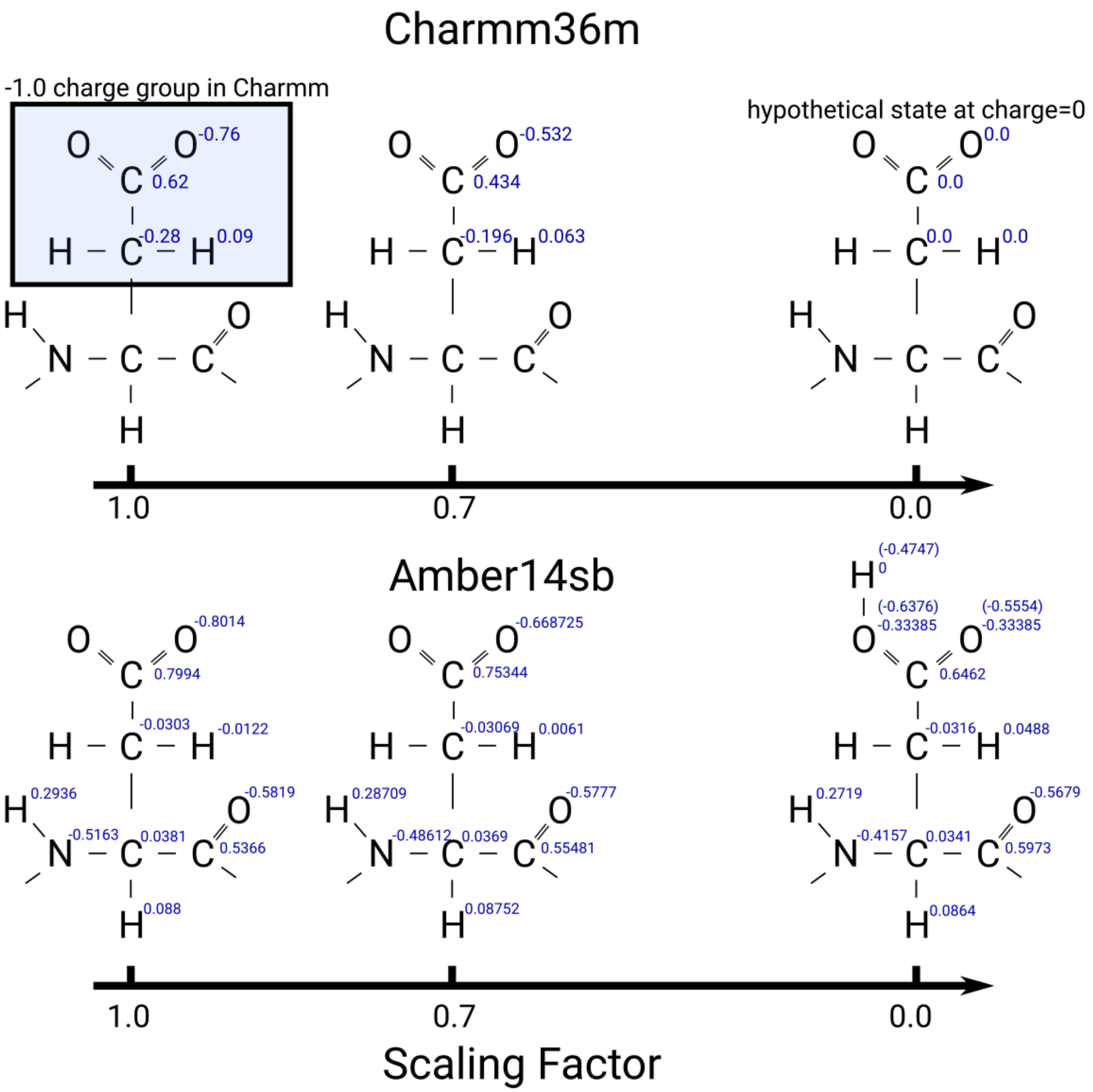

Fig. S21: Amino Acid charge scaling in 2 force fields with ASP as an example. In Charmm36m, every atom in the +-1.0 charge group is scaled with the scaling factor. In Amber14sb, the protonated ASP is taken as the hypothetical 0.0 charge ASP, and the charge is linearly interpolated between 1.0 and 0.0.

### 8. Code availability

All analysis code in this work is available on the github repository.
https://github.com/deGrootLab/Sfilter_Cylinder

All force field, starting structures, and parameter files for reproducing this paper are provided on the github repository.
https://github.com/deGrootLab/Charge_Scaling_in_Potassium_Channel_Simulations_paper
